# Rhizosphere bacterial colonization of beet occurs in discrete phases regardless of bioinoculation with the wild sea beet root community

**DOI:** 10.1101/2023.02.10.527839

**Authors:** Marcin Gołębiewski, Marcin Sikora, Justyna Mazur, Sonia Szymańska, Jarosław Tyburski, Katarzyna Hrynkiewicz, Werner Ulrich

## Abstract

Bioinoculation can increase crop yields under environmental stress. Plant colonization by microbes is an example of succession, with its distinct phases differing in community structure and diversity. This process needs to be studied to determine the optimal timing for bioinoculation and its effects. Haere, we show that, regardless of bio-inoculation, soil type and plant genotype, bacteria colonize the rhizosphere of axenic beets and tissues in two phases, differing in bacterial load, nestedness, community structure, diversity and assembly mechanism, and associated with taproot development. Communities remained stable after five weeks of growth in soil. The alpha diversity was greater and the bacterial load was lower in the late samples than in the early ones. Time, soil type and genotype determined community structure but not alpha diversity, bacterial load, nestedness or assembly mechanisms both in the rhizosphere and in the endosphere. Inoculation changed the community structure and members of Pseudomonadota and Bacillota of low abundance in the inoculant were recruited by beets.

Axenic beet colonization occurs through phases similar to other instances of microbial succession, and bacteria are recruited mostly randomly. The transition from the early to late phase involves a decrease in the bacterial load in plant tissues, which may be linked to plant growth and the arrest of bacterial cell division. Therefore, early inoculation seems to be favourable. Five weeks of growth in soil enabled formation of stable bacterial communities in both the rhizosphere and the endosphere. The influence of inoculation seems to be indirect, probably due to microbe-microbe interactions.

## Background

Bioinoculation, defined as the introduction of beneficial endophytic microbes into plant tissues, effectively enhances nutrient access and increases resistance to pathogens, leading to higher crop yields, especially in plants grown under environmental stress [1–3]. The construction of a successful bioinoculant (also known as biofertilizer) requires addressing three key challenges: i) identifying suitable microorganism(s), ii) devising an effective formulation and iii) determining the optimal method for its application. In terms of the first challenge, bioinoculants consisting of microbial communities rather than single strains offer a potential solution that ensures broader applicability, allowing for the use of different microbes tailored to the specific needs of various plant genotypes, such as strains and cultivars, in diverse soil conditions. For instance, the bioinoculation of *Arabidopsis thaliana* with a synthetic microbial community has been shown to restore growth under stressful conditions [4]. For the formulation challenge, various strategies involving liquids, slurries, or solids have been reviewed in the literature [5, 6], and bioinoculants are applied as seed coats [7], mixed with soil [8] or used as foliar sprays [9]. Our previous research revealed that beet genotype influences both endophytic and rhizosphere bacterial communities, and lyophilized beet roots can serve as a source of viable microorganisms for inoculation [10].

From an ecological standpoint, inoculation influences the structure of endophytic microbial communities, potentially altering the process of community assembly [11, 12]. Research on the dynamics of plant microbial colonization has demonstrated that this process occurs rapidly [13, 14] and is driven by selection caused by interactions with a host plant [15, 16] as well as by microbe‒ microbe interactions [17]. Consequently, the host genotype is one of the key factors influencing endophytic community structure [18], meaning that different varieties of the same plant species may exhibit varying responses to a given bioinoculant. Although soil has been identified as the primary source of endophytes, seeds have been found to be more important in certain cases [19, 20], and the phyllosphere can also serve as an additional source [21]. Therefore, the success or failure of a bioinoculant may also be influenced by soil conditions.

Endophytic communities evolve over time [22–24] and respond to changes in host developmental stages [25, 26] and to environmental conditions [27]. In fact, in the case of axenic plants, their colonization might be considered an instance of primary microbial succession. Microbial primary succession has shown similarities to plant succession in different environments and follows similar phases [27]. However, plant colonization differs from other instances of succession due to the additional layer of complexity introduced by plants, including disturbances and plant development, which to some extent govern this process.

Many ecological sets of communities were found to be nested; that is, species-poor communities are proper subsets of species-rich communities. Nestedness analysis is therefore a common tool for disentangling richness and structural effects on changes in community composition [28]. However, nestedness alone does not convey information on the processes governing the assembly [29], and other tools, such as βNTI or βNRI [30] coupled with Raup–Crick dissimilarity based on the Bray–Curtis index [31], are needed to paint a full picture of community assembly.

In studies on bioinoculation, tracing the entire lifespan of a plant is often impractical, particularly for large or perennial species. Moreover, the application and assessment of bioinoculants require determining the optimal timing for their use [32]. Consequently, questions arise regarding whether and, if so, when endophytic communities reach compositional stability and when to examine the influence of inoculants on host plants. However, it may be convenient to analyze the earliest stages, such as the seedling stage, for practical reasons, but this might not be the best approach if colonization requires more time. Additionally, it is interesting to investigate how bioinoculants influence rhizosphere and endophytic communities and whether this influence is dependent on soil type and plant genotype. Such data could be valuable in engineering novel bioinoculants.

Beet is an important crop and exceptional plant – one of those whose undomesticated ancestor still grows in the wild. The ancestor, sea beet, is genetically very similar to domesticated beets but strongly differs in ecology [33, 34]. Beets are cultivated for their high sucrose content (sugar beet), as a root (red beet) or leafy (chard) vegetable or as fodder. According to recent FAO data, approximately 30% of the world’s sucrose production comes from sugar beet [35]. The beet taproot forms early in plant development, and its growth can be divided into three phases differing physiologically and biochemically: pre-storage, transition/secondary growth and sucrose accumulation [36]. These phases presumably also differ in terms of the quantity and quality of root exudates [37, 38].

The sugar beet microbiome has been extensively studied (reviewed in [39]), and many bioinoculants have been proposed (e.g., [40, 41]), while data on the wild beet microbiome are scarce. The beet microbiome was studied either at very early stages of plant development [42] or, if a study involved the whole plant lifespan, was based on very short reads and a limited number of replicates [43]. To bridge these gaps, we decided to include sea beet in our study and address three questions: i) to determine the time required for the establishment of stable endophytic communities in beet plants—given that the analysed growth period encompassed all three phases of root development—we hypothesized that stability would be achieved within six weeks after planting in all genotypes; ii) to assess whether community assembly and the degree of nestedness vary over time and among different genotypes; and iii) to examine the extent and manner in which inoculation with lyophilized wild beet roots influences bacterial communities in the rhizosphere soils, as well as in bacteria-free sugar beet and sea beet plants. To answer these questions, we analyzed bacterial communities through 16S rRNA gene fragment sequencing at five time points in the rhizosphere, roots, and leaves of three beet genotypes cultivated in two soils with contrasting edaphic properties, with or without inoculation using lyophilized sea beet roots.

## Materials and Methods

### Experimental design

We cultivated three beet varieties in two different soils, where each combination of variety (genotype) and soil variant was inoculated with either the native microbiome of *B. vulgaris* ssp. *maritima* sourced from the wild or left non-inoculated. Our analysis included the examination of soils, roots, and leaves at five specific time points: T0, four weeks after planting in the soil and immediately prior to inoculation; T1, twenty-nine days after planting (one day post-inoculation); T2, thirty-five days after planting (seven days post-inoculation); T3, fifty-six days after planting (28 days post-inoculation); and T4, eighty-six days after planting (58 days post-inoculation). At the beginning of the experiment, twenty-five plants were grown in a 14 cm (h) × 15 cm (w) × 37 cm (l) pot filled with approximately 7 litres of soil. During each sampling step, we carefully removed five plants from each pot. A single plant was considered a technical replicate, and five plants from the same pot were considered biological replicates. We employed five biological replicates for each experimental variant. A schematic representation of the experimental design is depicted in Fig. 1.

**Fig. 1.**
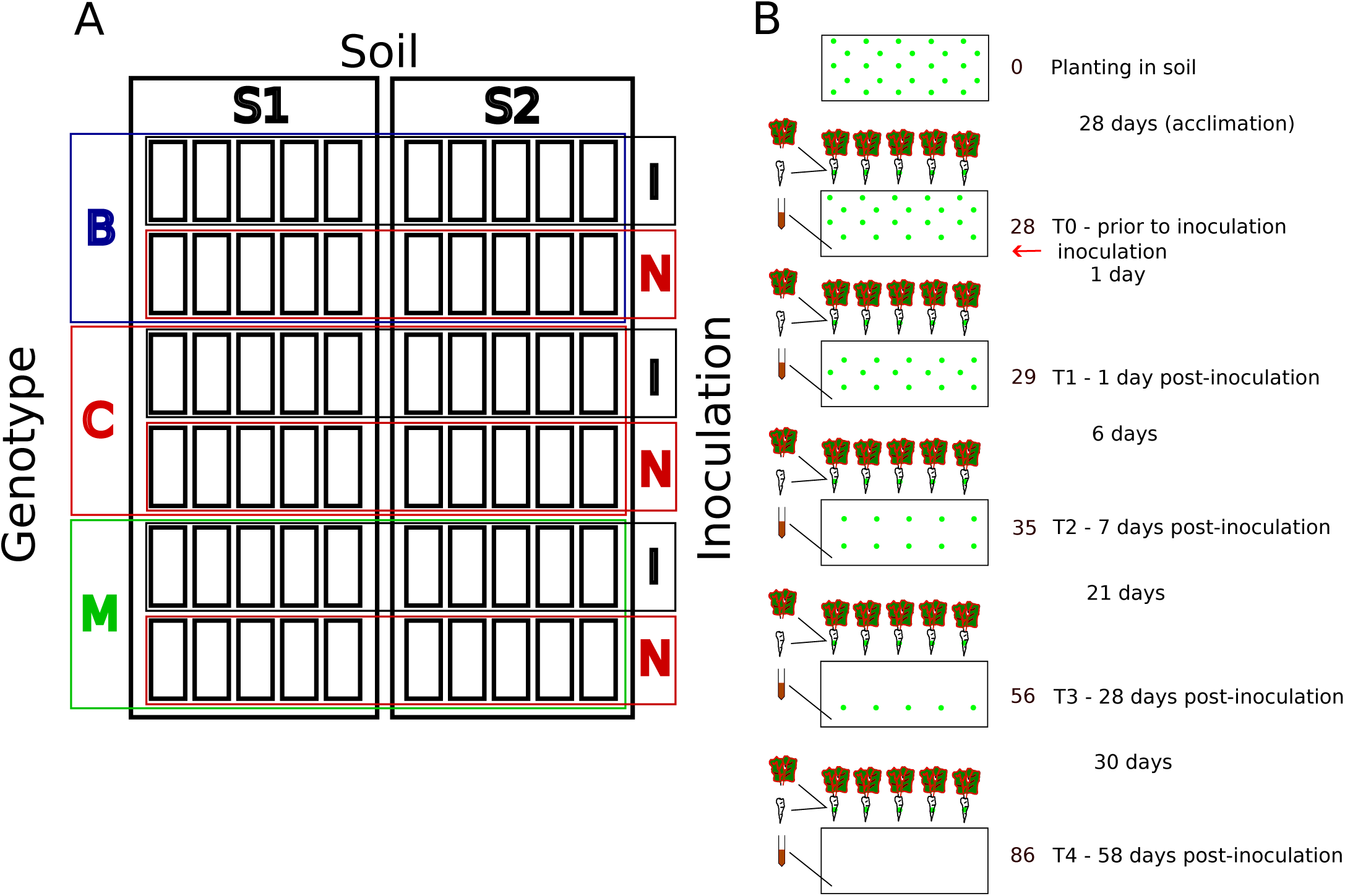
Experimental design. A) Experimental plots; there were two soils, S1 and S2, and three genotypes (B – *Beta vulgaris* ssp. *vulgaris* cv ‘Bravo’; C – B*. vulgaris* ssp. *vulgaris* cv. ‘Cassino’, M – *B. vulgaris* ssp. *maritima*), which were inoculated with either lyophilized roots of sea beets growing in the wild (I) or the noninoculated control (N). Each timepoint (soil × genotype × inoculation) consisted of five biological replicates (BRs-pots), and there were five technical replicates (TRs-plants) for each BR. B) A single experimental variant. Five technical replicates were collected at each timepoint from a given pot (BR). Leaves and soils were directly snap-frozen in liquid nitrogen, and roots were surface-sterilized prior to freezing.

### Soils

We selected two soils obtained from commercial sugar beet plantations. The beans were cultivated in the sampled fields for at least three successive years prior to sampling according to standard agricultural practices recommended by sugar-producing companies. Soils (0-40 cm depth) were sampled during the fall of 2017 before harvest. The samples were stored in sealed plastic bags at ∼15°C until use. The specific physicochemical characteristics of the soils are provided in Table SR1. To minimize the impact of their native microbiomes on the plants, the soils underwent a pasteurization process. This involved subjecting them to two rounds of autoclaving, each lasting for one hour at a temperature of 121°C, over a week.

### Plants

In our study, we utilized two varieties of sugar beet, namely, *Beta vulgaris* ssp. *vulgaris* cv.’Bravo’ and cv.’Casino,’ along with sea beet (*Beta vulgaris* ssp. *maritima*). The plant material was obtained from two different sources. The sugar beet cultivars (KHBC, Kutno, Poland) were obtained from commercial seeds, while the seeds of sea beet were obtained from the National Germplasm Resources Laboratory in Beltsville, MD. Prior to use, the seeds (with the coating removed) were surface-sterilized. The efficiency of sterilization was evaluated through plating, as detailed in Supplementary Methods SM1.

After germination, the seedlings were carefully dissected and subjected to micropropagation in the presence of cefotaxime and vancomycin. The resulting plantlets were further subjected to three rounds of micropropagation on antibiotic-free media. Subsequently, the explants were rooted and acclimated to *ex vitro* conditions (for further details, refer to Supplementary Method SM2) before being planted in their respective soils.

### Inoculant preparation

In August 2017, sea beet plants growing along the northern Adriatic coast of Croatia were collected for our research purposes. Since the collection was conducted in non-protected areas of public grounds and solely for scientific investigation, specific permission was not required in compliance with the Nagoya Protocol and applicable EU, Croatian, and Polish laws. The collected plants were promptly refrigerated in styrofoam boxes containing cooling pads that had been frozen at-80°C prior to the sampling campaign. This procedure ensured that the temperature was close to 0°C until the plants were processed. Upon arrival at the laboratory, the roots were separated from the aboveground parts, and the roots were subjected to surface sterilization. Subsequently, they were homogenized using a Warring blender combined with trehalose (at a concentration of 1 mg/g) and then lyophilized following a previously described method [10]. The obtained fine powder inoculant was stored at-80°C.

### Culturable bacteria density assessment and determination of salinity resistance in the inoculant microbiome

Bacterial density assessment was performed on Luria–Bertani (LB) agar obtained from BD Difco, Poland, supplemented with 100 µg/ml nystatin from Sigma Aldrich, Poland, to inhibit fungal growth. The agar plates were incubated for 7 days at 26°C.

For the estimation of salt tolerance, a previously published methodology [10] was employed. A Biolog Micro Station plate reader from Biolog, CA, was utilized for this purpose.

### Plant growth conditions

The plants were cultivated in a controlled environment using an artificially lit growth chamber. To ensure a clean and sterile growth environment, the ventilation outlets were equipped with HEPA filters. The air in the chamber was constantly subjected to sterilization using UV radiation emitted by flow lamps from ULTRAVIOL, Poland.

Throughout the experiment, the temperature in the growth chamber was maintained at a constant 20°C. The photoperiod followed a cycle of 16 hours of light and 8 hours of darkness, simulating a day/night cycle. LED lighting panels emitting white light were configured to provide a photosynthetically active radiation (PAR) intensity of 100 µmol * m-2 * s-1 at the soil level.

Watering of the plants was carried out as per their specific requirements using sterile deionized water.

### Inoculation

For plant inoculation, 100 mg of bioinoculant was used, which was an equivalent of ∼ 1 g of fresh roots. The bioinoculant was thoroughly mixed with the surface soil located in close proximity to the plants, covering an approximate radius of 1 cm. To facilitate the inoculation process, the plants were watered twice on the same day: once before inoculation and once after inoculation. On each occasion, half of the typical amount of water was administered.

### Sampling

Soil, root, and leaf samples were collected using sterile tools to maintain aseptic conditions. The soil samples were promptly snap-frozen in liquid nitrogen upon collection. Prior to freezing, the roots were subjected to surface sterilization, whereas the leaves were not subjected to sterilization. Additional information regarding the surface sterilization procedure can be found in the Supplementary Methods section, specifically in SM3.

### DNA extraction

DNA extraction was performed on all sample types using a combination of bead beating and flocculation, followed by purification using silica columns. The detailed protocols outlining the specific steps can be found in the Supplementary Methods section SM4.

### Library preparation and sequencing

To generate 16S rRNA gene fragment libraries, we followed the established protocol and made necessary modifications to reduce host rRNA amplification. The libraries were subsequently sequenced using MiSeq (Illumina, CA) at CMIT NCU, as previously described [44]. Further details regarding the modifications made for decreasing host rRNA amplification can be found in the Supplementary Methods section SM5.

### qPCR

Real-time PCR was conducted using a LightCycler 480 machine (Roche, Switzerland) along with a LightCycler 480 SYBR I Master kit (Roche, Switzerland) and Roche consumables. The reactions were carried out in a 10 µl volume, utilizing 1 ng of template DNA and 5 pmol of each primer per reaction. The specific primer sequences and cycling conditions can be found in Supplementary Methods SM6 and Table S1.

Each reaction was performed in four technical replicates, and purified amplicons were used to generate standard curves. The C_t_ values, determined using the second derivative algorithm of the LightCycler software, were exported into CSV files and further analysed using R as described in the SM.

### Bioinformatics and statistics

We denoised and merged the sequencing reads with DADA2 [45] and used the resulting amplicon sequence variants (ASVs) for downstream analyses. The sequences were then classified with the assignTaxonomy function of DADA2 using SILVA v.132 [46] as a reference database. A Relaxed Neighbor-Joining tree was constructed using clearcut [47], based on an alignment computed using Mothur v.1.44.3 [48] with SILVA v.132 as a reference as previously described [10]. Alpha diversity was assessed as Shannon’s H’, species richness (S) as the observed number of ASVs, while evenness was assessed as Shannon’s E (E=H’/ln(S)). For beta diversity analysis, we computed generalized UniFrac distance matrices based on the tree and ASV table using the GUniFrac package [49]. To ensure the comparability of alpha and beta diversity indices across samples with varying sequencing depths, we subsampled the ASV table 100 times to 900 sequences per sample. The averaged values, rounded to the nearest integer, were used in downstream analyses. Primers specific for particular ASVs were designed using the DECIPHER package [50]. We reconstructed the metabolic potential of the bacterial communities using PICRUSt2 [51] with default parameters. Inoculant influence was assessed with sourcetracker2 [52] on the non-rarefied dataset. We calculated reduced values for inoculated samples by subtracting the mean ‘influence’ for the respective non-inoculated samples. The relevant R and Mothur scripts can be found in Supplementary Methods SM7.

We compared the sample means with Kruskal-Wallis or Wilcoxon test using standard R functions or with robust ANOVA [53] implemented in the Rfit package. When applicable Benjamini-Hochberg correction FDR was used. The weighted NODF metric was used to assess the degree of nestedness, and the analysis was carried out using NODF software [54] run on Windows 7. We used the vegan R package [55] for ordinations and testing of grouping significance. Differences in community assembly process shares were assessed with the βNTI and Raup–Crick indices based on Bray–Curtis dissimilarity using the iCAMP package [29]. Differentially abundant ASVs, taxa and PICRUSt2-predicted traits were identified with DESeq2 [56] and, in certain cases, with ALDEx2 [57], while signature ASVs were identified with the biosigner package [58]. The code for performing the computations can be found in Supplementary Methods SM7.

## Results

### Beet plants generated by micropropagation of seedlings emerging from surface-sterilized seeds are nearly axenic

The beet seeds were virtually devoid of bacteria, regardless of genotype. Surface sterilization caused a decrease in bacterial 16S rRNA gene counts below the detection limit (Fig. SR2A). Seedlings emerging from sterilized seeds and propagated once on Murashige and Skoog media supplemented with cefotaxime and vancomycin (see Supplementary Method SM2) proved to be axenic (Fig. SR2B).

### The communities in the soils, roots and leaves differed significantly

The bacterial communities in the roots and leaves of axenic beet plants grown in pasteurized soils differed significantly in terms of structure (9.57% of variance explained, Fig. 2A), alpha diversity, where the diversity and richness of ASVs followed a pattern of soils > roots > leaves (Shannon’s H’: robust ANOVA F = 583.60, p < 0.001; Shannon’s E: F = 39.97, p < 0.001; richness: F = 583.60, p < 0.001), and bacterial load, which was lower in the leaves than in the roots (Wilcoxon test W = 259 411, p < 0.001) (Fig. 2B). Globally, ASV80, classified as *Achromobacter*, and ASV23, classified as *Chryseobacterium*, were found to be characteristic of (i.e., significantly more abundant in) plant tissues, while ASV3, ASV21 and ASV27 (*Pseudoxanthomonas*, *Sphingopyxis* and *Pedobacter*, respectively) were characteristic of soils. ASV7 (*Cellvibrio*) was typical for roots, and ASV13 (*Sphigobacterium*) was typical for leaves. Generally, differentially abundant ASVs affiliated with *Gammaproteobacteria* were characteristic of roots and soils, while *Bacteroidia*-associated ASVs were typical of leaves (Fig. 2E, Table SR3). At the genus level, *Bacillus*, *Brevundimonas*, *Pedobacter*, *Pseudoxanthomonas* and *Stenotrophomonas* were characteristic of soils, while *Cellvibrio* and *Flavobacterium* were more abundant in roots, and *Sphingobacterium* was more abundant in leaves (Fig. 2G, Supplementary ResultsF1). The core microbiome in the three compartments was limited to a few ASVs (8 in soils, 3 in roots and 1 in leaves; Table 1), which were mainly members of Alphaproteobacteria, *Gammaproteobacteria* and *Bacteroidia*.

**Fig 2.**
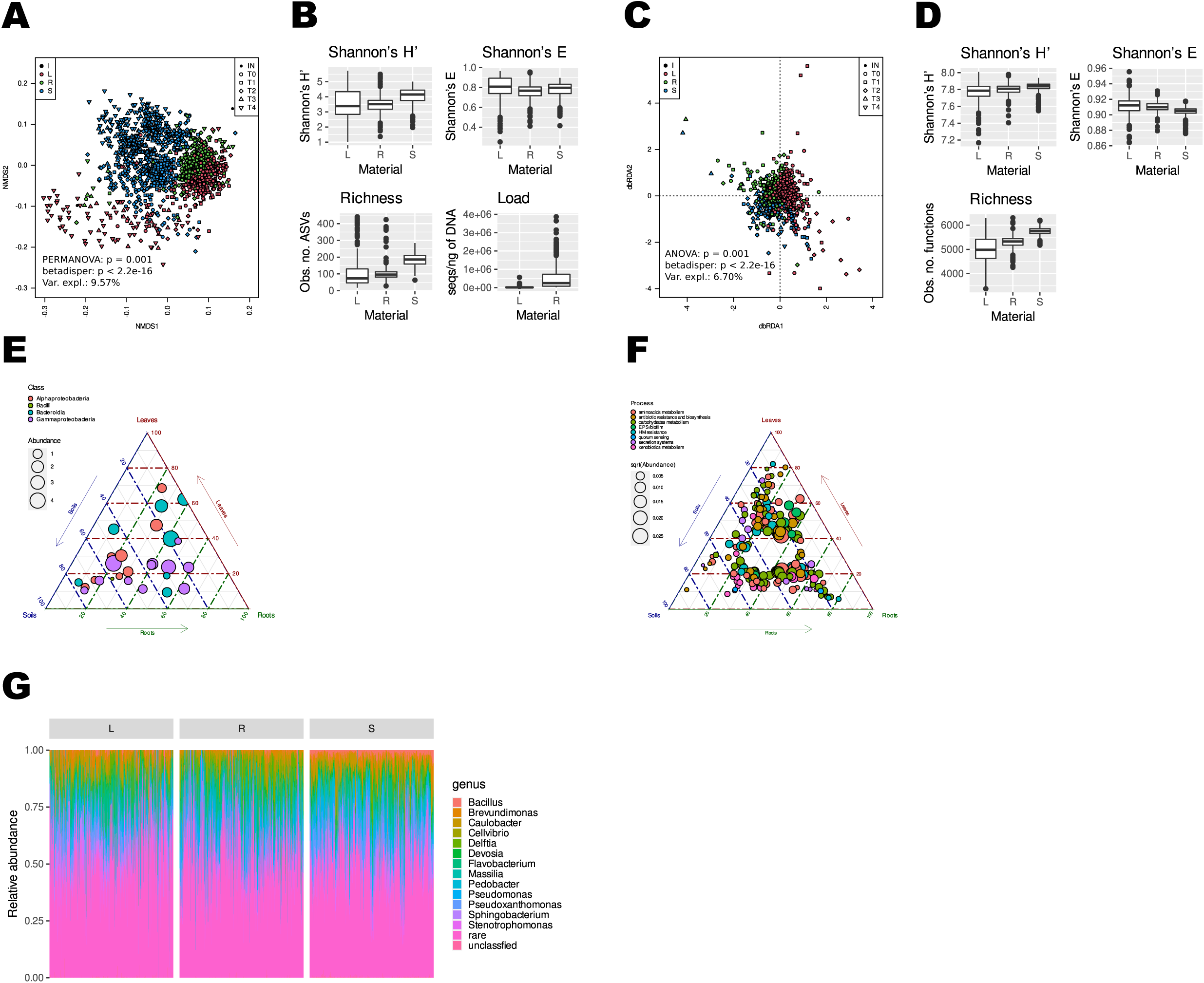
Bacterial communities and sets of functions encoded in their genomes (functional potential) in soils, roots and leaves differ in terms of structure, alpha diversity and bacterial load. L – leaves, R – roots, S – soils, I – inoculant (lyophilized wild-growing sea beet roots). A) NMDS analysis of the d05 generalized UniFrac distance matrix; black, inoculant (lyophilized wild sea beet roots); blue, soils; green, roots; red, leaves; shape, timepoint; circles, two weeks after planting in soil (T0); squares, fifteen days (T1); diamonds, three weeks (T2); upward-pointing triangles, four weeks (T3); downward-pointing triangles, six weeks (T4). B) Alpha diversity of ASVs. Boxplots of Shannon’s diversity index (H’)-upper left, Shannon’s evenness – upper right, richness (observed number of ASVs) – lower left, number of bacterial 16S rRNA gene sequences per ng of DNA – lower right. C) dbRDA of the Morisita-Horn distance matrix derived from functional sets imputed by PICRUSt2; D) alpha diversity of functions. Boxplots of Shannon’s diversity index (H’) (upper left), Shannon’s evenness (upper right), and richness (observed number of functions) (lower left). E) Ternary plot of ASVs displaying differential abundance in soils, roots, and leaves. The mean ASV abundance is shown as a circle, and the color denotes the classification at the class level: Alphaproteobacteria, red; Bacilli, green; Bacteroidia, blue; and Gammaproteobacteria, purple. F) Ternary plot of pathways displaying differential abundance in soils, roots and leaves; the scale is arbitrary. The color indicates which broad category a given pathway belongs to: antibiotic resistance and biosynthesis – red, carbohydrate metabolism – yellow‒green, heavy metal resistance – green, quorum sensing – blue, and secretion systems – purple. G) Taxonomic structure of the soil (S), root (R), and leaf (L) communities at the genus level; the genera identified by DESeq2 as significantly more abundant in a specific compartment are marked with a letter denoting the compartment: s – soil, r – root, l – leaf.

**Table 1.**
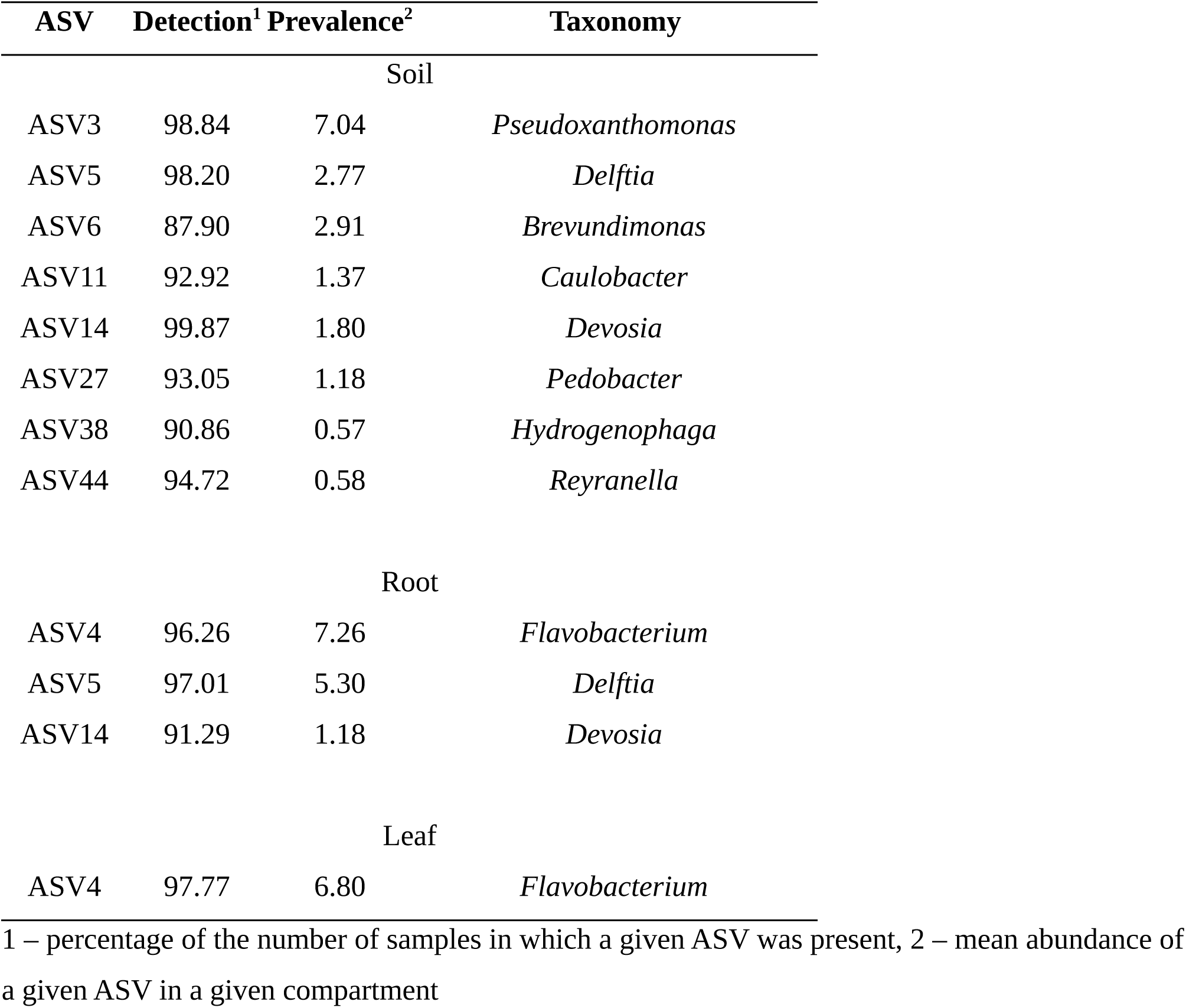
Core microbiomes of the soil, root and leaf samples. A prevalence cutoff of 0.1% and a detection rate of 90% were used.

| ASV | Detection<sup>1</sup> | Prevalence<sup>2</sup> | Taxonomy |
| --- | --- | --- | --- |
| Soil |  |  |  |
| ASV3 | 98.84 | 7.04 | *Pseudoxanthomonas* |
| ASV5 | 98.20 | 2.77 | *Delftia* |
| ASV6 | 87.90 | 2.91 | *Brevundimonas* |
| ASV11 | 92.92 | 1.37 | *Caulobacter* |
| ASV14 | 99.87 | 1.80 | *Devosia* |
| ASV27 | 93.05 | 1.18 | *Pedobacter* |
| ASV38 | 90.86 | 0.57 | *Hydrogenophaga* |
| ASV44 | 94.72 | 0.58 | *Reyranella* |
| Root |  |  |  |
| ASV4 | 96.26 | 7.26 | *Flavobacterium* |
| ASV5 | 97.01 | 5.30 | *Delftia* |
| ASV14 | 91.29 | 1.18 | *Devosia* |
| Leaf |  |  |  |
| ASV4 | 97.77 | 6.80 | *Flavobacterium* |

1 – percentage of the number of samples in which a given ASV was present, 2 – mean abundance of a given ASV in a given compartment

**Table 2.**
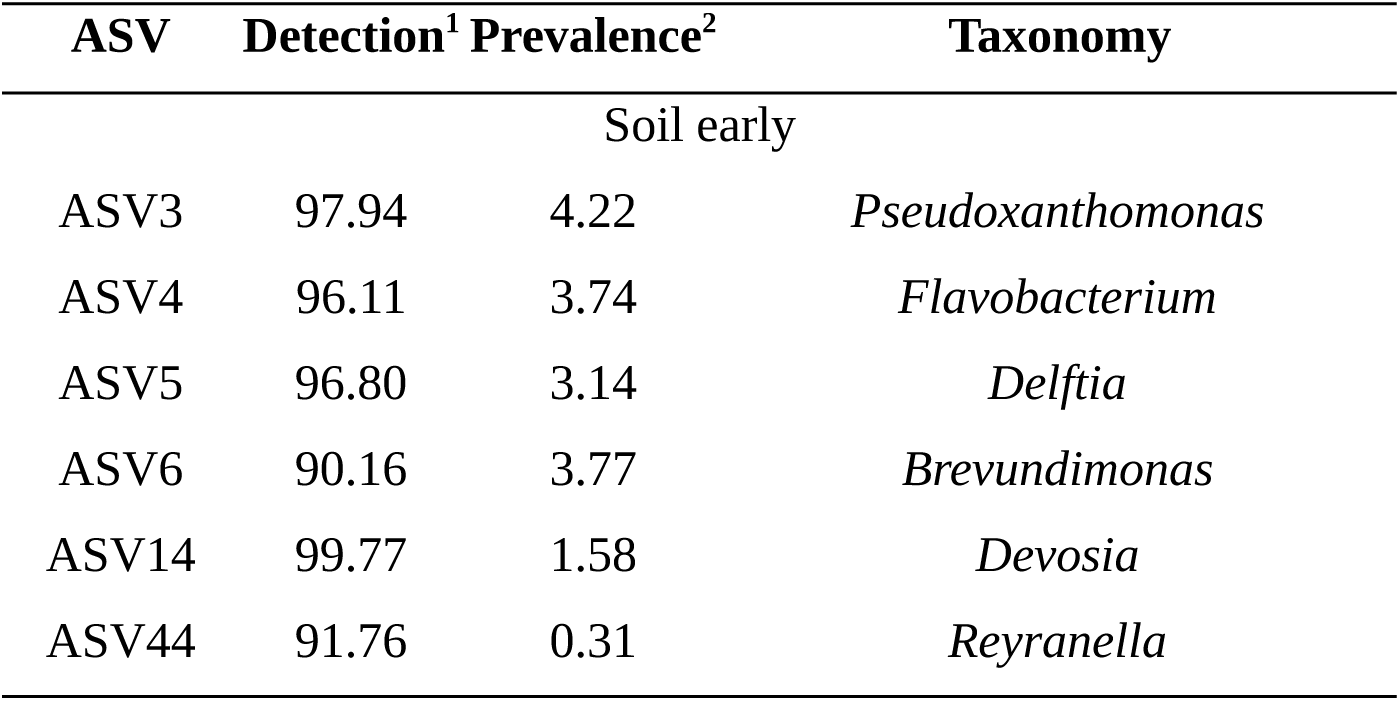
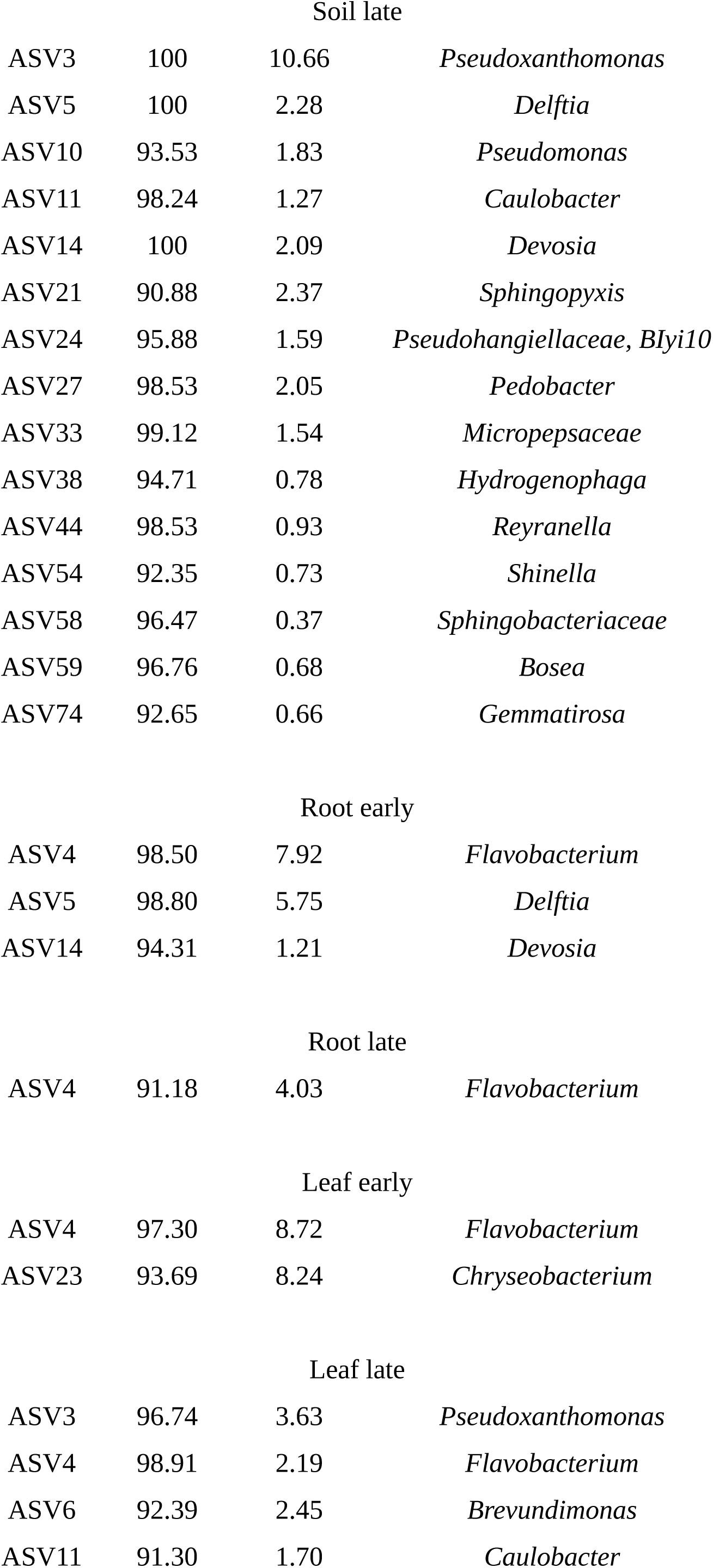
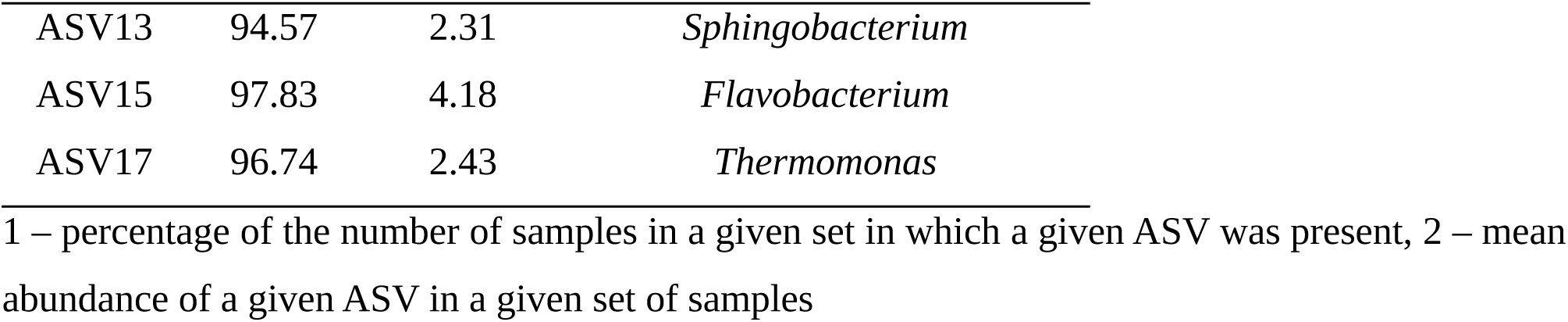
Late and early core microbiomes. A prevalence cutoff of 0.1% and a detection rate of 90% were used.

**Tabel 3.**
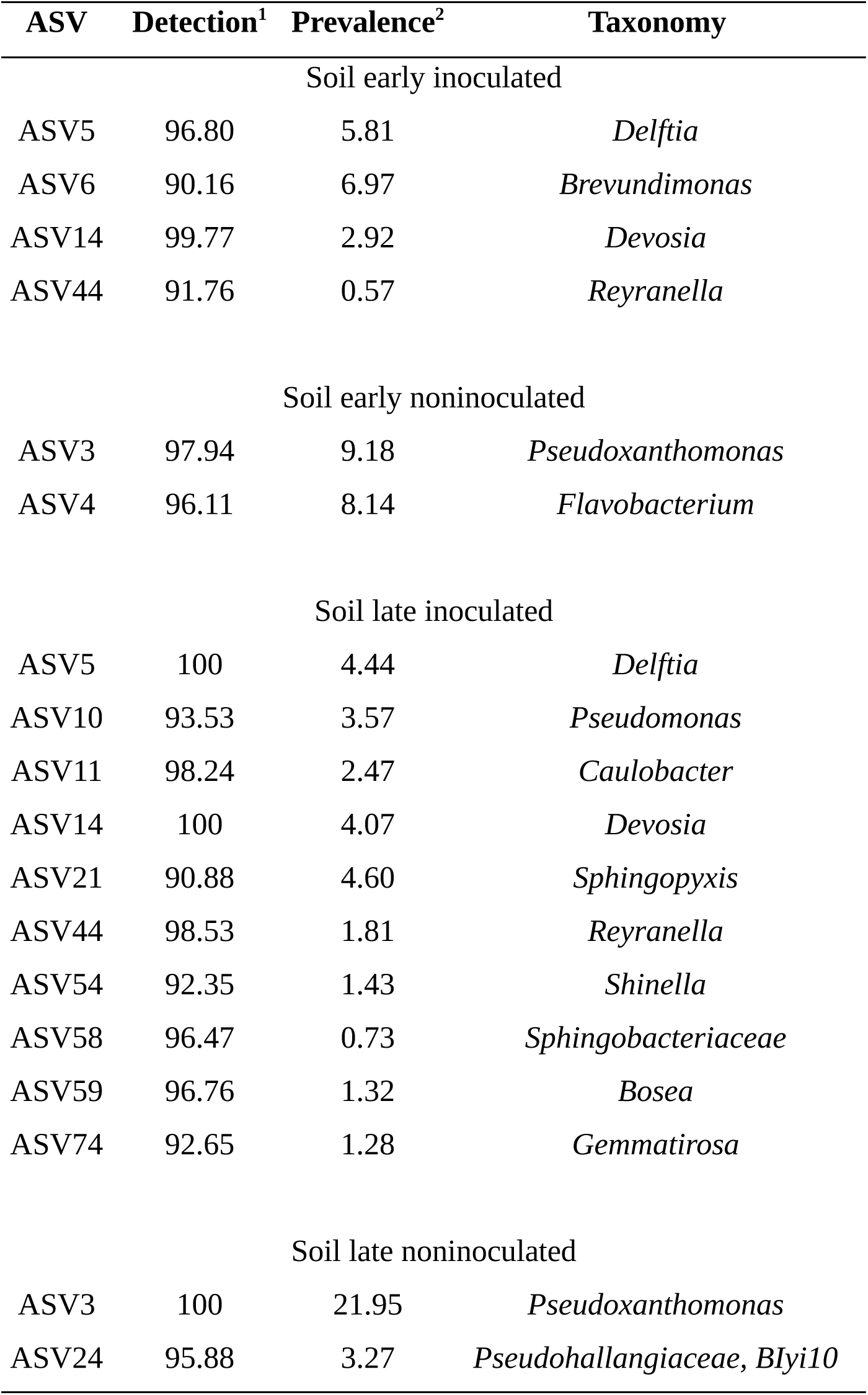
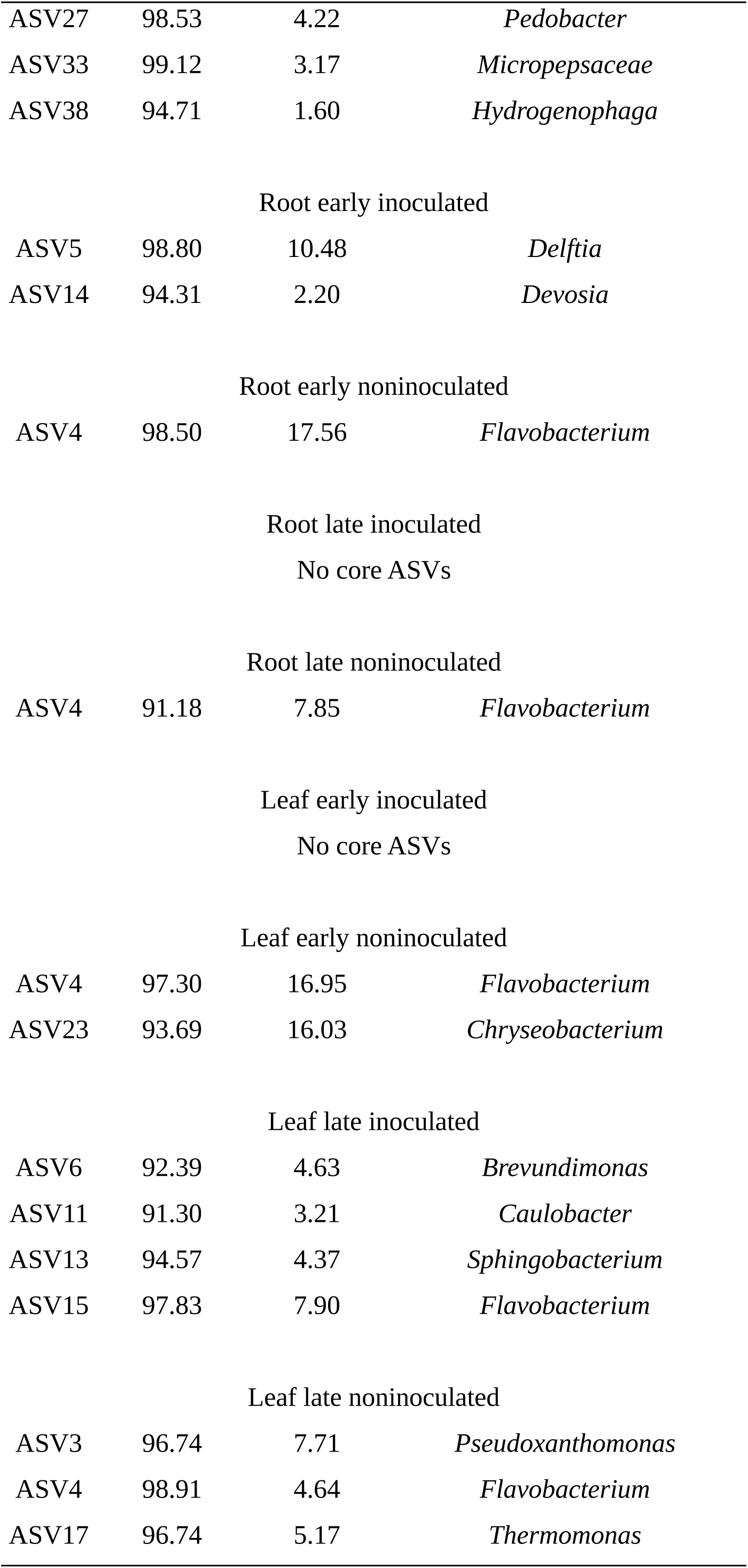
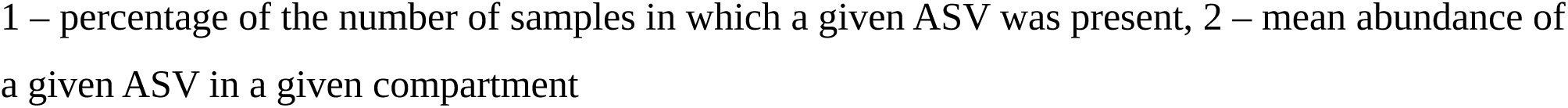
Core microbiomes of inoculated and noninoculated samples. A prevalence cutoff of 0.1% and a detection rate of 90% were used.

The differences, albeit smaller, were also visible at the level of PICRUSt2-predicted metabolic capabilities, which were also grouped according to material (Fig. 2C, 6.70% variance explained), and the alpha diversity of functions followed the pattern observed for ASVs (Fig. 2D, Shannon’s H’: robust ANOVA F = 104.44, p < 0.001; Shannon’s E: F = 173.20, p < 0.001; richness: F = 851.62, p < 0.001). Functions related to competition between microorganisms (antibiotic resistance and biosynthesis, quorum sensing) appeared to be characteristic of soils and roots, while carbohydrate metabolism-related functions were predicted to be more frequent in the genomes of soil-and leaf-dwelling bacteria (Fig. 2F; Table SR4).

As material explained a far greater fraction of the variance than any other variable (Table SR5), to determine the influence of other variables, further analyses were carried out on the data divided into soil, root and leaf sets.

### The first weeks of axenic beet growth in soil can be divided into two stages differing in community structure, diversity, bacterial load, predicted metabolic capabilities and nestedness

The time point was the second most important grouping variable, regardless of the material (5.82% of the variance explained in the whole dataset). Three ‘early’ time-points (T0, T1 and T2) clustered together and were significantly different from the ‘late’ time-points (T3 and T4). The percentage of variance explained by this grouping was 15.64% in the leaves, 3.05% in the roots and 9.67% in the soils (Fig. 3A). Henceforth, the belonging of a sample to the ‘early’ or ‘late’ cluster will be called its ‘status’.

**Fig. 3.**
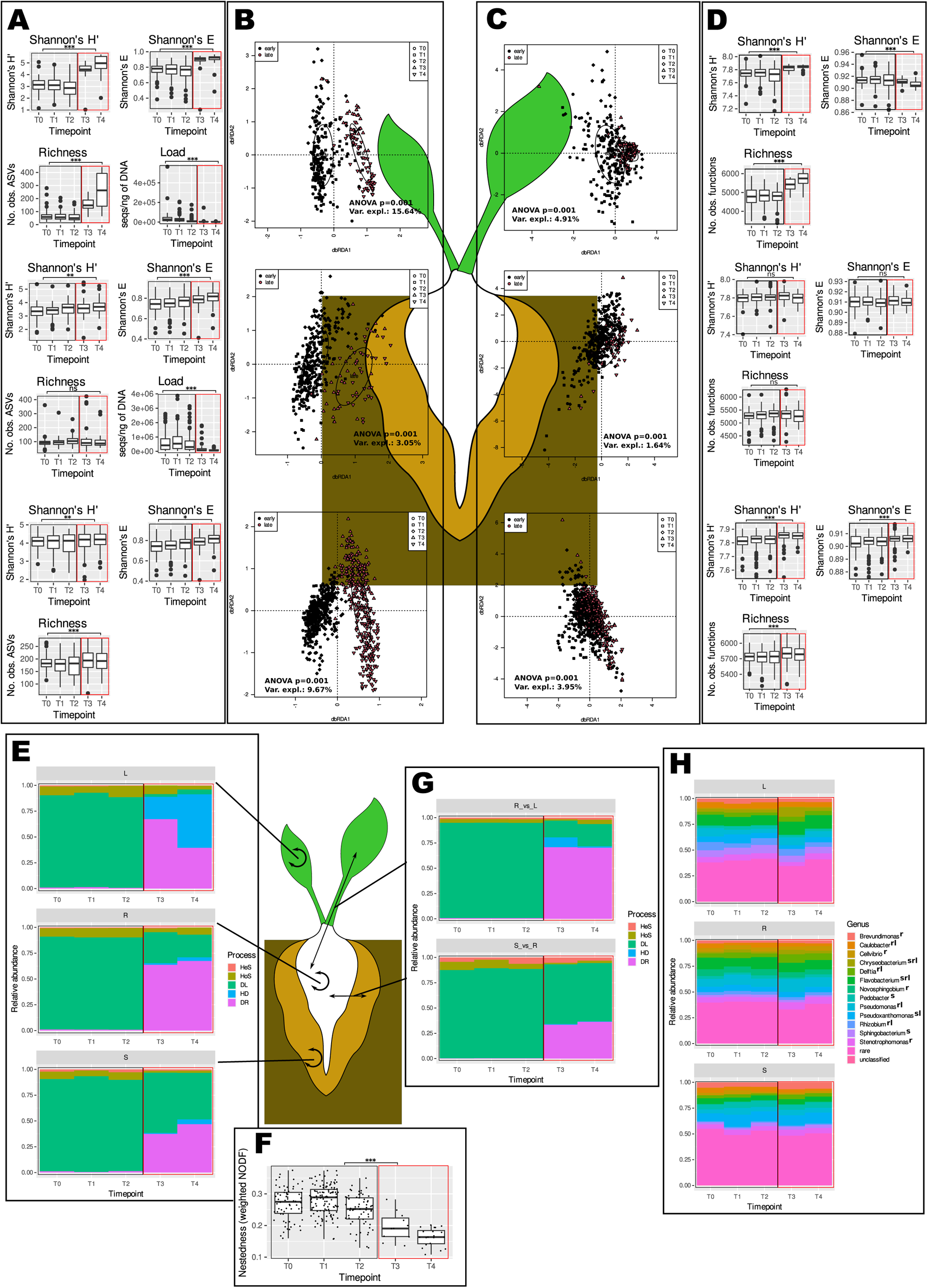
The bacterial communities and their functional potentials differed between the samples collected until the 35th day post planting (early samples) and those collected later (late ones). The graphs in panels A, B, C, D, E, and H show (top to bottom) leaves, roots and soils. The black outlines indicate early samples, while the red outlines indicate late samples. The significance of differences in means (Wilcoxon’s test) in panels A, D and F is shown: *** – p < 0.001, ** – p < 0.01, * – p < 0.05, ns – not significant. A) The alpha diversity, evenness, and species richness of ASV sets are greater and the bacterial load is lower in late samples than in early samples. B) Communities differ according to dbRDA analysis of d05 generalized UniFrac distance matrices (results of a permutational test are shown on graphs). C) The functional potential of bacterial communities differs according to dbRDA analysis of Morisita-Horn distance matrices (results of a permutational test are shown on graphs). D) Alpha diversity, evenness and functional richness are greater in soil and leaf samples but not in roots. E) Dispersal limitation and homogenous selection shape communities in early samples, while random processes (drift) and dispersal limitation (in the case of soils and roots) or homogenizing dispersal (in the case of leaves) shape late communities. F) Nestedness is low in all samples but lower in late than in early ones. G) Dispersal limitation and selection (mostly homogenous) govern the transfer of organisms from soil to roots and from roots to leaves in early samples, while random processes (drift) are more important in late ones. H) Taxonomic structure at the genus level is different in early samples than in late ones. Differentially represented (DR) taxa were identified with DESeq2 and ALDEx2 (taxa identified by both methodologies were considered DR), and the upper indices next to the taxon names indicate the kinds of samples in which there was a difference: s – soils, r – roots, and l – leaves.

Differences between the early and late clusters were observed regardless of the material, soil and genotype (Figs. SR4,5,6) and were also visible in the alpha diversity measurements, which were greater, and, in the case of the plant tissue samples, in the bacterial load, which was lower in the late samples (Fig. 3B). The former effect was most pronounced in leaves and least pronounced in soils, while the decrease in the number of bacterial 16S rRNA gene sequences was greater in roots than in leaves. Different organisms were characteristic of early and late samples in soils, roots and leaves. The organisms characteristic of soils and roots were of low abundance (Fig. SR8ACE; Supplementary Results F1).

The traits characteristic of the genomes of organisms thriving in late and early samples differed among the soils, roots and leaves (Fig. 3C; SupplementaryResultsF2). Early soils harboured organisms whose genomes were enriched in genes involved in diverse array of functions, among which genes related to methane metabolism, protein and nucleotide rescue from glyoxal glycation, and heavy metal resistance were most prominent. On the other hand, the metabolism of aromatic compounds was characteristic of the genomes of organisms dwelling in late soil samples (Fig. SR9F). Genes involved in root biofilm formation, exopolysaccharide synthesis and the regulation of the amino acid pool were characteristic of early samples, and the metabolism of aromatic compounds was characteristic of late samples (Fig. SR9D). Toxin/antitoxin systems were characteristic of leaves in general, polysaccharide (chitin, pectin) utilization was of greater abundance in early leaf samples, while aromatic compound metabolism was typical of late leaf samples (Fig. SR9B). The functional diversity was significantly greater in the late leaf and soil samples than in the root samples (Fig. 3D).

The degree of nestedness calculated for the soil–root–leaf matrices (for each plant (technical replicate) separately) was very low, essentially did not deviate from the expected values derived from a null model (Supplementary File 3) and decreased with time. The difference between the early and late samples was significant (Fig. 3F; Table SR6).

Dispersal limitation (DL) dominated mechanisms governing the entry of bacteria into roots and their transfer to leaves in early samples, while other stochastic processes (drift) were more pronounced in late samples. The share of DL was greater in the case of soil → root transfer than in the case of root → leaf transfer. Interestingly, the levels of selection, albeit generally low, were greater in the early samples than in the late samples (Fig. 3G). A similar pattern was observed when maintenance of the soil, root and leaf communities was assessed (i.e., samples from the same biological replicate were compared); however, in the case of leaf communities, DL was replaced with homogenizing dispersal (Fig. 3E).

### Inoculation with lyophilized wild beet roots influences bacterial communities in soils and plants

#### Inoculant characterization

Reads affiliated with *Pseudomonadota* (formerly *Proteobacteria*), *Bacteroidota* (formerly *Bacteroidetes*) and *Bacillota* (formerly *Firmicutes*) were found in libraries prepared from DNA isolated from inoculant samples. The most abundant genera were *Pseudoxanthomonas* and *Brevundimonas* (>5% each), while *Pedobacter*, *Devosia, Caulobacter, Flavobacterium, Rhizobium, Sphingobacterium, Pseudomonas*, *Cellvibrio, Thermomonas,* and *Dyadobacter* were less abundant (∼2-5%; Fig. 4A). A total of 55% of the reads were rare genera (<2% abundance). The diversity, measured as Shannon’s H’, was 4.40 ± 0.75, the evenness was 0.93 ± 0.003, and 144 ± 125 ASVs were detected in the inoculated samples (rarefied data, n = 3), while 437 ASVs were detected in the non-rarefied dataset. The cultivable bacterial density was 4.0 ± 0.09 × 105 cfu/g (n = 6), while the bacterial 16S rRNA count was 2.0 ± 0.5 × 10^4^ copies/ng of DNA, which translates to 1.4 ± 0.587 × 10^8^ copies/g of inoculant (n = 8). The bacteria were able to grow in the presence of up to 900 mM NaCl (Fig. 4B).

**Fig. 4.**
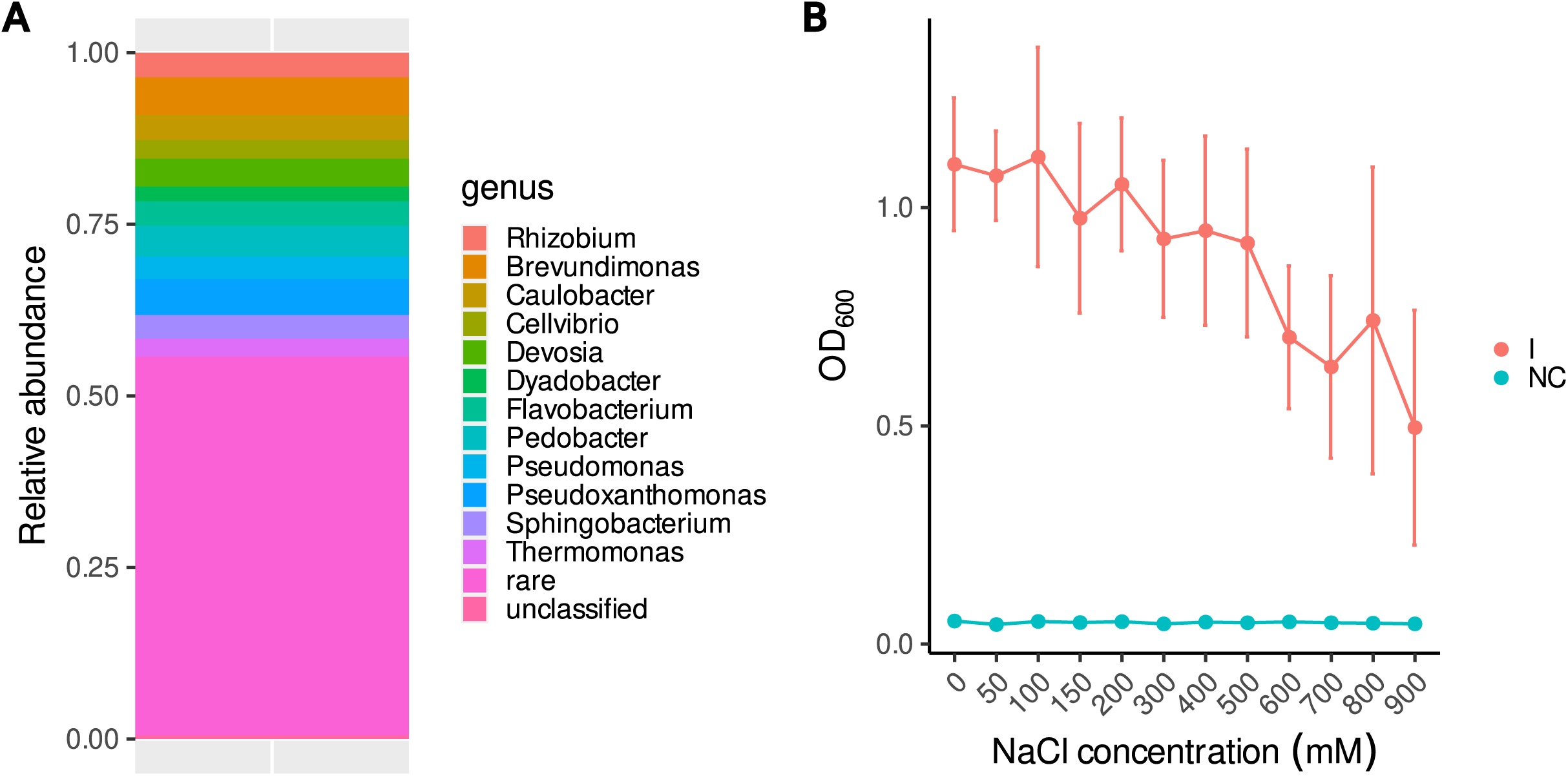
Inoculant characterization. A – Taxonomic structure of the bacterial community. B – bacteria from the inoculant were able to grow in the presence of up to 900 mM NaCl; I – inoculant, NC – noninoculated control.

### The community structure in inoculated and noninoculated samples differed significantly, and the differing ASVs differed depending on compartment, soil and genotype

Inoculation had no influence on bacterial alpha diversity (Fig. 5A), and its effect on bacterial community structure was small but significant in each of the studied compartments and greater in soils than in roots or leaves (0.48, 0.17 and 0.09% of explained variance, respectively, Fig. 5B). The bacterial load in the plant samples did not differ between the inoculated and non-inoculated plants, regardless of the experimental treatment (Fig. 5A and Fig. SR35D). Further analyses showed that inoculation significantly impacted the bacterial communities in all the experimental soil treatments, regardless of their status, but only in certain variants in the case of the plant samples (Figs. SR32-34, Table SR7). The mean d05 generalized UniFrac distance between the inoculated samples and the inoculant was surprisingly slightly greater (0.3810 ± 0.0547) than that between the inoculated samples and the non-inoculated samples (0.3699 ± 0.0554), and the difference was significant (Wilcoxon test, W = 3.3482e+10, p < 0.001).

**Fig. 5.**
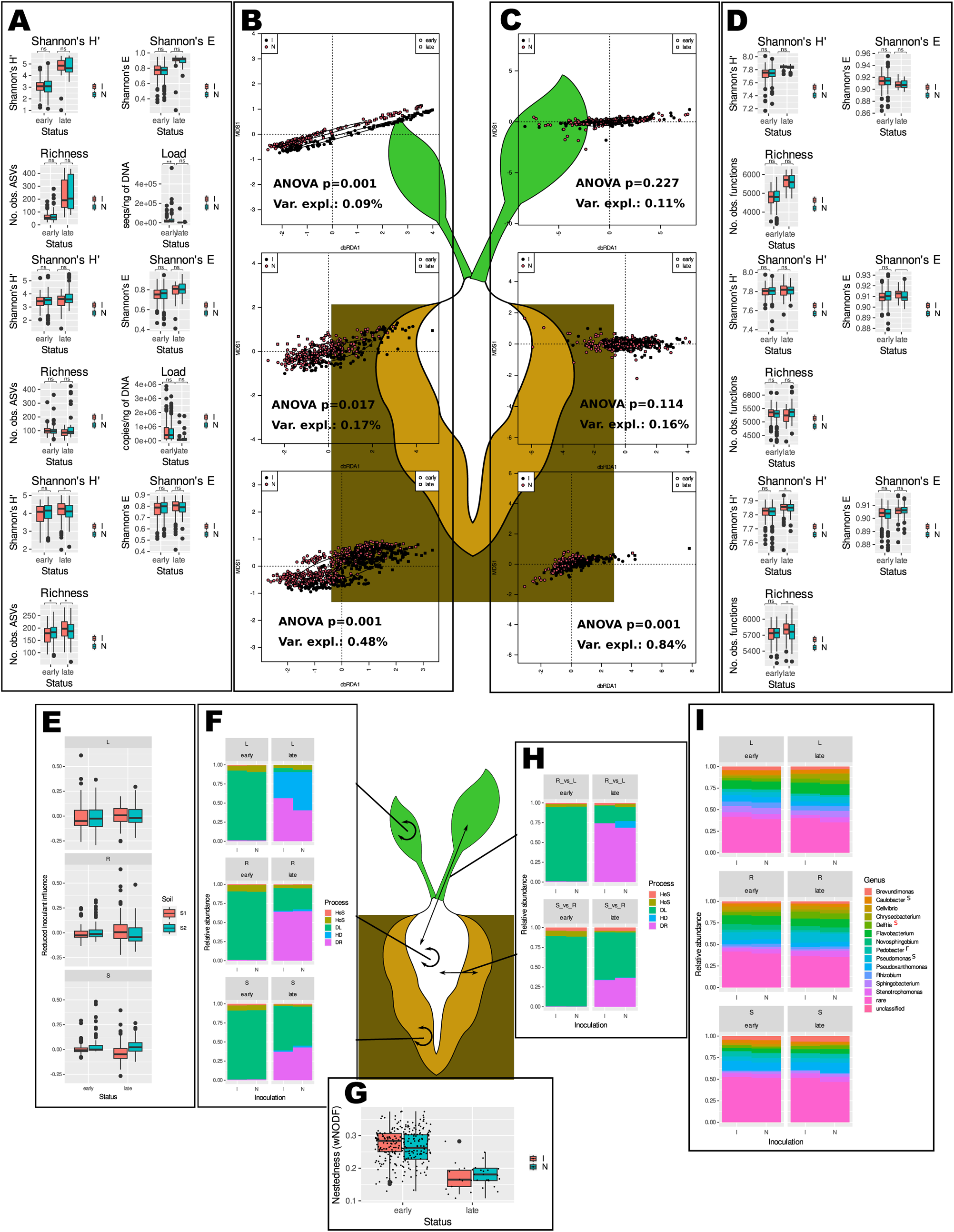
Inoculation slightly influenced the bacterial community structure and functional potential but had no effect on the alpha diversity, taxonomic composition, nestedness or community assembly processes. The graphs in panels A, B, C, D, E, F and I show (top to bottom) leaves, roots and soils. The significance of differences in means (Wilcoxon’s test) in panels A and D is shown: *** – p < 0.001, ** – p < 0.01, * – p < 0.05, ns – not significant. A) Diversity, evenness, species richness and bacterial load did not differ between inoculated and noninoculated samples, with the exception of soil richness. B) Inoculated and noninoculated samples clustered together regardless of compartment. dbRDA analysis of d05 generalized UniFrac distance matrices (results of a permutational test are shown on graphs); C) Sets of functions predicted to be encoded in genomes of bacteria from inoculated and noninoculated samples differ regardless of material (results of a permutational test are shown on graphs); D) Diversity, evenness, and richness of predicted functions do not differ in inoculated and noninoculated samples; E) Reduced inoculant contribution to inoculated samples is very low and does not differ between soils; F) There is no difference in shares of community assembly processes between inoculated and noninoculated samples regardless of material; G) Nestedness in soil-root-leaf sets is not influenced by inoculation; H) Processes governing transfer of bacteria from soils to roots and from roots to leaves are not influenced by inoculation; I) Taxonomic structure of bacterial communities at the genus level is similar in inoculated and noninoculated samples. Differentially represented (DR) taxa were identified with DESeq2 and ALDEx2 (taxa identified by both methodologies were considered DR), and the upper indices next to the taxon names indicate the kinds of samples in which there was a difference: s – soils, r – roots, and l – leaves. The colour denotes the status: early (black) or late (red).

The effect of inoculation was even smaller for the predicted functional potential, both for alpha-diversity (Fig. 5D) and beta-diversity (Fig. 5C), and in the plant samples, the effect was not significant. Inoculation influenced only early samples of genotypes C and M and late samples of genotype B in soils (Figs. SR36-38 and Table SR7).

The nestedness level did not change in response to inoculation (Fig. 5F), and differences in the proportions of community assembly processes were visible only in the case of late leaves, where the percentage of homogenizing dispersal was lower in inoculated samples (Fig. 5E and 5G).

Taxonomic composition at the genus level was similar in inoculated and non-inoculated samples, and only three non-rare genera were differentially abundant in soils, while one was differentially abundant in roots (Fig. 5H, Supplementary Results F1). The sets of ASVs and functional characteristics of the inoculated and non-inoculated samples were different in the early and late samples as well as in each experimental variant (Figs. SR40 and SR42-47, SR41 and SR48-53, respectively, as well as SupplementaryResultsF1 and SupplementaryResultsF2). Of the 437 ASVs detected in the inoculant, 268 were found exclusively in the inoculated samples, although they were rare (lowly abundant). However, when rarefied data were used, only fifteen such ASVs were found (Table SR9). No ASV present in the inoculant was found only in the non-inoculated samples. Globally, 29 ASVs were identified with a biosigner as a signature for both inoculated and non-inoculated samples. The ASVs were classified mainly as *Alpha*-and *Gammaproteobacteria* as well as *Flavobacteria* and *Chitinophaga* (Table SR8). Characteristic ASVs could be found mainly in soils; in the case of plant samples, they were detected only in certain plant variants. The organisms that differentiated inoculated late samples from non-inoculated samples were different for each combination of material, soil and genotype. The influence of inoculation was most visible in the soil samples (the greatest number of ASVs differentiating between the inoculated and non-inoculated samples), while in the roots and leaves, there were only single ASVs in certain variants. These bacteria belonged mainly to *Proteobacteria* and *Firmicutes*. Differences in functional potential comprised diverse functions, and genes involved in antibiotic biosynthesis and resistance were frequently found to be more highly represented in inoculated samples than in non-inoculated samples, potentially indicating an increased level of competition.

## Discussion

### How much time is needed to establish stable endophytic communities in axenic plants of different genotypes grown in various soils?

Despite differences in community composition caused by soil and beet genotype, identical patterns were found in each experimental variant: samples collected until the 35th day after planting in soil (early ones) were similar in all measured parameters, and the same was true for the samples collected after day 35 (late ones). Such a situation seems to be common in microbial succession (e.g. [27]) and is similar to the classical plant succession model of Cowles [59]. Interestingly, the evolution of the rhizosphere communities over time followed the pattern described above. On the one hand, this similarity might be interpreted as a result of the rhizosphere community being driven by plant developmental stage, possibly via changes in root exudation (reviewed, e.g., in [38]). On the other hand, it was found that time is a stronger driver of rhizosphere bacterial community structure than plant development [60].

Changes in endophytic communities over time due to developmental stage and seasonality have been demonstrated in various plants, both perennial [23] and annual ones [24], and might be considered an instance of succession consisting of stages at which different community assembly mechanisms are important [61]. We are not aware of a plant study reporting such microbial succession in axenic, *in vitro*-propagated plants. In the case of plant colonization by microbes, succession should be modulated by plant development, as conditions in the plant interior depend on developmental stage, which has been demonstrated, e.g., for beet roots [36], and could be influenced by host genotype. Indeed, both soil type and genotype influenced both rhizosphere and endophytic community structure but not alpha diversity. We expected that sea beet, as a wild plant, would recruit more diverse bacterial communities than sugar beet cultivars, as was the case for other plants, e.g., wheat [62], both at the level of taxonomy and function. As wild plants need to cope with a broader spectrum of environmental conditions, we assumed that they would need greater microbiome functional potential. However, there was no clear trend in alpha diversity, suggesting that under the conditions used here, there was no such need. Revealing the greater plasticity of wild beet compared to cultivars would require more adverse conditions, such as drought, salinity or infertile soil. A relatively small genotype effect was also found in other studies, e.g., on the cotton rhizosphere [63] or in the willow rhizosphere and root communities [64].

In this study, we observed a decrease in the bacterial load in late samples. The lower load could be due to several factors: greater selection leading to the elimination of certain organisms, dilution caused by increases in plant tissue volume and weight and bacterial cell division arrest, which may result from plant-or microbe-derived compounds responsible for quorum sensing. Alternatively, bacteriostatic compounds might be produced by plants or bacteria.

The succession phases observed here seem to be in line with beet taproot growth phases, with our early phase corresponding to transition/secondary growth onset and the late phase corresponding to the beginning of the sucrose storage phase [36]. Therefore, we believe that under stable environmental conditions, the late phase could last at least until flowering (in the case of a perennial—sea beet) or overwintering (in the case of a biannual—sugar beet), which are the next major events in beet life. In this sense, the beet endophytic and rhizosphere communities became stable three weeks after the axenic plants were planted in the soil.

### Do community assembly processes and nestedness differ over time and for various genotypes?

The fact that alpha diversity followed a pattern of soils > roots > leaves prompted us to perform nestedness analyzes on soil, root and leaf triplets, which allowed us to speculate on possible community assembly mechanisms [28]. The NODF index was consistently low in all variants and, in most cases, did not deviate from that expected by chance, which, together with little overlap between the soil, root and leaf communities, points to random colonization as the prevailing community assembly mechanism [28]. Nestedness might be influenced both by random (e.g., random sampling (colonization) or incidental death) or deterministic (environmental filtering, selection or extinction) processes. The significant difference in the degree of nestedness between the early and late phases and between the sea beet and sugar beet varieties suggested that the proportions of stochastic/deterministic processes might also differ over time. To corroborate these results, we predicted the shares of assembly processes using the iCAMP package. As in other systems (e.g., glacier forefront [65] or field after nudation [66]), stochastic community assembly processes dominated both in the early and late phases, explaining the low level of nestedness. High shares of stochastic community assembly mechanisms may be caused by the fact that soil bacterial communities, from which endophytic organisms are recruited, are highly functionally redundant (i.e., there are many organisms with a given trait/set of traits). This view is supported by predicted gene content sets being closer (less separated) than sets of ASVs. However, on the one hand, high similarity of marker sequences does not necessarily mean highly similar gene content (e.g., due to horizontal gene transfer), and on the other hand, even genomes dissimilar in terms of a marker sequence encode a set of core functions and may share non-core traits, as indicated by pangenome conception [67].

The high level of randomness observed in our system may also be attributed to relatively low coverage, as the abundance of some organisms might have fallen below the detection level, thus decreasing community similarity. Dispersal limitation was commonly thought to be the most important mechanism shaping communities at early stages of succession, which was also observed in other systems (reviewed in). However, there are also studies reporting selection as the main microbial community assembly mechanism during plant development [16]. All this said, the predictions might be inaccurate due to inherent limitations of the iCAMP methodology [29] and low sequencing depth (as discussed earlier). In particular, dispersal limitation may be considered both a deterministic process that is difficult to distinguish from selection and a random process [68]. Currently, there is no means of partitioning DL to deterministic and random components. It seems plausible that dispersal limitation and selection act in concert during bacterial colonization of axenic plants and that the proportions of these two mechanisms might depend on environmental conditions. The low level of selection detected in our study suggested that under optimal growth conditions, plants do not exert high selective pressure on bacteria; however, the apparent low selection might stem from high functional redundancy in the soil bacterial community. The low selection level also points to dilution because plant growth is the mechanism causing a decrease in the bacterial load in late plant samples.

### How does inoculation of the wild sea beet root community influence bacterial communities in axenic beets?

Inoculation influenced beet-associated bacterial communities only slightly but significantly. Because we used lyophilized roots as an inoculant, its effect may stem not only from bacteria being introduced to soil but also from organic matter (particularly nitrogen compounds and carbohydrates). The typical composition of beet roots suggests that in 100 mg of lyophilizate, we expected that the influence of the inoculant would be greater in less N-and OM-rich soil (S1).

Conversely, the influence was smaller in S1 than in S2, and we concluded that this difference was mainly caused by the organisms added to the soil.

As we expected, the influence of inoculation was small; therefore, we grew axenic plants in pasteurized soils. The presence of a handful of ASVs from inoculant in inoculated plant samples and their absence from non-inoculated ones suggested that organisms thriving in the inoculant entered the plants. However, it is possible that they were present in non-inoculated samples below the detection threshold—as always, it is impossible to prove that an entity is not present in a given sample. The low abundance of inoculant-derived endophytes shows that they do not increase community dissimilarities directly and suggests that microbe‒microbe interactions and/or fertilization effects are responsible for the majority of differences in bacterial community structure observed between inoculated and non-inoculated samples. The greatest inoculation influence in late soil and early plant samples suggested that the selective pressure in soils is weaker than that in plants and/or organisms not able to enter plants that start growing during week 8 of growth in soil.

We expected that inoculation would cause homogenization of communities (i.e., would make the mean distance between inoculated communities smaller than that between non-inoculated communities), but this did not prove to be true. It seems that the inoculant increases the number of organisms that may be recruited by plants, causing a decrease in similarity. Notably, inoculation did not change time-or genotype-driven beta diversity patterns, which agreed with the low values of variance explained by this variable. The fact that the bacterial load was lower in the later samples suggested that the appropriate inoculation time may be crucial for the successful application of biofertilizers, at least in beet. However, a definite answer to the question of the right time to inoculate would require a carefully designed experiment, as it is also plausible that bacteria introduced to soil during the late phase of colonization might have a better chance of entering plants due to a lower level of selection and dispersal limitation in this phase.

## Conclusions

Regardless of the soil type and genotype, the colonization of axenic beet plants occurred in at least two phases—up to the 35th day of growth in soil—and lasted until the end of the experiment, akin to microbial primary succession in other environments. Plants govern bacterial succession in rhizosphere soil, as they follow the same temporal pattern as succession in the endosphere. Bacterial communities are compositionally stable, and the bacterial load and nestedness are much lower in the late phase than in the early phase. Colonization is largely random at the taxonomic level— various strains (ASVs) encoding similar functions are recruited from a pool of functionally redundant soil bacteria. Regardless of the soil type and genotype, inoculation slightly influenced the bacterial communities, but significantly fewer organisms were recruited by the plants. The scarcity of inoculant-derived strains in the endosphere suggests that inoculation acts mostly indirectly, probably via microbe‒microbe interactions. As both bacterial cell entry to the endosphere and bacterial division seem to be arrested in the late phase, early application of bioinoculants seems to be the right choice.

## Supporting information

Supplementary Results

Supplementary ResultsF1

Supplementary ResultsF2

Supplementary ResultsF3

## Acknowledgments

We are grateful to Prof. Wiesław Nowak and Dr. Łukasz Pepłowski for making the computational resources available.

## List of abbreviations

NGI – next generation inoculant

## Declarations

Ethics approval and consent to participate

Not applicable

## Consent for publication

Not applicable

## Availability of data and materials

The datasets generated during this study are available in the NCBI SRA repository under BioProjects no. **PRJNA615328** (soil communities), **PRJNA713813** (root communities), **PRJNA713992** (leaf communities) and **PRJNA847007** (inoculants and some leaf and root communities). Files with differentially represented features (ASVs, higher taxa, predicted KO functions and pathways) as well as raw qPCR results and colony count data are available on figshare (10.6084/m9.figshare.24230686, 10.6084/m9.figshare.24298081, 10.6084/m9.figshare.24298135, 10.6084/m9.figshare.24298327, 10.6084/m9.figshare.24298405, <u>10.6084/m9.figshare.20652645</u> and <u>10.6084/m9.figshare.20652915</u>). R scripts allowing the generation of all figures and tables are part of Supplementary Methods SM7. The materials used in this work, excluding the samples used in their entirety, are available from the corresponding author upon reasonable request.

## Authors’ contributions

MG conceived the study, obtained financing, designed the experiments, participated in laboratory work, analyzed the data, supervised the study and drafted the manuscript, MS performed the laboratory work, participated in writing, JM performed the laboratory work, SS performed the laboratory work, JT participated in designing the experiments, performed the laboratory work, participated in writing, KH participated in designing the experiments, participated in writing, WU analyzed the data and participated in writing. All the authors have read and approved the final manuscript.

