## Supplementary Methods for "Rhizosphere bacterial colonization of beet occurs in discrete phases regardless of bioinoculation with the wild sea beet root community"

**Colonization of beet plants by rhizosphere bacteria takes place in two phases regardless of bioinoculation**

**SM1. Surface sterilization of beet seeds**

Seeds were placed in sterile, deionized water-filled Petri dish and incubated for 1 hour. Coating was removed from commercial seeds using forceps, then the seeds were placed in sterile 50 ml Falcon Tubes (BD) and incubated in 70% EtOH for 5 minutes. Next stages were performed in sterile laminar flow hood: after decanting EtOH, seeds were incubated in 50% commercial bleach (Domestos, Unilever) with delicate mixing for 20 minutes. The seeds were rinsed with sterile, deionized water until bleach smell was not perceptible (at least 15 times). Sterilization efficacy was tested by plating 100 µl of last rinse on an LB plate and incubation at room temperature (RT). The seeds were deemed sterile if no colonies formed after 48 hours.

**SM2. Micropropagation and generation of plants with lowered bacterial load**

Sterilized seeds were germinated on agar-solidified Murashige and Skoog (MS, Duchefa) medium (). When two true leaves developed, plantlets were dissected and meristems were explanted on MS supplemented BAP (Sigma, 500 µg/l) and various combinations of antibiotics (Table SM1). Finally, 300 mg/l cefotaxim (Duchefa) and 25 mg/l vancomycin (Duchefa) were chosen to kill seed-borne bacteria and maintain sterility of material. Three subcultures were performed, shoot tips containing apical meristems were isolated and planted on the abovementioned medium. Then, plantlets were transferred to MS supplemented with 2 mg/l of NAA (Sigma) and cultured until roots were developed. Afterwards, plants were transferred to sterile sand:vermiculite (1:1, v:v) and grown for two weeks under plastic foil, until they developed second pair of true leaves. At this stage plants were fed with 0.5 x Hoagland medium and acclimated by gradual removal of foil. The acclimation step lasted four weeks. Then the plants were re-planted to autoclaved soil and grown for four weeks. During this stage plants were watered with sterile, deionized water.

**SM3. Roots surface sterilization**

Plants were uprooted and loose soil was washed out with plenty of sterile water. Roots were rinsed with 200 ml of 0.5% Tween-20 (using a wash bottle) and then incubated for 2 minutes in 30 ml of 5% NaOCl (POCh) in 50 ml Falcon tubes (BD). Incubated roots were rinsed with sterile water (ca. 600 ml) until bleach smell was not perceptible.

**SM4. DNA extraction**

DNA was isolated with in-house made buffers taken from US patent no. US7459548B2 prepared with molecular biology grade chemicals from Sigma, generally according to Qiagen protocol for DNeasy PowerLyzer PowerSoil kit with minor modifications:

- 100 mg of plant material were used, the material was crushed in liquid nitrogen by squeezing sample bag with gloved fingers

- Bead beating was carried out in reinforced screw-cap 2 ml tubes (Omni) with the use of balls of stainless steel (bearing balls, made of AISI316 steel, ), of two sizes (small: 1.6 mm diameter, large: 2.3 mm), for roots 4 large and 4 small ones were used, for leaves 1 large and 4 small ones, while for soil 600 mg of 1 mm glass beads (Sigma) were used. One cycle of 45 s was used at 4000 rpm for roots, 2500 rpm for leaves and 4000 rpm for soil.

- Centrifugations at points 5, 8 and 10 of the protocol were carried out for 1 min at 14 000 x g

- 50 µl of C6 solution were used to eluate DNA from columns

- DNA concentration was measured with Qubit HS kit

- samples were placed in 96-well plates and diluted to 1 ng/µl with PCR-grade water

**SM5. Bacterial 16S rRNA amplicon libraries preparation**

**Table SM2**. Primers. Universal sequences (M13 and M13R) underlined, a pad boosting melting temperature of M13RB786r over 65°C in bold, x stands for 9 nt MID sequence.

| **Primer** | **Sequence** | **Pair** | **Reference** |
| --- | --- | --- | --- |
| M13B357f | GTT TTC CCA GTC ACG ACC CT A CGG GAG GCA GCA G | M13RB786r | Thiem et al. 2018 |
| M13RB786r | CAG GAA ACA GCT ATG AC**C** **GTC GTC GTC** GGA CCA GGG TAT CTA AWC C | M13B357f | Thiem et al. 2018# |
| P5xM13 | AAT GAT ACG GCG ACC ACC GAG ATC TAC AC x GTT TTC CCA GTC ACG AC | P7xM13R | Thiem et al. 2018 |
| P7xM13R | CAA GCA GAA GAC GGC ATA CGA GAT x CAG GAA ACA GCT ATG AC | P5xM13 | Thiem et al. 2018 |
| mtblF | GTT GAG TGC GCG ATC ATG ACA GGA C-ddC | - | This work |
| mtblR | CGA CTT TCA CTT TCA ACC CGA TTC ACC G-ddC | - | This work |
| chblR | GAT AGT TTC CAC CGC CTG TCC AGG G-ddC | - | This work |
| B335F | CAD ACT CCT ACG GGA GGC | B769R | Dorn-In et al. 2015 |
| B769R | ATC CTG TTT GMT MCC CVC RC | B335F | Dorn-In et al. 2015 |
| ASV8p5f | CTT GCT GTT TTG ACG TTA CCG A | ASV8p2r | This work |
| ASV8p2r | GCT TTC ACA TCC AAC TTA ACA A | ASV8p5f | This work |

\# the pad sequence was erroneously omitted in Thiem et al. 2018

**SM5a.** First PCR round – amplification of V3-4 (357-786) bacterial 16S rRNA fragments

Reactions were carried out on Biorad C1000 Touch thermal cycler in 20 µl volume in 96-well plates (GenoPlast) using the following mixes:

- for soil: 2 ng of template DNA (2 µl), 0.8 mM dNTP mix, 5 pmol of M13B357f and M13RB786ru each, 1 x Taq buffer B (EURx), 1 U of TiTaq (EURx)

- for roots, leaves and inoculant: 2 ng of template DNA (2 µl), 0.8 mM dNTP mix, 5 pmol of M13B357f and M13RB786r each, 50 pmol of chblR (blocking primer for chloroplastic 16S rRNA gene sequence), 150 pmol mtblF and mtblR each (blocking primers for mitochondrial 16S rRNA gene sequence), 1 x Taq buffer B (EURx), 1 U of TiTaq (EURx)

The following reagent was obtained through BEI Resources, NIAID, NIH as part of the Human Microbiome Project: Genomic DNA from Microbial Mock Community B (Staggered, Low Concentration), v5.2L, for 16S rRNA Gene Sequencing, HM-783D. Recommended volume of 1 µl was used as template.

Thermal profile was: 95ºC – 3 min., 25 cycles of: 95ºC – 15 s, 54ºC – 20 s, 72ºC – 25 s and final extension 72ºC – 5 min.

**SM5b.** First round product clean-up and concentration measurement

Clean-up beads were prepared from Sera-mag SpeedBeads according to [dx.doi.org/10.17504/protocols.io.bkguktww](https://dx.doi.org/10.17504/protocols.io.bkguktww)

- 30 µl of beads suspention were added to each well and the plate was shaken on a plate shaker (Bionovo) for 1 min. at 1400 rpm

- plate was incubated for 10 min. at RT

- plate was placed on a magnetic rack (in-house made) and incubated for 10 min.

- supernatant was removed using multichannel pipette

- 100 µl of freshly prepared 80% EtOH were added to each well and the plate was incubated for 2 min. in the magnetic rack

- supernatant was removed

- two above steps were repeated

- pellet was air-dried for 3 min. at RT

- 15 µl of PCR grade water were added to each well

- plate was incubated for 10 min. at RT # at this point the plate can be sealed and frozen until needed

- plate was placed on the magnetic rack and incubated for 5 min.

- 2 µl were transferred from each well to a fresh LightCycler plate (Roche) containing 48 µl of Qubit HS mix (ThermoFisher)

- plate was sealed and placed in LightCycler 480 machine (Roche) and the following program was run: filters: 465 nm excitation, 510 nm detection; acquisition mode: single; 10 min. of hold at 37ºC, measurement of fluorescence

- data were exported and DNA concentration was calculated in LibreOffice Calc using standard curve measured for Qubit HS standards

- samples (in the original plate) were diluted to 1 ng/µl with PCR-grade water

**SM5c.** Second PCR round – adapters and indexes introduction

Reactions were set up in 96-well plates (Genoplast) with epMotion 5075 laboratory robot (Eppendorf) and run on Biorad C1000 Touch thermal cycler (Biorad). Volume was 20 µl.

Reactions mix: 2 ng of template DNA (from first round), 0.8 mM dNTP mix, 10 pmol indexing primers, 1 x Taq buffer B (EURx), 1 U TiTaq (EURx)

Thermal profile: 95ºC – 3 min.; 10 cycles of: 95ºC – 30 s, 54ºC – 15 s, 72ºC – 40 s; final extension 72ºC – 5 min.

**SM5d.** Second round product clean-up and quantification

Clean-up and quantification were carried out as in SM5b.

**SM5f.** Final library preparation and quantification

20 ng (volume depended on concentration) were taken from each well and transferred to an eppendorf tube.

Final pool was purified with Sera-mag beads (50 µl in 1.5 ml eppendorf tube, using 1.5 x volume of beads).

Concentration and purity were measured with NanoDrop ND1000.

1 µl of final pool diluted to 0.5 ng/µl was analyzed with Bioanalyzer DNA HS kit.

Based on Bioanalyzer results the pool was diluted to ~0.1 pM and quantified using KAPA Library Quantification Kit (KK4854) on Roche LightCycler 480 according to the manufacturer protocol.

**SM6. qPCR analysis**

qPCR reactions were performed with FastSTART kit (Roche) on LightCycler 480 instrument in 10 µl consisting of 1 x master mix, 5 pmol of each primer and 1 ng of template. Threshold cycles (C_t_s) were determined in LightCycler software using second derivative method and exported as csv files. The C_t_s were converted to copies per nanogram DNA using standard equation using slope and intercept from a standard curve generated on serial dilutions of a purified amplicon: n = 10^((Ct-intercept)/slope)^. Efficiency was calculated using an equation E = 10^(-1/slope)^. Standard curve fitting was carried out in R using lm() (see SM7p for a script).

**SM6a.** 16S rRNA fragment counts (bacterial load assessment)

Bacterial load assessment was carried out using B335F - B769R primer pair (Dorn-In et al. 2015, Table SM2). The following thermal profile was used: 95°C – 5 min., 45 cycles (roots), 50 cycles (leaves) or 55 cycles (seeds and seedlings) of 95°C – 10 s, 60°C – 20 s, 72°C – 20 s, melting curve – ramp from 60°C to 95°C. Standard curve was generated on *Escherichia coli* DH10B amplicon + 1 ng of mixed leaf and root DNA to account for mitochondrial and plastidic sequences.

**SM6b.** ASV10 counts

Abundance of ASV10 (classified as *Pseudomonas*) was assessed using ASV8p5f – ASV8p2r pair (Table SM2) designed using DECIPHER (see script in SM7m). The following thermal profile was used: 95°C – 5 min., 40 cycles (roots) or 50 cycles (leaves) of, 95°C – 10 s, 61°C – 20 s, 72°C – 20 s, melting curve – ramp from 61°C to 95°C. Standard curve was generated on an amplicon produced using mixed beet roots and leaves DNA as a template.

**SM7. Bioinformatic and statistical analyses**

Computations were performed on a Dell Precision T7600 workstation with 16 physical cores (32 threads) and 128 GB RAM running Ubuntu 20.04. R v.4.1.3 and mothur v.1.44.3 were used.

**SM7a.** ASVs generation

ASVs were generated using dada2 v.1.14 (Callahan et al. 2016), using the following trimming parameters in filterAndTrim: trunclen=c(260,240), maxEE=c(2,4) for soils and roots, for leaves trunclen=c(280,210), maxEE=c(4,6), and for inoculant trunclen=c(290, 230), maxEE=c(5,10). Trimming cutoffs were determined by looking at mean quality graphs generated with the dada2’s plotQualityProfile function. Files generated from negative controls were removed prior to analysis, as they contained single reads or no reads at all.

Samples L769 and L777 were removed from the leaves dataset, as after filtering no reads left. In the roots dataset an additional parameter matchIDs=T had to be used in filterAndTrim.

In R:

```
library(dada2)
path <- '/home/mgoleb/Dokumenty/betamikro_WP5_x' # path should point to a directory where reads from a single sequencing run are placed (i.e. from soils, roots, leaves or inoculant)
fnFs <- sort(list.files(path, pattern="_R1.fastq.gz")) # list of R1 files
fnRs <- sort(list.files(path, pattern="_R2.fastq.gz")) # list of R2 files
sample.names <- sapply(strsplit(fnFs, "_"), `[`, 4)
# Specify the full path to the fnFs and fnRs
fnFs <- file.path(path, fnFs)
fnRs <- file.path(path, fnRs)
filt_path <- file.path(path, "filtered") # Place filtered files in filtered/ subdirectory
filtFs <- file.path(filt_path, paste0(sample.names, "_F_filt.fastq.gz"))
filtRs <- file.path(filt_path, paste0(sample.names, "_R_filt.fastq.gz"))
out <- filterAndTrim(fnFs, filtFs, fnRs, filtRs, truncLen=c(280,210), maxN=0, maxEE=c(4,6), truncQ=2, rm.phix=TRUE, compress=TRUE, multithread=TRUE)# set truncLen, maxEE and matchIDs parameters as required for a given dataset
fnFs <- fnFs[-grep("L769|L777", fnFs) ] # only for leaves
fnRs <- fnRs[-grep("L769|L777", fnRs) ] # only for leaves
filtFs <- filtFs[-grep("L769|L777", filtFs) ] # only for leaves
filtRs <- filtRs[-grep("L769|L777", filtRs) ] # only for leaves
sample.names <- sample.names[ - grep("L769|L777", sample.names) ] # only for leaves
errF <- learnErrors(filtFs, multithread=TRUE)
errR <- learnErrors(filtRs, multithread=TRUE)
derepFs <- derepFastq(filtFs, verbose=TRUE)
derepRs <- derepFastq(filtRs, verbose=TRUE)
# Name the derep-class objects by the sample names
names(derepFs) <- sample.names
names(derepRs) <- sample.names
dadaFs <- dada(derepFs, err=errF, multithread=TRUE)
dadaRs <- dada(derepRs, err=errR, multithread=TRUE)
mergers <- mergePairs(dadaFs, derepFs, dadaRs, derepRs, verbose=TRUE)
seqtab <- makeSequenceTable(mergers)
seqtab.nochim <- removeBimeraDenovo(seqtab, method="consensus", multithread=TRUE, verbose=TRUE)
dim(seqtab.nochim)
sum(seqtab.nochim)/sum(seqtab)
getN <- function(x) sum(getUniques(x))
track <- cbind(out, sapply(dadaFs, getN), sapply(mergers, getN), rowSums(seqtab), rowSums(seqtab.nochim))
colnames(track) <- c("input", "filtered", "denoised", "merged", "tabled", "nonchim")
rownames(track) <- sample.names
taxa <- assignTaxonomy(seqtab.nochim, "~/db/silva/v132/silva_nr_v132_train_set.fa.gz", multithread=T)
seqs1.fasta <- data.frame(seq=colnames(seqtab.nochim), row.names=paste(">ASV", seq(1, dim(seqtab.nochim)[2], 1), "\n", sep=""))
write.table(seqs1.fasta, "seqs.fasta", quote=F, sep="", col.names=F)
seqs1.count_table <- as.data.frame(t(seqtab.nochim))
rownames(seqs1.count_table) <- paste(">ASV", seq(1, dim(seqtab.nochim)[2], 1), "\n", sep="")
seqs1.count_table$total <- rowSums(seqs1.count_table)
seqs1.count_table <- seqs1.count_table[, c(ncol(seqs1.count_table), 1:(ncol(seqs1.count_table)-1))]
seqs1.count_table$Representative_sequence <- paste("ASV", seq(1, dim(seqtab.nochim)[2], 1), sep="")
seqs1.count_table <- seqs1.count_table[, c(ncol(seqs1.count_table), 1:(ncol(seqs1.count_table)-1))]
write.table(seqs1.count_table, "seqs.count_table", quote=F, sep="\t", row.names=F)
saveRDS(seqtab.nochim, 'betamikro_WP5_x.rds') # substitute ‘soils’, ‘roots’, ‘leaves’ or ‘inoculant’ for x
quit()
```

**SM7b.** Joining datasets, alignment and tree construction

Datasets were joined in R using mergeSequenceTables of dada2. The sequences were then exported as Mothur-compatible fasta and count_table files to enable relaxed neighbor joining tree construction in Mothur v. 1.43 using clearcut.

In R:

library(dada2)

seqtab.soil <- readRDS("betamikro_WP5_soils.rds")

seqtab.root <- readRDS("betamikro_WP5_roots.rds")

seqtab.inoculant <- readRDS("betamikro_WP5_inoculant.rds")

rownames(seqtab.inoculant) <- sub(".*-", "", rownames(seqtab.inoculant) ) # there were numbers at the beginning of names, they need to be removed

seqtab.leaves <- readRDS("betamikro_WP5_leaves.rds")

seqtabs <- list(seqtab.soil, seqtab.root, seqtab.inoculant, seqtab.leaves)

seqtab.complete <- mergeSequenceTables(tables=seqtabs, repeats='sum')

seqs.fasta <- data.frame(seq=colnames(seqtab.complete), row.names=paste(">ASV", seq(1, dim(seqtab.complete)[2], 1), "\n", sep=""))

write.table(seqs.fasta, "seqs.fasta", quote=F, sep="", col.names=F)

seqs.count_table <- as.data.frame(t(seqtab.complete))

rownames(seqs.count_table) <- paste(">ASV", seq(1, dim(seqtab.complete)[2], 1), "\n", sep="")

seqs.count_table$total <- rowSums(seqs.count_table)

seqs.count_table <- seqs.count_table[, c(ncol(seqs.count_table), 1:(ncol(seqs.count_table)-1))]

seqs.count_table$Representative_sequence <- paste("ASV", seq(1, dim(seqtab.complete)[2], 1), sep="")

seqs.count_table <- seqs.count_table[, c(ncol(seqs.count_table), 1:(ncol(seqs.count_table)-1))]

write.table(seqs.count_table, "seqs.count_table", quote=F, sep="\t", row.names=F)

quit()

In bash:

mothur "#align.seqs(fasta=seqs.fasta, reference=~/db/silva/v132/silva.nr_v132.align, processors=30);"

mothur "#remove.seqs(fasta=seqs.align, count=seqs.count_table, accnos=seqs.flip.accnos);"

mothur "#screen.seqs(fasta=seqs.pick.align, count=seqs.pick.count_table, start=6430, end=23439, processors=30);"

mothur "#filter.seqs(fasta=seqs.pick.good.align, vertical=T, trump=., processors=30);"

mothur "#unique.seqs(fasta=seqs.pick.good.filter.fasta, count=seqs.pick.good.count_table);"

mothur "#dist.seqs(fasta=seqs.pick.good.filter.unique.fasta, output=lt, processors=30);"

mothur "#clearcut(phylip=seqs.pick.good.filter.unique.phylip.dist);" > clearcut.log 2>&1 &

**SM7c.** Sequence classification

Sequences were classified using naive Bayesian classifier implemented in dada2’s assignTaxonomy with SILVA v. 132 reference.

In R:

library(dada2)

seqtab.complete.taxa <- assignTaxonomy(seqtab.complete, "~/db/silva/v132/silva_nr_v132_train_set.fa.gz", multithread=T)

bacteria.tax <- as.data.frame(seqtab.complete.taxa)

rownames(bacteria.tax) <- paste0("ASV", seq(1, dim(seqtab.complete)[2], 1) )

bacteria <- seqtab.complete

colnames(bacteria) <- paste("ASV", seq(1, dim(seqtab.complete)[2], 1), sep="")

bacteria <- bacteria[ , -union( grep("Mitochondria", bacteria.tax$Family), grep("Chloroplast", bacteria.tax$Order) ) ] # removal of ASVs classified as mitochondria and chloroplasts

bacteria.tax <- bacteria.tax[ rownames(bacteria.tax) %in% colnames(bacteria), ]

saveRDS(bacteria.tax, "betamikro_bacteria.tax.rds")

quit()

**SM7d.** Preprocessing, rarefaction and UniFrac distances computation

Sample data were read in from a csv file, ASV table was rarefied to 900 sequences per sample 100 times and mean values rounded to the nearest integer were used for downstream analyses (Golebiewski et al. 2014).

In R:

```
library(phytools)
library(ape)
library(GUniFrac)
sdata <- read.table("WP5.design.csv", header=T, sep=",")
rownames(sdata) <- sdata$sample
sdata$Sample <- gsub("G|K|L", "", rownames(sdata))
sdata$Material <- substr(rownames(sdata), 1,1)
sdata$Material <- gsub("G", "S", sdata$Material)
sdata$Material <- gsub("K", "R", sdata$Material)
sdata$status <- 'late'
sdata$status[ grep("T0|T1|T2", sdata$Time) ] <- 'early'
sdata$groupTime <- paste0( sdata$Material, sdata$Soil, sdata$Genotype, sdata$Time, sdata$Inoculation )
sdata$groupstatus <- paste0( sdata$Material, sdata$Soil, sdata$Genotype, sdata$status, sdata$Inoculation )
sdata <- sdata[ rownames(sdata) %in% rownames(bacteria),  ]
bacteria <- bacteria[ rownames(bacteria) %in% rownames(sdata), ] # only those samples, for which there are data
sdata <- sdata[ rownames(sdata) %in% rownames(bacteria), ]
bacteria <- bacteria[ sort(rownames(bacteria)), ]
sdata <- sdata[ sort(rownames(sdata)), ]
identical(rownames(bacteria), rownames(sdata)) # must be TRUE
bacteria.tree <- ape::read.tree(file="seqs.pick.good.filter.unique.phylip.tre") # tree generated in mothur, script in tree_generation.bash
bacteria.tree$edge.length[which(is.na(bacteria.tree$edge.length))] <- 0
midpoint.root(bacteria.tree)
is.rooted(bacteria.tree) # must be TRUE
saveRDS(bacteria.tree, "betamikro_bacteria.tree.rds")
bacteria <- bacteria[ , colnames(bacteria) %in% bacteria.tree$tip.label ] # only ASVs that are in the tree
saveRDS(bacteria, "betamikro_bacteria.asvtab.rds")
### This block of code allows avoiding pitfalls of single rarefying (Golebiewski et al. 2014) ######
set.seed(667)
temp <- Rarefy(otu.tab=bacteria, depth=900)$otu.tab.rff
for( i in 1:99 ){ print( i ); temp <- temp + Rarefy(otu.tab=bacteria, depth=900)$otu.tab.rff }
bacteria.rarefied <- round( temp/100 )
bacteria.rarefied <- bacteria.rarefied[ , colSums(bacteria.rarefied) > 0 ] # removal of all-zero columns
####################################################################################################
sdata.rarefied <- sdata[ rownames(sdata) %in% rownames(bacteria.rarefied),  ] # data only for samples that had over 900 reads
bacteria.rarefied <- bacteria.rarefied[ rownames(bacteria.rarefied) %in% rownames(sdata.rarefied), ] # only those samples, for which there are data
bacteria.rarefied <- bacteria.rarefied[ sort(rownames(bacteria.rarefied)), ]
sdata.rarefied <- sdata.rarefied[ sort(rownames(sdata.rarefied)), ]
identical(rownames(bacteria.rarefied), rownames(sdata.rarefied)) # must be TRUE

sdata.rarefied$TI <- paste0(sdata.rarefied$Time, '-', sdata.rarefied$Inoculation)

bacteria.unifracs <- GUniFrac( bacteria.rarefied, bacteria.tree, alpha=c(0, 0.5))$unifracs
bacteria.rarefied.uunifrac <- bacteria.unifracs[, , "d_UW"]
bacteria.rarefied.d05unifrac <- bacteria.unifracs[, , "d_0.5"]

quit()
```

**SM7e**. PICRUSt2 analysis

Prediction of metabolic potential was carried out with PICRUSt2 on whole data (without rarefying) for finding differentially represented functions and pathways and on rarefied data for alpha and beta-divesity analyses.

In R:

```
# Preparation of rarefied data for alpha and beta-diversity analyses
seqs.rarefied.count_table <- as.data.frame(t(bacteria.rarefied))
rownames(seqs.rarefied.count_table) <- paste0("Otu", sprintf("%05d", as.numeric(sub("ASV", "", rownames(seqs.rarefied.count_table)))))
seqs.rarefied.count_table$total <- rowSums(seqs.rarefied.count_table)
seqs.rarefied.count_table <- seqs.rarefied.count_table[, c(ncol(seqs.rarefied.count_table), 1:(ncol(seqs.rarefied.count_table)-1))]
seqs.rarefied.count_table$Representative_sequence <- rownames(seqs.rarefied.count_table)
seqs.rarefied.count_table <- seqs.rarefied.count_table[, c(ncol(seqs.rarefied.count_table), 1:(ncol(seqs.rarefied.count_table)-1))]
seqs.rarefied.count_table <- seqs.rarefied.count_table[ seqs.rarefied.count_table$total > 0, ]
write.table(seqs.rarefied.count_table, "seqs.rarefied.count_table", quote=F, sep="\t", row.names=F)

seqs.rarefied.fasta <- seqs1.fasta
seqs.rarefied.fasta$name <- sub("\n", "", rownames(seqs.rarefied.fasta))
seqs.rarefied.fasta$name <- sub(">", "", seqs.rarefied.fasta$name)
rownames(seqs.rarefied.fasta) <- paste0(">Otu", sprintf("%05d", as.numeric(sub("ASV", "", seqs.rarefied.fasta$name))), "\n")
seqs.rarefied.fasta$name <- sub("\n", "", sub(">", "", rownames(seqs.rarefied.fasta)))
seqs.rarefied.fasta <- seqs.rarefied.fasta[ seqs.rarefied.fasta$name %in% seqs.rarefied.count_table$Representative_sequence, ]
seqs.rarefied.fasta$name <- NULL
write.table(seqs.rarefied.fasta, "seqs.rarefied.fasta", quote=F, sep="", col.names=F)
```

In bash:

```
mothur "#make.shared(count=seqs.rarefied.count_table)" # seqs.rarefied.count_table – data rarefied data

mothur "#make.biom(shared=seqs.rarefied.asv.shared)"

picrust2_pipeline.py -s seqs.rarefied.fasta -i seqs.rarefied.asv.asv.biom -o picrust2_out -p 16 > picrust2.log 2>&1 &

add_descriptions.py -i EC_metagenome_out/pred_metagenome_unstrat.tsv.gz -m EC \
                    -o EC_metagenome_out/pred_metagenome_unstrat_descrip.tsv.gz

add_descriptions.py -i KO_metagenome_out/pred_metagenome_unstrat.tsv.gz -m KO \
                    -o KO_metagenome_out/pred_metagenome_unstrat_descrip.tsv.gz

add_descriptions.py -i pathways_out/path_abun_unstrat.tsv.gz -m METACYC \
                    -o pathways_out/path_abun_unstrat_descrip.tsv.gz

mv picrust2_out rarefied_picrust2_out

# Preparation of non-rarefied data for differentially abundant features identification
```

In bash:

```
rename_asvs_to_otus.perl -f seqs1.fasta -n Otu

mothur "#make.shared(count=seqs1.count_table)" # seqs1.count_table – data without inoculant

mothur "#make.biom(shared=seqs1.asv.shared)"

picrust2_pipeline.py -s seqs1.fasta -i seqs1.asv.asv.biom -o picrust2_out -p 16 > picrust2.log 2>&1 &

add_descriptions.py -i EC_metagenome_out/pred_metagenome_unstrat.tsv.gz -m EC \
                    -o EC_metagenome_out/pred_metagenome_unstrat_descrip.tsv.gz

add_descriptions.py -i KO_metagenome_out/pred_metagenome_unstrat.tsv.gz -m KO \
                    -o KO_metagenome_out/pred_metagenome_unstrat_descrip.tsv.gz

add_descriptions.py -i pathways_out/path_abun_unstrat.tsv.gz -m METACYC \
                    -o pathways_out/path_abun_unstrat_descrip.tsv.gz

mv picrust2_out nonrarefied_picrust2_out
```

**SM7f.** Alpha diversity calculation and boxplots

In R:

library(vegan)

library(ggplot2)

\# amplicon data

bacteria.rarefied.diversity <- data.frame( row.names=rownames(bacteria.rarefied), sobs=specnumber(bacteria.rarefied), H=diversity(bacteria.rarefied), E=diversity(bacteria.rarefied)/log(specnumber(bacteria.rarefied)), material=sdata.rarefied$Organ, soil=sdata.rarefied$Soil, genotype=sdata.rarefied$Genotype, time=sdata.rarefied$Time, status=sdata.rarefied$status, inoculation=sdata.rarefied$Inoculation)

svg( "bacteria.rarefied.diversity.H.material.svg", width=1.75, height=1.75, pointsize=3 ) # Fig. 2B

p <- ggplot( bacteria.rarefied.diversity, aes_(x=bacteria.rarefied.diversity$material, y=bacteria.rarefied.diversity$H) ) + geom_boxplot() + scale_y_continuous() + xlab("Material") + ylab("Shannon’s H’") + ggtitle("Shannon’s H’")

print(p)

dev.off()

svg( "bacteria.rarefied.diversity.E.material.svg", width=1.75, height=1.75, pointsize=3 ) # Fig. 2B

p <- ggplot( bacteria.rarefied.diversity, aes_(x=bacteria.rarefied.diversity$material, y=bacteria.rarefied.diversity$E) ) + geom_boxplot() + scale_y_continuous() + xlab("Material") + ylab("Shannon’s E") + ggtitle("Shannon’s E")

print(p)

dev.off()

svg( "bacteria.rarefied.diversity.sobs.material.svg", width=1.75, height=1.75, pointsize=3 ) # Fig. 2B

p <- ggplot( bacteria.rarefied.diversity, aes_(x=bacteria.rarefied.diversity$material, y=bacteria.rarefied.diversity$sobs) ) + geom_boxplot() + scale_y_continuous() + xlab("Material") + ylab("Obs. no. ASVs") + ggtitle("Richness")

print(p)

dev.off()

for( index in c('H', 'E', 'sobs') ){

if( index == 'H' ){

title = "Shannon's H'";

ylabel <- "Shannon's H'";

} else if( index == "E" ){

title = "Shannon's E";

ylabel <- "Shannon's E";

} else {

title = "Richness";

ylabel <- "No. obs. ASVs";

}

for( material in c('S', 'R', 'L') ){

chunk <- paste0('bacteria.rarefied.diversity.', material );

file1 <- paste0('bacteria.rarefied.diversity', '.', material, '.', index, '.ggplot.svg'); # Fig. SR4

file4 <- paste0('bacteria.rarefied.diversity', '.', material, '.', index, '.no_facetting.ggplot.svg'); # Fig. 4B,D,F

assign(chunk, bacteria.rarefied.diversity[ bacteria.rarefied.diversity$material == material, ]);

svg( file1, width=7, height=3.5, pointsize=6 );

p <- ggplot( get(chunk), aes_(x=get(chunk)$time, y=get(chunk)[[index]], fill=get(chunk)$inoculation) ) + geom_boxplot() + facet_wrap( ~ get(chunk)$soil * get(chunk)$genotype ) + scale_x_discrete(drop=T) + xlab("Timepoint") + ylab(ylabel) + labs(fill="Inoculation") + ggtitle(title);

print( p );

dev.off();

svg( file4, width=1.75, height=1.75, pointsize=3 );

p <- ggplot( get(chunk), aes_(x=get(chunk)$time, y=get(chunk)[[index]]) ) + geom_boxplot() + scale_x_discrete(drop=T) + xlab("Timepoint") + ylab(ylabel) + ggtitle(title);

print( p );

dev.off();

}

file2 <- paste0('bacteria.rarefied.diversity_time.', index, '.ggplot.svg'); # Fig. SR

file3 <- paste0('bacteria.rarefied.diversity_status.', index, '.ggplot.svg'); # Fig. SR

svg( file2, width=7, height=10.5, pointsize=3 );

p <- ggplot(bacteria.rarefied.diversity, aes_(x=bacteria.rarefied.diversity$time, y=bacteria.rarefied.diversity[[index]], fill=bacteria.rarefied.diversity$inoculation)) + geom_boxplot() + facet_wrap( ~ bacteria.rarefied.diversity$material * bacteria.rarefied.diversity$soil * bacteria.rarefied.diversity$genotype, ncol=3 ) + scale_x_discrete(drop=T) + xlab("Timepoint") + ylab(ylabel) + labs(fill="Inoculation")

print(p);

dev.off();

svg( file3, width=7, height=10.5, pointsize=3 );

p <- ggplot(bacteria.rarefied.diversity, aes_(x=bacteria.rarefied.diversity$genotype, y=bacteria.rarefied.diversity[[index]], fill=bacteria.rarefied.diversity$inoculation)) + geom_boxplot() + facet_wrap( ~ bacteria.rarefied.diversity$material * bacteria.rarefied.diversity$soil * bacteria.rarefied.diversity$status, ncol=4 ) + scale_x_discrete(drop=T) + xlab("Genotype") + ylab(ylabel) + labs(fill="Inoculation")

print(p);

dev.off();

}

\# PICRUSt2 simulated metabolic potential at the level of functions

\# Generating Shannon's H', Shannon's E and observed number of ASVs

picrust_ko.diversity <- data.frame( row.names=rownames(picrust_ko), sobs=specnumber(picrust_ko), H=diversity(picrust_ko), E=diversity(picrust_ko)/log(specnumber(picrust_ko)), material=sdata.rarefied$Material, soil=sdata.rarefied$Soil, genotype=sdata.rarefied$Genotype, time=sdata.rarefied$Time, status=sdata.rarefied$status, inoculation=sdata.rarefied$Inoculation)

picrust_ko.diversity <- picrust_ko.diversity[ -grep("INOK", rownames(picrust_ko.diversity)), ] # no need for inoculant here

svg( "picrust_ko.diversity.H.material.svg", width=1.75, height=1.75, pointsize=3 ) # Fig. 2D

p <- ggplot( picrust_ko.diversity, aes_(x=picrust_ko.diversity$material, y=picrust_ko.diversity$H) ) + geom_boxplot() + scale_y_continuous() + xlab("Material") + ylab("Shannon’s H’") + ggtitle("Shannon’s H’")

print(p)

dev.off()

svg( "picrust_ko.diversity.E.material.svg", width=1.75, height=1.75, pointsize=3 ) # Fig. 2D

p <- ggplot( picrust_ko.diversity, aes_(x=picrust_ko.diversity$material, y=picrust_ko.diversity$E) ) + geom_boxplot() + scale_y_continuous() + xlab("Material") + ylab("Shannon’s E") + ggtitle("Shannon’s E")

print(p)

dev.off()

svg( "picrust_ko.diversity.sobs.material.svg", width=1.75, height=1.75, pointsize=3 ) # Fig. 2D

p <- ggplot( picrust_ko.diversity, aes_(x=picrust_ko.diversity$material, y=picrust_ko.diversity$sobs) ) + geom_boxplot() + scale_y_continuous() + xlab("Material") + ylab("Obs. no. functions") + ggtitle("Richness")

print(p)

dev.off()

for( index in c('H', 'E', 'sobs') ){

if( index == 'H' ){

title = "Shannon's H'";

ylabel <- "Shannon's H'";

} else if( index == "E" ){

title = "Shannon's E";

ylabel <- "Shannon's E";

} else {

title = "Richness";

ylabel <- "No. obs. functions";

}

for( material in c('S', 'R', 'L') ){

chunk <- paste0('picrust_ko.diversity.', material );

file1 <- paste0('picrust_ko.diversity', '.', material, '.', index, '.ggplot.svg'); # Fig. SR4

file4 <- paste0('picrust_ko.diversity', '.', material, '.', index, '.no_facetting.ggplot.svg'); # Fig. 3B,D,F

assign(chunk, picrust_ko.diversity[ picrust_ko.diversity$material == material, ]);

svg( file1, width=7, height=3.5, pointsize=6 );

p <- ggplot( get(chunk), aes_(x=get(chunk)$time, y=get(chunk)[[index]], fill=get(chunk)$inoculation) ) + geom_boxplot() + facet_wrap( ~ get(chunk)$soil * get(chunk)$genotype ) + scale_x_discrete(drop=T) + xlab("Timepoint") + ylab(ylabel) + labs(fill="Inoculation") + ggtitle(title);

print( p );

dev.off();

svg( file4, width=1.75, height=1.75, pointsize=3 );

p <- ggplot( get(chunk), aes_(x=get(chunk)$time, y=get(chunk)[[index]]) ) + geom_boxplot() + scale_x_discrete(drop=T) + xlab("Timepoint") + ylab(ylabel) + ggtitle(title);

print( p );

dev.off();

}

file2 <- paste0('picrust_ko.diversity_time.', index, '.ggplot.svg'); # Fig. SR

file3 <- paste0('picrust_ko.diversity_status.', index, '.ggplot.svg'); # Fig. SR

svg( file2, width=7, height=10.5, pointsize=3 );

p <- ggplot(picrust_ko.diversity, aes_(x=picrust_ko.diversity$time, y=picrust_ko.diversity[[index]], fill=picrust_ko.diversity$inoculation)) + geom_boxplot() + facet_wrap( ~ picrust_ko.diversity$material * picrust_ko.diversity$soil * picrust_ko.diversity$genotype, ncol=3 ) + scale_x_discrete(drop=T) + xlab("Timepoint") + ylab(ylabel) + labs(fill="Inoculation")

print(p);

dev.off();

svg( file3, width=7, height=10.5, pointsize=3 );

p <- ggplot(picrust_ko.diversity, aes_(x=picrust_ko.diversity$genotype, y=picrust_ko.diversity[[index]], fill=picrust_ko.diversity$inoculation)) + geom_boxplot() + facet_wrap( ~ picrust_ko.diversity$material * picrust_ko.diversity$soil * picrust_ko.diversity$status, ncol=4 ) + scale_x_discrete(drop=T) + xlab("Genotype") + ylab(ylabel) + labs(fill="Inoculation")

print(p);

dev.off();

}

for( material in c('L', 'R', 'S') ){

for( meas in c('H', 'E', 'sobs') ){

chunk <- paste0( 'picrust_ko.diversity.', material );

wilctest <- paste0( chunk, '.', meas, '.status.wilcox');

assign( chunk, picrust_ko.diversity[ picrust_ko.diversity$material == material, ] );

assign( wilctest, wilcox.test(as.formula(paste(meas, "~ status")), data=get(chunk)) );

}

}

**SM7g.** Ordination plots and testing significance of grouping

```
library(vegan)
library(PairwiseAdonis)

### amplicon data
# whole data – showing clustering according to material and status (first time)

## Fig. 2A
bacteria.rarefied.d05unifrac.nmds <- metaMDS(bacteria.rarefied.d05unifrac, k=2, try=100, trymax=500)
bacteria.rarefied.d05unifrac.Material.adonis <- adonis2( as.dist(bacteria.rarefied.d05unifrac) ~ Material, data=sdata.rarefied, permu=999 )
bacteria.rarefied.d05unifrac.Material.betadisper <- betadisper( as.dist(bacteria.rarefied.d05unifrac), sdata.rarefied$Material, bias.adjust=T )
bacteria.rarefied.d05unifrac.Material.betadisper.anova <- anova( bacteria.rarefied.d05unifrac.Material.betadisper )

svg( "bacteria.rarefied.d05unifrac.nmds.svg", width=3.5, height=3.5, pointsize=6); # Fig. 2A
par(mar=c(4,4,2,4)+0.1, mfrow=c(1,1), las=2);
ordiplot(bacteria.rarefied.d05unifrac.nmds, type='none', main="");
points( bacteria.rarefied.d05unifrac.nmds, pch=19+as.numeric(as.factor(sdata.rarefied$Time)), col='black', bg=as.numeric(as.factor(sdata.rarefied$Material)) );
legend('topright', legend=levels(as.factor(sdata.rarefied$Time)), pch=19+sort(unique(as.numeric(as.factor(sdata.rarefied$Time)))));
legend('topleft', legend=levels(as.factor(sdata.rarefied$Material)), pch=21, pt.bg=sort(unique(as.numeric(as.factor(sdata.rarefied$Material)))));
dev.off();

## Fig. 3B
# dividing according to material, Fig. 4A (leaves), C (roots), E (soils)
for( material in c("S", "R", "L") ){
	sampledata <- paste0("sdata.rarefied.", material);
	d05unifrac <- paste0('bacteria.rarefied.', material, '.', 'd05unifrac');
	ord <- paste0('bacteria.rarefied.', material, '.d05unifrac.nmds')
	dbrdaTime <- paste0('bacteria.rarefied.', material, '.', 'd05unifrac.Time.dbrda');
	dbrdaStatus <- paste0('bacteria.rarefied.', material, '.', 'd05unifrac.status.dbrda');
	adonisTime <- paste0('bacteria.rarefied.', material, '.', 'd05unifrac.Time.adonis');
	betadisperTime <- paste0('bacteria.rarefied.', material, '.d05unifrac.Time.betadisper');
	betadisperTimeAnova <- paste0('bacteria.rarefied.', material, '.d05unifrac.Time.betadisper.anova');
	adonisStatus <- paste0('bacteria.rarefied.', material, '.', 'd05unifrac.status.adonis');
	betadisperStatus <- paste0('bacteria.rarefied.', material, '.d05unifrac.status.betadisper');
	betadisperStatusAnova <- paste0('bacteria.rarefied.', material, '.d05unifrac.status.betadisper.anova');
	dbrdaanovaTime <- paste0('bacteria.rarefied.', material, '.', 'd05unifrac.Time.dbrda.anova');
	dbrdaanovaStatus <- paste0('bacteria.rarefied.', material, '.', 'd05unifrac.status.dbrda.anova');
	filedbrdaTime <- paste0('bacteria.rarefied.', material, '.', 'd05unifrac.Time.dbrda.svg');
	filedbrdaStatus <- paste0('bacteria.rarefied.', material, '.d05unifrac.status.dbrda.svg');
	filenmds <- paste0('bacteria.rarefied.', material, '.d05unifrac.nmds.svg');
	assign( otutable, bacteria.rarefied[grep(string, sdata.rarefied$Material), ] );
	assign( sampledata, sdata.rarefied[grep(string, sdata.rarefied$Material), ] );
	assign( d05unifrac, bacteria.rarefied.d05unifrac[ sdata.rarefied$Material == material, sdata.rarefied$Material == material ] );
	assign( ord, metaMDS(get(d05unifrac), k=2, try=100, trymax=500) );
	assign( dbrdaTime, dbrda(get(d05unifrac) ~ get(sampledata)$Time) );
	assign( dbrdaanovaTime, anova(get(dbrdaTime)) );
	assign( dbrdaStatus, dbrda(get(d05unifrac) ~ get(sampledata)$status) );
	assign( dbrdaanovaStatus, anova(get(dbrdaStatus)) );
	assign( adonisTime, adonis2(get(d05unifrac) ~ Time, data=get(sampledata), permu=999) );
	assign( betadisperTime, betadisper(as.dist(get(d05unifrac)), get(sampledata)$Time, bias.adjust=T) );
	assign( betadisperTimeAnova, anova(get(betadisperTime)) );
	assign( adonisStatus, adonis2(get(d05unifrac) ~ status, data=get(sampledata), permu=999) );
	assign( betadisperStatus, betadisper(as.dist(get(d05unifrac)), get(sampledata)$status, bias.adjust=T) );
	assign( betadisperStatusAnova, anova(get(betadisperStatus)) );
	# dbRDA for Time, points show Time, ellipses show status
	svg( filedbrdaTime, width=3.5, height=3.5, pointsize=6);
	par(mar=c(4,4,2,4)+0.1, mfrow=c(1,1), las=2);
	ordiplot(get(dbrdaTime), type='none');
	points( get(dbrdaTime), pch=20+as.numeric(as.factor(get(sampledata)$Time)), col='black', bg=as.numeric(as.factor(get(sampledata)$status)) );
	ordiellipse( get(dbrdaTime), groups=get(sampledata)$status, label=T);
	legend('topright', legend=levels(as.factor(get(sampledata)$Time)), pch=20+sort(unique(as.numeric(as.factor(get(sampledata)$Time)))));
	legend('topleft', legend=levels(as.factor(get(sampledata)$status)), pch=21, pt.bg=sort(unique(as.numeric(as.factor(get(sampledata)$status)))));
	dev.off();
	# nMDS, points show Time, ellipses show status
	svg( filenmds, width=3.5, height=3.5, pointsize=6);
	par(mar=c(4,4,2,4)+0.1, mfrow=c(1,1), las=2);
	ordiplot(get(ord), type='none');
	points( get(ord), pch=20+as.numeric(as.factor(get(sampledata)$Time)), col='black', bg=as.numeric(as.factor(get(sampledata)$status)) );
	ordiellipse( get(ord), groups=get(sampledata)$status, label=T);
	legend('topright', legend=levels(as.factor(get(sampledata)$Time)), pch=20+sort(unique(as.numeric(as.factor(get(sampledata)$Time)))));
	legend('topleft', legend=levels(as.factor(get(sampledata)$status)), pch=21, pt.bg=sort(unique(as.numeric(as.factor(get(sampledata)$status)))));
	dev.off();
}

# dividing according to material * soil * genotype, checking for significance of status - Fig. SR4, 5, 6 and of Inoculation - Figs. SR9 (leaves), 10 (roots) and 11 (soils)
for( material in c('L', "R", "S") ){
	for( soil in c('S1', 'S2') ){
		for( genotype in c('B', 'C', 'M') ){
			d05unifrac <- paste0( 'bacteria.rarefied.d05unifrac.', material, '.', soil, '.', genotype );
			sampledata <- paste0( 'sdata.rarefied.', material, '.', soil, '.', genotype );
			print( d05unifrac );
			if( nrow(get(sampledata)[ get(sampledata)$status == 'early', ]) > 2 & nrow(get(sampledata)[ get(sampledata)$status == 'late', ]) > 2 ){ # no statistics on 1 case...
				ordvarpart <- paste0(d05unifrac, '.varpart'); # needs to be computed beforehand

				ordnmds <- paste0(d05unifrac, '.nmds');
				ordadonis <- paste0(d05unifrac, '.adonis');
				ordadonis1 <- paste0(d05unifrac, '.status_x_inoculation.adonis');
				ordbetadisper <- paste0(d05unifrac, '.betadisper');
				betadisperanova <- paste0(ordbetadisper, '.anova');
				orddbrda <- paste0(d05unifrac, '.dbrda');
				orddbrda1 <- paste0(d05unifrac, '.status_x_inoculation.dbrda');
				orddbrda2 <- paste0(d05unifrac, '.inoculation.dbrda');
				dbrdaanova <- paste0(orddbrda, '.anova');
				dbrdaanova1 <- paste0(orddbrda1, '.anova');
				dbrdaanova2 <- paste0(orddbrda2, '.anova');
				filenmds <- paste0(ordnmds, '.svg');
				filedbrda <- paste0(orddbrda, '.svg');
				filedbrda1 <- paste0(orddbrda1, '.svg');
				assign( d05unifrac, as.dist(bacteria.rarefied.d05unifrac[ sdata.rarefied$Material == material & sdata.rarefied$Soil == soil & sdata.rarefied$Genotype == genotype, sdata.rarefied$Material == material & sdata.rarefied$Soil == soil & sdata.rarefied$Genotype == genotype ]) );
				assign( sampledata, sdata.rarefied[ sdata.rarefied$Material == material & sdata.rarefied$Soil == soil & sdata.rarefied$Genotype == genotype,  ] );
				assign( ordnmds, metaMDS(as.dist(get(d05unifrac)), k=2, try=100, trymax=500) );
				assign( ordadonis, adonis2(as.dist(get(d05unifrac)) ~ status, data=get(sampledata), permu=999) );
				assign( ordbetadisper, betadisper(get(d05unifrac), get(sampledata)$status, bias.adjust=T) );
				assign( betadisperanova, anova(get(ordbetadisper), permu=999) );
				assign( orddbrda, dbrda(get(d05unifrac) ~ status, data=get(sampledata)) );
				assign( dbrdaanova, anova(get(orddbrda), permu=999) );
				assign( orddbrda1, dbrda(get(d05unifrac) ~ status * Inoculation, data=get(sampledata)) );
				assign( dbrdaanova1, anova(get(orddbrda1), permu=999) );
				assign( orddbrda2, dbrda(get(d05unifrac) ~ Inoculation + Condition(status), data=get(sampledata)) );
				assign( dbrdaanova2, anova(get(orddbrda2), permu=999) );
				explvar_status <- paste0(sprintf("%.2f", 100 * get(ordvarpart)$part$indfract[1,3]), '%');
				explvar_inoculation <- paste0(sprintf("%.2f", 100 * get(ordvarpart)$part$indfract[3,3]), '%');
				permanovaf <- sprintf( "%.2f", get(ordadonis)$F[1] );
				permanovap <- sprintf( "%.3f", get(ordadonis)$"Pr(>F)"[1] );
				betadisperf <- sprintf( "%.2f", get(betadisperanova)$"F value"[1] );
				betadisperp <- sprintf( "%.2e", get(betadisperanova)$"Pr(>F)"[1] );
				dbrdaanovaf <- sprintf( "%.2f", get(dbrdaanova)$F[1] );
				dbrdaanovap <- sprintf( "%.3f", get(dbrdaanova)$"Pr(>F)"[1] );
				dbrdaanova2f <- sprintf( "%.2f", get(dbrdaanova2)$F[1] );
				dbrdaanova2p <- sprintf( "%.3f", get(dbrdaanova2)$"Pr(>F)"[1] );

				title <- paste0( "PERMANOVA F = ", permanovaf, ", p = ", permanovap, "\nbetadisper F = ", betadisperf, ", p = ", betadisperp, "\nVariance explained: ", explvar_status );
				svg( filenmds, width=3.5, height=3.5, pointsize=6 );
				par(mar=c(4,4,4,2)+0.1, mfrow=c(1,1), las=2);
				ordiplot(get(ordnmds), type='none', main=title);
				points( get(ordnmds), pch=20+as.numeric(as.factor(get(sampledata)$Time)), col='black', bg=as.numeric(as.factor(get(sampledata)$status)) );
				ordiellipse( get(ordnmds), groups=get(sampledata)$status, label=T);
				legend('topright', legend=levels(as.factor(get(sampledata)$Time)), pch=20+sort(unique(as.numeric(as.factor(get(sampledata)$Time)))));
				legend('topleft', legend=levels(as.factor(get(sampledata)$status)), pch=21, pt.bg=sort(unique(as.numeric(as.factor(get(sampledata)$status)))));
				dev.off();

				title <- paste0("ANOVA F = ", dbrdaanovaf, ", p = ", dbrdaanovap, "\nbetadisper F = ", betadisperf, ", p = ", betadisperp, "\nVariance explained: ", explvar_status ); # Fig. SR4
				svg( filedbrda, width=3.5, height=3.5, pointsize=6 );
				par(mar=c(4,4,4,2)+0.1, mfrow=c(1,1), las=2);
				ordiplot(get(orddbrda), type='none', main=title);
				points( get(orddbrda), pch=20+as.numeric(as.factor(get(sampledata)$Time)), col='black', bg=as.numeric(as.factor(get(sampledata)$status)) );
				ordiellipse( get(orddbrda), groups=get(sampledata)$status, label=T);
				legend('topright', legend=levels(as.factor(get(sampledata)$Time)), pch=20+sort(unique(as.numeric(as.factor(get(sampledata)$Time)))));
				legend('topleft', legend=levels(as.factor(get(sampledata)$status)), pch=21, pt.bg=sort(unique(as.numeric(as.factor(get(sampledata)$status)))));
				dev.off();

				title <- paste0("ANOVA F = ", dbrdaanova2f, ", p = ", dbrdaanova2p, "\nVariance explained: status - ", explvar_status, ', inoculation - ', explvar_inoculation ); # Figs. SR9, 10, 11
				svg( filedbrda1, width=3.5, height=3.5, pointsize=6 );
				par(mar=c(4,4,4,2)+0.1, mfrow=c(1,1), las=2);
				ordiplot(get(orddbrda1), type='none', main=title);
				points( get(orddbrda1), pch=20+as.numeric(as.factor(get(sampledata)$Inoculation)), col='black', bg=as.numeric(as.factor(get(sampledata)$status)) );
				ordiellipse( get(orddbrda1), groups=get(sampledata)$status, label=T);
				legend('topright', legend=levels(as.factor(get(sampledata)$Inoculation)), pch=20+sort(unique(as.numeric(as.factor(get(sampledata)$Inoculation)))));
				legend('topleft', legend=levels(as.factor(get(sampledata)$status)), pch=21, pt.bg=sort(unique(as.numeric(as.factor(get(sampledata)$status)))));
				dev.off();
				row <- c( material, soil, genotype, get(ordvarpart)$part$indfract$Adj.R.squared[1], get(ordvarpart)$part$indfract$Adj.R.squared[3], get(ordanova_status)$"Pr(>F)"[1], get(ordanova_inoc)$"Pr(>F)"[1] );
				print( row );
				bacteria.inoculation_influence.d05unifrac <- rbind( bacteria.inoculation_influence.d05unifrac, row );


			}
		}
	}
}


### PICRUSt2-predicted functions – using Morisita-Horn distance
# whole data – checking for significance of material
picrust_ko.horn <- vegdist(picrust_ko, method='horn')
picrust_ko.horn.Material.adonis <- adonis2( as.dist(picrust_ko.horn) ~ Material, data=sdata.rarefied, permu=999 )
picrust_ko.horn.Material.betadisper <- betadisper( as.dist(picrust_ko.horn), sdata.rarefied$Material, bias.adjust=T )
picrust_ko.horn.Material.betadisper.anova <- anova( picrust_ko.horn.Material.betadisper )

picrust_ko.horn.Time.adonis <- adonis2( as.dist(picrust_ko.horn) ~ Time, data=sdata.rarefied, permu=999 )
picrust_ko.horn.Time.betadisper <- betadisper( as.dist(picrust_ko.horn), sdata.rarefied$Time, bias.adjust=T )
picrust_ko.horn.Time.betadisper.anova <- anova( picrust_ko.horn.Time.betadisper )

picrust_ko.horn.Material.dbrda <- dbrda(as.dist(picrust_ko.horn) ~ Material, data=sdata.rarefied)

svg( "picrust_ko.horn.Material.dbrda.svg", width=3.5, height=3.5, pointsize=6); # Fig. 3A
par(mar=c(4,4,2,4)+0.1, mfrow=c(1,1), las=2);
ordiplot(picrust_ko.horn.Material.dbrda, type='none', main="");
points( picrust_ko.horn.Material.dbrda, pch=19+as.numeric(as.factor(sdata.rarefied$Time)), col='black', bg=as.numeric(as.factor(sdata.rarefied$Material)) );
legend('topright', legend=levels(as.factor(sdata.rarefied$Time)), pch=19+sort(unique(as.numeric(as.factor(sdata.rarefied$Time)))));
legend('topleft', legend=levels(as.factor(sdata.rarefied$Material)), pch=21, pt.bg=sort(unique(as.numeric(as.factor(sdata.rarefied$Material)))));
dev.off();


# data divided according to material – checking for significance of status
for( material in c("S", "R", "L") ){
	sampledata <- paste0("sdata.rarefied.", material);
	otutable <- paste0("picrust_ko.", material);
	dhorn <- paste0('picrust_ko.', material, '.', 'horn');
	ord <- paste0('picrust_ko.', material, '.horn.nmds')
	vpart <- paste0(dhorn, '.varpart');
	dbrdaTime <- paste0('picrust_ko.', material, '.', 'horn.Time.dbrda');
	dbrdaStatus <- paste0('picrust_ko.', material, '.', 'horn.status.dbrda');
	adonisTime <- paste0('picrust_ko.', material, '.', 'horn.Time.adonis');
	betadisperTime <- paste0('picrust_ko.', material, '.horn.Time.betadisper');
	betadisperTimeAnova <- paste0('picrust_ko.', material, '.horn.Time.betadisper.anova');
	adonisStatus <- paste0('picrust_ko.', material, '.', 'horn.status.adonis');
	betadisperStatus <- paste0('picrust_ko.', material, '.horn.status.betadisper');
	betadisperStatusAnova <- paste0('picrust_ko.', material, '.horn.status.betadisper.anova');
	dbrdaanovaTime <- paste0('picrust_ko.', material, '.', 'horn.Time.dbrda.anova');
	dbrdaanovaStatus <- paste0('picrust_ko.', material, '.', 'horn.status.dbrda.anova');
	filedbrdaTime <- paste0('picrust_ko.', material, '.', 'horn.Time.dbrda.svg');
	filedbrdaStatus <- paste0('picrust_ko.', material, '.horn.status.dbrda.svg');
	filenmds <- paste0('picrust_ko.', material, '.horn.nmds.svg');
	assign( otutable, picrust_ko[ sdata.rarefied$Material == material, ] );
	assign( sampledata, sdata.rarefied[grep(material, sdata.rarefied$Material), ] );
	assign( dhorn, vegdist(get(otutable), dist='horn') );
	assign( vpart, varpart(get(dhorn), ~ status, ~ Soil, ~ Genotype, ~ Inoculation, data=get(sampledata)) );
	assign( ord, metaMDS(get(dhorn), k=2, try=100, trymax=500) );
	assign( dbrdaTime, dbrda(get(dhorn) ~ get(sampledata)$Time) );
	assign( dbrdaanovaTime, anova(get(dbrdaTime), permu=999) );
	assign( dbrdaStatus, dbrda(get(dhorn) ~ status, data=get(sampledata)) );
	assign( dbrdaanovaStatus, anova(get(dbrdaStatus)) );
	assign( adonisTime, adonis2(get(dhorn) ~ Time, data=get(sampledata), permu=999) );
	assign( betadisperTime, betadisper(get(dhorn), get(sampledata)$Time, bias.adjust=T) );
	assign( betadisperTimeAnova, anova(get(betadisperTime), permu=999) );
	assign( adonisStatus, adonis2(get(dhorn) ~ status, data=get(sampledata), permu=999) );
	assign( betadisperStatus, betadisper(get(dhorn), get(sampledata)$status, bias.adjust=T) );
	assign( betadisperStatusAnova, anova(get(betadisperStatus), permu=999) );
	# dbRDA for Time, points show Time, ellipses show status
	svg( filedbrdaTime, width=3.5, height=3.5, pointsize=6);
	par(mar=c(4,4,2,4)+0.1, mfrow=c(1,1), las=2);
	ordiplot(get(dbrdaTime), type='none');
	points( get(dbrdaTime), pch=20+as.numeric(as.factor(get(sampledata)$Time)), col='black', bg=as.numeric(as.factor(get(sampledata)$status)) );
	ordiellipse( get(dbrdaTime), groups=get(sampledata)$status, label=T);
	legend('topright', legend=levels(as.factor(get(sampledata)$Time)), pch=20+sort(unique(as.numeric(as.factor(get(sampledata)$Time)))));
	legend('topleft', legend=levels(as.factor(get(sampledata)$status)), pch=21, pt.bg=sort(unique(as.numeric(as.factor(get(sampledata)$status)))));
	dev.off();
	# nMDS, points show Time, ellipses show status
	svg( filenmds, width=3.5, height=3.5, pointsize=6);
	par(mar=c(4,4,2,4)+0.1, mfrow=c(1,1), las=2);
	ordiplot(get(ord), type='none');
	points( get(ord), pch=20+as.numeric(as.factor(get(sampledata)$Time)), col='black', bg=as.numeric(as.factor(get(sampledata)$status)) );
	ordiellipse( get(ord), groups=get(sampledata)$status, label=T);
	legend('topright', legend=levels(as.factor(get(sampledata)$Time)), pch=20+sort(unique(as.numeric(as.factor(get(sampledata)$Time)))));
	legend('topleft', legend=levels(as.factor(get(sampledata)$status)), pch=21, pt.bg=sort(unique(as.numeric(as.factor(get(sampledata)$status)))));
	dev.off();
}


# data divided according to experimental variant – checking for significance of status and inoculation
picrust_ko.inoculation_influence.horn <- data.frame( matrix(ncol=7, nrow=0) );
for( material in c("S", "R", "L") ){
	otutable <- paste0("picrust_ko.horn.", material);
	assign(otutable, picrust_ko[ sdata.rarefied$Material == material , sdata.rarefied$Material == material  ])
	sampledata <- paste0("sdata.rarefied.", material);
	assign(sampledata, sdata.rarefied[ sdata.rarefied$Material == material, ]);
	distmat <- paste0(otutable, '.horn');
	assign( distmat, vegdist(get(otutable), dist='horn') );
	var_part <- paste0(distmat, ".varpart");
	assign( var_part, varpart(get(distmat), ~ status, ~ Soil, ~ Genotype, ~ Inoculation, data=get(sampledata)) );
	for( soil in c("S1", "S2") ){
		for( genotype in c("B", "C", "M") ){
			otutable <- paste0("picrust_ko.", material, ".", soil, ".", genotype);
			assign(otutable, picrust_ko[ sdata.rarefied$Material == material & sdata.rarefied$Soil == soil & sdata.rarefied$Genotype == genotype, sdata.rarefied$Material == material & sdata.rarefied$Soil == soil & sdata.rarefied$Genotype == genotype ])
			distmat <- paste0(otutable, '.horn');
			assign( distmat, vegdist(get(otutable), dist='horn') );
			sampledata <- paste0("sdata.rarefied.", material, ".", soil, ".", genotype);
			assign(sampledata, sdata.rarefied[ sdata.rarefied$Material == material & sdata.rarefied$Soil == soil & sdata.rarefied$Genotype == genotype, ]);
			var_part <- paste0(otutable, ".varpart");
			assign( var_part, varpart(get(otutable), ~ status, ~ Inoculation, data=get(sampledata)) );
			ord_status <- paste0(otutable, ".status.rda");
			assign(ord_status, dbrda(get(otutable) ~ status + Condition(Inoculation), data=get(sampledata)) );
			ord_inoc <- paste0(otutable, ".inoc.rda");
			assign( ord_inoc, dbrda(get(otutable) ~ Inoculation + Condition(status), data=get(sampledata)) );
				ordanova_inoc <- paste0(ord_inoc, ".anova");
			assign( ordanova_inoc, anova(get(ord_inoc), permu=999) );
			ordanova_status <- paste0(ord_status, ".anova");
			assign( ordanova_status, anova(get(ord_status), permu=999) );
			row <- c( material, soil, genotype, get(var_part)$part$indfract$Adj.R.squared[1], get(var_part)$part$indfract$Adj.R.squared[3], get(ordanova_status)$"Pr(>F)"[1], get(ordanova_inoc)$"Pr(>F)"[1] );
			print( row );
			picrust_ko.inoculation_influence.horn <- rbind( picrust_ko.inoculation_influence.horn, row );
		}
	}
}
colnames(picrust_ko.inoculation_influence.horn) <- c("Material", "Soil", "Genotype", "var_expl_status", "var_expl_inoc", "p_status", "p_inoc")
picrust_ko.inoculation_influence.horn$var_expl_status <- as.numeric(picrust_ko.inoculation_influence.horn$var_expl_status)
picrust_ko.inoculation_influence.horn$var_expl_inoc <- as.numeric(picrust_ko.inoculation_influence.horn$var_expl_inoc)

svg( "picrust_ko.status_influence.svg", width=3.5, height=3.5, pointsize=3 );
p <- ggplot(picrust_ko.inoculation_influence.horn, aes(x=Genotype, y=var_expl_status, fill=Soil)) + geom_bar(stat="identity", position=position_dodge()) + coord_cartesian(ylim=c(0.0, 0.25)) + facet_wrap( ~ Material ) + scale_x_discrete(drop=T) + xlab("Genotype") + ylab("Explained variation fraction")
print(p);
dev.off();

svg( "picrust_ko.inoculation_influence.svg", width=3.5, height=3.5, pointsize=3 );
p <- ggplot(picrust_ko.inoculation_influence.horn, aes(x=Genotype, y=var_expl_inoc, fill=Soil)) + geom_bar(stat="identity", position=position_dodge()) + coord_cartesian(ylim=c(0.0, 0.1)) + facet_wrap( ~ Material ) + scale_x_discrete(drop=T) + xlab("Genotype") + ylab("Explained variation fraction")
print(p);
dev.off();

### PICRUSt2-predicted pathways – using Morisita-Horn distances
# whole data – checking for significance of materials


# data divided according to material – checking for significance of status.aldex


# data divided according to experimental variant – checking for significance of status and inoculation
```

**SM7h.** Variance partitioning

```
library(vegan)
### amplicon data, generalized d05 UniFrac distance used
# whole data
bacteria.rarefied.d05unifrac.varpart <- varpart( as.dist(bacteria.rarefied.d05unifrac), ~ Material, ~ Soil, ~ Genotype, ~ Time, data=sdata.rarefied )

bacteria.inoculation_influence.d05unifrac <- data.frame( matrix(ncol=7, nrow=0) );
for( material in c("S", "R", "L") ){
	otutable <- paste0("bacteria.rarefied.d05unifrac.", material);
	assign(otutable, bacteria.rarefied.d05unifrac[ sdata.rarefied$Material == material , sdata.rarefied$Material == material  ])
	sampledata <- paste0("sdata.rarefied.", material);
	assign(sampledata, sdata.rarefied[ sdata.rarefied$Material == material, ]);
	var_part <- paste0(otutable, ".varpart");
	assign( var_part, varpart(as.dist(get(otutable)), ~ status, ~ Soil, ~ Genotype, ~ Inoculation, data=get(sampledata)) );
	for( soil in c("S1", "S2") ){
		for( genotype in c("B", "C", "M") ){
			otutable <- paste0("bacteria.rarefied.d05unifrac.", material, ".", soil, ".", genotype);
			assign(otutable, bacteria.rarefied.d05unifrac[ sdata.rarefied$Material == material & sdata.rarefied$Soil == soil & sdata.rarefied$Genotype == genotype, sdata.rarefied$Material == material & sdata.rarefied$Soil == soil & sdata.rarefied$Genotype == genotype ])
			sampledata <- paste0("sdata.rarefied.", material, ".", soil, ".", genotype);
			assign(sampledata, sdata.rarefied[ sdata.rarefied$Material == material & sdata.rarefied$Soil == soil & sdata.rarefied$Genotype == genotype, ]);
			var_part <- paste0(otutable, ".varpart");
			assign( var_part, varpart(as.dist(get(otutable)), ~ status, ~ Inoculation, data=get(sampledata)) );
			ord_status <- paste0(otutable, ".status.rda");
			assign(ord_status, dbrda(as.dist(get(otutable)) ~ status + Condition(Inoculation), data=get(sampledata)) );
			ord_inoc <- paste0(otutable, ".inoc.rda");
			assign( ord_inoc, dbrda(as.dist(get(otutable)) ~ Inoculation + Condition(status), data=get(sampledata)) );
				ordanova_inoc <- paste0(ord_inoc, ".anova");
			assign( ordanova_inoc, anova(get(ord_inoc), permu=999) );
			ordanova_status <- paste0(ord_status, ".anova");
			assign( ordanova_status, anova(get(ord_status), permu=999) );
			row <- c( material, soil, genotype, get(var_part)$part$indfract$Adj.R.squared[1], get(var_part)$part$indfract$Adj.R.squared[3], get(ordanova_status)$"Pr(>F)"[1], get(ordanova_inoc)$"Pr(>F)"[1] );
			print( row );
			bacteria.inoculation_influence.d05unifrac <- rbind( bacteria.inoculation_influence.d05unifrac, row );
		}
	}
}
colnames(bacteria.inoculation_influence.d05unifrac) <- c("Material", "Soil", "Genotype", "var_expl_status", "var_expl_inoc", "p_status", "p_inoc")
bacteria.inoculation_influence.d05unifrac$var_expl_status <- as.numeric(bacteria.inoculation_influence.d05unifrac$var_expl_status)
bacteria.inoculation_influence.d05unifrac$var_expl_inoc <- as.numeric(bacteria.inoculation_influence.d05unifrac$var_expl_inoc)

svg( "bacteria.status_influence.svg", width=7, height=7, pointsize=3 );
p <- ggplot(bacteria.inoculation_influence.d05unifrac, aes_(x=bacteria.inoculation_influence.d05unifrac$Genotype, y=bacteria.inoculation_influence.d05unifrac$var_expl_status, fill=bacteria.inoculation_influence.d05unifrac$Soil)) + geom_bar(stat="identity", position=position_dodge()) + coord_cartesian(ylim=c(0.0, 0.3)) + facet_wrap( ~ bacteria.inoculation_influence.d05unifrac$Material ) + scale_x_discrete(drop=T) + xlab("Genotype") + ylab("Explained variation fraction")
print(p);
dev.off();

svg( "bacteria.inoculation_influence.svg", width=7, height=7, pointsize=3 );
p <- ggplot(bacteria.inoculation_influence.d05unifrac, aes_(x=bacteria.inoculation_influence.d05unifrac$Genotype, y=bacteria.inoculation_influence.d05unifrac$var_expl_inoc, fill=bacteria.inoculation_influence.d05unifrac$Soil)) + geom_bar(stat="identity", position=position_dodge()) + coord_cartesian(ylim=c(0.0, 0.03)) + facet_wrap( ~ bacteria.inoculation_influence.d05unifrac$Material ) + scale_x_discrete(drop=T) + xlab("Genotype") + ylab("Explained variation fraction")
print(p);
dev.off();


### PICRUSt2-predicted functions
picrust_ko.inoculation_influence.horn <- data.frame( matrix(ncol=7, nrow=0) );
for( material in c("S", "R", "L") ){
	otutable <- paste0("picrust_ko.horn.", material);
	assign(otutable, picrust_ko[ sdata.rarefied$Material == material , sdata.rarefied$Material == material  ])
	sampledata <- paste0("sdata.rarefied.", material);
	assign(sampledata, sdata.rarefied[ sdata.rarefied$Material == material, ]);
	distmat <- paste0(otutable, '.horn');
	assign( distmat, vegdist(get(otutable), dist='horn') );
	var_part <- paste0(distmat, ".varpart");
	assign( var_part, varpart(get(distmat), ~ status, ~ Soil, ~ Genotype, ~ Inoculation, data=get(sampledata)) );
	for( soil in c("S1", "S2") ){
		for( genotype in c("B", "C", "M") ){
			otutable <- paste0("picrust_ko.", material, ".", soil, ".", genotype);
			assign(otutable, picrust_ko[ sdata.rarefied$Material == material & sdata.rarefied$Soil == soil & sdata.rarefied$Genotype == genotype, sdata.rarefied$Material == material & sdata.rarefied$Soil == soil & sdata.rarefied$Genotype == genotype ])
			distmat <- paste0(otutable, '.horn');
			assign( distmat, vegdist(get(otutable), dist='horn') );
			sampledata <- paste0("sdata.rarefied.", material, ".", soil, ".", genotype);
			assign(sampledata, sdata.rarefied[ sdata.rarefied$Material == material & sdata.rarefied$Soil == soil & sdata.rarefied$Genotype == genotype, ]);
			var_part <- paste0(otutable, ".varpart");
			assign( var_part, varpart(get(otutable), ~ status, ~ Inoculation, data=get(sampledata)) );
			ord_status <- paste0(otutable, ".status.rda");
			assign(ord_status, dbrda(get(otutable) ~ status + Condition(Inoculation), data=get(sampledata)) );
			ord_inoc <- paste0(otutable, ".inoc.rda");
			assign( ord_inoc, dbrda(get(otutable) ~ Inoculation + Condition(status), data=get(sampledata)) );
				ordanova_inoc <- paste0(ord_inoc, ".anova");
			assign( ordanova_inoc, anova(get(ord_inoc), permu=999) );
			ordanova_status <- paste0(ord_status, ".anova");
			assign( ordanova_status, anova(get(ord_status), permu=999) );
			row <- c( material, soil, genotype, get(var_part)$part$indfract$Adj.R.squared[1], get(var_part)$part$indfract$Adj.R.squared[3], get(ordanova_status)$"Pr(>F)"[1], get(ordanova_inoc)$"Pr(>F)"[1] );
			print( row );
			picrust_ko.inoculation_influence.horn <- rbind( picrust_ko.inoculation_influence.horn, row );
		}
	}
}
colnames(picrust_ko.inoculation_influence.horn) <- c("Material", "Soil", "Genotype", "var_expl_status", "var_expl_inoc", "p_status", "p_inoc")
picrust_ko.inoculation_influence.horn$var_expl_status <- as.numeric(picrust_ko.inoculation_influence.horn$var_expl_status)
picrust_ko.inoculation_influence.horn$var_expl_inoc <- as.numeric(picrust_ko.inoculation_influence.horn$var_expl_inoc)

svg( "picrust_ko.status_influence.svg", width=3.5, height=3.5, pointsize=3 );
p <- ggplot(picrust_ko.inoculation_influence.horn, aes(x=Genotype, y=var_expl_status, fill=Soil)) + geom_bar(stat="identity", position=position_dodge()) + coord_cartesian(ylim=c(0.0, 0.25)) + facet_wrap( ~ Material ) + scale_x_discrete(drop=T) + xlab("Genotype") + ylab("Explained variation fraction")
print(p);
dev.off();

svg( "picrust_ko.inoculation_influence.svg", width=3.5, height=3.5, pointsize=3 );
p <- ggplot(picrust_ko.inoculation_influence.horn, aes(x=Genotype, y=var_expl_inoc, fill=Soil)) + geom_bar(stat="identity", position=position_dodge()) + coord_cartesian(ylim=c(0.0, 0.1)) + facet_wrap( ~ Material ) + scale_x_discrete(drop=T) + xlab("Genotype") + ylab("Explained variation fraction")
print(p);
dev.off();


### PICRUSt2-predicted pathways
```

**SM7i.** Preparation of taxonomy barplots

```
bacteria <- readRDS("betamikro_WP5_complete_bacteria.rds")
bacteria.tax <- readRDS("betamikro_WP5_complete_bacteria.tax.rds")
sdata <- read.table("WP5.design.csv", header=T, sep=",")
sdata$status <- 'early'
sdata$status[sdata$Time == 'T3' | sdata$Time == 'T4'] <- 'late'
rownames(sdata) <- sdata$sample
sdata$Material <- substr(rownames(sdata), 1,1)
sdata$Material[sdata$Material == "G"] <- "S"
sdata$Material[sdata$Material == "K"] <- "R"
sdata <- sdata[ rownames(sdata) %in% rownames(bacteria), ]
bacteria.t <- t(bacteria)
for( taxlevelpl in c("phyla", "classes", "orders", "families", "genera") ){
	if( taxlevelpl == "phyla" ){
		taxlevelsngl <- "Phylum";
		} else if( taxlevelpl == "classes" ){
		taxlevelsngl <- "Class";
		} else if( taxlevelpl == "orders" ){
		taxlevelsngl <- "Order";
		} else if( taxlevelpl == "families" ){
		taxlevelsngl <- "Family";
		} else {
		taxlevelsngl <- "Genus";
		}

	taxtmp <-  as.data.frame(bacteria.t);
	taxtmp$taxlvl <- bacteria.tax[[taxlevelsngl]];
	taxtmp <-  aggregate(. ~ taxlvl, data=taxtmp, FUN='sum');
	rownames(taxtmp) <- taxtmp$taxlvl;
	taxtmp$taxlvl <- NULL;
	taxtmp <- t(taxtmp)
	taxtmp <- taxtmp/rowSums(taxtmp)
	taxtmp <- as.data.frame(t(taxtmp));
	assign(taxlevelpl, as.data.frame(t(taxtmp)));
	taxlevelpl_unclassified <- paste0(taxlevelpl, "_unclassified");
	taxlevelpl_rare <- paste0(taxlevelpl, "_rare");
	taxlevelpl_inoc <- paste0(taxlevelpl, "_inoc");
	taxlevelpl_final <- paste0(taxlevelpl, "_final");
	assign( taxlevelpl_unclassified, get(taxlevelpl)[ , grep("unknown|unclassified|_ge|_fa|_or|_cl|_ph", colnames(get(taxlevelpl))) ] );
	assign( taxlevelpl_final, get(taxlevelpl)[ , !colnames(get(taxlevelpl)) %in% colnames(get(taxlevelpl_unclassified)) ] );
	assign( taxlevelpl_rare, get(taxlevelpl_final)[ , colSums(get(taxlevelpl_final))/sum(get(taxlevelpl_final)) < 0.02] );
	assign( taxlevelpl_final, get(taxlevelpl_final)[ , !colnames(get(taxlevelpl_final)) %in% colnames(get(taxlevelpl_rare)) ] );
	sdata_noinoc <- sdata[ -grep("I", rownames(sdata)), ];
	tempobj <- get( taxlevelpl_final );
	tempobj$rare <- rowSums(get(taxlevelpl_rare));
	tempobj$unclassified <- rowSums(get(taxlevelpl_unclassified));
	assign( taxlevelpl_final, tempobj );
	assign( taxlevelpl_inoc, get(taxlevelpl_final)[ grep("I", rownames(get(taxlevelpl_final))), ] );
	assign( taxlevelpl_final, get(taxlevelpl_final)[ -grep("I", rownames(get(taxlevelpl_final))), ] );
	tempobj <- get( taxlevelpl_final );
	tempobj$Time <- sdata_noinoc$Time;
	tempobj$Material <- sdata_noinoc$Material;
	tempobj_aggregated_time <- aggregate(. ~ Time *Material, data=tempobj, FUN="mean")
	tempobj$Soil <- sdata_noinoc$Soil;
	tempobj$Genotype <- sdata_noinoc$Genotype;
	tempobj$Inoculation <- sdata_noinoc$Inoculation;
	tempobj$status <- sdata_noinoc$status;
	tempobj$sample <- sdata_noinoc$sample;
	assign( taxlevelpl_final, tempobj );
	taxlevelpl_final_molten <- paste0(taxlevelpl_final, "_molten");
	assign( taxlevelpl_final_molten, melt(get(taxlevelpl_final), id.vars=c('sample', 'Material', 'status', 'Time', 'Soil', 'Genotype', 'Inoculation'), value.name='abundance', variable.name=taxlevelsngl) );
	file_material <- paste0(taxlevelpl, "_material.svg");
	file_material_status <- paste0(taxlevelpl, "_material_status.svg");
	file_material_time <- paste0(taxlevelpl, "_material_time.svg");
	file_all_vars <- paste0(taxlevelpl, "_all_vars.svg");


	tempobj_aggregated_time_molten <- melt(tempobj_aggregated_time, id.vars=c('Material', 'Time'), value.name='abundance', variable.name=taxlevelsngl)
	svg(file_material_time, width=3.5, height=2, pointsize=6);
	p <- ggplot( tempobj_aggregated_time_molten, aes_(x=tempobj_aggregated_time_molten$Time, y=tempobj_aggregated_time_molten$abundance, fill=tempobj_aggregated_time_molten[[taxlevelsngl]])) +
	geom_bar(position='fill', stat='identity', width=1) +
	facet_wrap( ~ Material, scales='free_x') +
	xlab("Timepoint") + ylab("Relative abundance") + labs(fill=taxlevelsngl) +
	theme_gray(base_size=8) +
	theme(axis.text=element_text(size=6), legend.text=element_text(size=6), axis.ticks.x=element_blank());
	print( p );
	dev.off();

	svg(file_all_vars, width=7, height=10.5, pointsize=6);
	p <- ggplot( get(taxlevelpl_final_molten), aes_(x=get(taxlevelpl_final_molten)$sample, y=get(taxlevelpl_final_molten)$abundance, fill=get(taxlevelpl_final_molten)[[taxlevelsngl]])) + geom_bar(position='fill', stat='identity', width=1) + facet_grid(Soil + Genotype ~ Material + status, scales='free_x') + xlab("") + ylab("Relative abundance") + labs(fill=taxlevelsngl) + theme(axis.title.x=element_blank(), axis.text.x=element_blank(), axis.ticks.x=element_blank());
	print( p );
	dev.off();
}

# Fig. 2E and 5G
sdata_noinoc <- sdata[ -grep("I", rownames(sdata)), ]
genera.t <- genera
genera_unclassified <- as.data.frame(genera.t[, grep("unknown|unclassified|_ge|_fa|_or|_cl|_ph", colnames(genera.t))])
genera_final <- genera.t[ , !colnames(genera.t) %in% colnames(genera_unclassified) ]
genera_final_inoculant <- genera_final[ grep("I", rownames(genera_final)), ]
genera_rare <- genera_final[ , colSums(genera_final)/sum(genera_final) < 0.02 ]
genera_inoculant_rare <- genera_final_inoculant[ , colSums(genera_final_inoculant)/sum(genera_final_inoculant) < 0.02 ]
genera_final <- as.data.frame(genera_final[ , !colnames(genera_final) %in% colnames(genera_rare) ])
genera_final_inoculant <- as.data.frame(genera_final_inoculant[ , !colnames(genera_final_inoculant) %in% colnames(genera_inoculant_rare) ])
genera_final$rare <- rowSums(genera_rare)
genera_final_inoculant$rare <- rowSums(genera_inoculant_rare)
genera_final$unclassfied <- rowSums(genera_unclassified)
genera_unclassified_inoculant <- genera_unclassified[ grep("I", rownames(genera_unclassified)), ]
genera_final_inoculant$unclassified <- rowSums(genera_unclassified_inoculant)
genera_final_noinoc <- genera_final[ -grep("I", rownames(genera_final)), ]
genera_final_noinoc$sample <- rownames(genera_final_noinoc)
genera_final_inoculant$sample <- rownames(genera_final_inoculant)
genera_final_noinoc$Material <- sdata_noinoc$Material
genera_final.molten <- melt(genera_final_noinoc, id.vars=c('sample', 'Material'), value.name='abundance', variable.name='genus')
genera_final_inoculant.molten <- melt(genera_final_inoculant, id.vars=c('sample'), value.name='abundance', variable.name='genus')


svg("genera_material.svg", width=7, height=3.5, pointsize=6) # Fig. 2E
p <- ggplot( genera_final.molten, aes(x=sample, y=abundance, fill=genus) ) + geom_bar(position='fill', stat='identity', width=1) + facet_wrap( ~ Material, ncol=3, scales='free_x') + xlab("") + ylab("Relative abundance") + theme(axis.title.x=element_blank(), axis.text.x=element_blank(), axis.ticks.x=element_blank())
print(p)
dev.off()
quit()
```

**SM7j.** Finding signature ASVs with biosigner

```
bacteria.rarefied.nzv <- nearZeroVar(bacteria.rarefied)
bacteria.rarefied.lv <- bacteria.rarefied[ , -bacteria.rarefied.nzv$Position ]
inocSign <- biosigner::biosign(bacteria.rarefied.lv, sdata.rarefied$Inoculation)

bacteria.rarefied.complete_signature <- bacteria.tax[ rownames(bacteria.tax) %in% getSignatureLs(inocSign)$complete, ]
bacteria.rarefied.inocSign <- bacteria.rarefied[ ,colnames(bacteria.rarefied) %in% getSignatureLs(inocSign)$complete ]
bacteria.rarefied.inocSign.aggregated <- aggregate(bacteria.rarefied.inocSign ~ sdata.rarefied$Inoculation, FUN='mean')
rownames(bacteria.rarefied.inocSign.aggregated) <- bacteria.rarefied.inocSign.aggregated$"sdata.rarefied$Inoculation"
bacteria.rarefied.inocSign.aggregated$"sdata.rarefied$Inoculation" <- NULL
bacteria.rarefied.inocSign.aggregated.t <- t(bacteria.rarefied.inocSign.aggregated)
bacteria.rarefied.complete_signature <- cbind( bacteria.rarefied.complete_signature, bacteria.rarefied.inocSign.aggregated.t)
write.table(bacteria.rarefied.complete_signature, "bacteria.rarefied.complete_signature.csv", sep="\t", dec=".")


bacteria.rarefied.complete_signature <- bacteria.tax[ rownames(bacteria.tax) %in% getSignatureLs(inocSign)$complete, ]
bacteria.rarefied.inocSign <- bacteria.rarefied[ ,colnames(bacteria.rarefied) %in% getSignatureLs(inocSign)$complete ]
bacteria.rarefied.inocSign.aggregated <- aggregate(bacteria.rarefied.inocSign ~ sdata.rarefied$Inoculation, FUN='mean')
rownames(bacteria.rarefied.inocSign.aggregated) <- bacteria.rarefied.inocSign.aggregated$"sdata.rarefied$Inoculation"
bacteria.rarefied.inocSign.aggregated$"sdata.rarefied$Inoculation" <- NULL
bacteria.rarefied.inocSign.aggregated.t <- t(bacteria.rarefied.inocSign.aggregated)
bacteria.rarefied.complete_signature <- cbind( bacteria.rarefied.complete_signature, bacteria.rarefied.inocSign.aggregated.t)
write.table(bacteria.rarefied.complete_signature, "bacteria.rarefied.complete_signature.csv", sep="\t", dec=".", col.names=NA )


# late samples only

for( material in c("S", "R", "L") ){
	otutable <- paste0("bacteria.rarefied.", material);
	otutablenzv <- paste0(otutable, ".nzv");
	otutablelv <- paste0(otutable, ".lv");
	sampledata <- paste0("sdata.rarefied.", material );
	signat <- paste0("inocSign.", material);
	comp_signat <- paste0(otutable, ".complete_signature");
	file <- paste0(comp_signat, ".csv");
	print( file );
	assign(otutable, bacteria.rarefied[ sdata.rarefied$Material == material, ]);
	assign(sampledata, sdata.rarefied[ sdata.rarefied$Material == material, ]);
	assign(otutablenzv, nearZeroVar(get(otutable)));
	print( otutablenzv );
	assign(otutablelv, get(otutable)[ , -get(nzv)$Position ] );
	assign(signat, biosigner::biosign(get(otutablelv), get(sampledata)$Inoculation));
	if( length(getSignatureLs(get(signat))$complete) ){
		assign( comp_signat, bacteria.tax[ rownames(bacteria.tax) %in% getSignatureLs(get(signat))$complete, ] );
		temp <- as.data.frame( get(otutable)[ , colnames(get(otutable)) %in% getSignatureLs(get(signat))$complete ] );
		rownames(temp) <- rownames(get(otutable));
		temp.aggregated <- aggregate( . ~ get(sampledata)$Inoculation, data=temp, FUN='mean' );
		rownames(temp.aggregated) <- temp.aggregated$"get(sampledata)$Inoculation";
		temp.aggregated$"get(sampledata)$Inoculation" <- NULL;
		temp.aggregated.t <- t(temp.aggregated);
		assign( comp_signat, cbind( get(comp_signat), temp.aggregated.t ) );
		write.table( get(comp_signat), file, sep="\t", dec=".", col.names=NA );
	} else {
		print( "No features selected" );
	}

	for( status in c('early', 'late') ){
		otutable <- paste0("bacteria.rarefied.", material, ".", status);
		otutablenzv <- paste0(otutable, ".nzv");
		otutablelv <- paste0(otutable, ".lv");
		sampledata <- paste0("sdata.rarefied.", material, ".", status);
		signat <- paste0("inocSign.", material, ".", status );
		comp_signat <- paste0(otutable, ".complete_signature");
		file <- paste0(comp_signat, ".csv");
		print( file );
		assign(otutable, bacteria.rarefied[ sdata.rarefied$Material == material & sdata.rarefied$status == status, ]);
		assign(sampledata, sdata.rarefied[ sdata.rarefied$Material == material & sdata.rarefied$status == status, ]);
		assign(otutablenzv, nearZeroVar(get(otutable)));
		print( otutablenzv );
		assign(otutablelv, get(otutable)[ , -get(nzv)$Position ] );
		assign(signat, biosigner::biosign(get(otutablelv), get(sampledata)$Inoculation));
		if( length(getSignatureLs(get(signat))$complete) ){
			assign( comp_signat, bacteria.tax[ rownames(bacteria.tax) %in% getSignatureLs(get(signat))$complete, ] );
			temp <- as.data.frame( get(otutable)[ , colnames(get(otutable)) %in% getSignatureLs(get(signat))$complete ] );
			rownames(temp) <- rownames(get(otutable));
			temp.aggregated <- aggregate( . ~ get(sampledata)$Inoculation, data=temp, FUN='mean' );
			rownames(temp.aggregated) <- temp.aggregated$"get(sampledata)$Inoculation";
			temp.aggregated$"get(sampledata)$Inoculation" <- NULL;
			temp.aggregated.t <- t(temp.aggregated);
			assign( comp_signat, cbind( get(comp_signat), temp.aggregated.t ) );
			write.table( get(comp_signat), file, sep="\t", dec=".", col.names=NA );
		} else {
			print( "No features selected" );
		}
	}
	for( soil  in c("S1", "S2") ){
		otutable <- paste0("bacteria.rarefied.", material, ".", soil);
		otutablenzv <- paste0(otutable, ".nzv");
		otutablelv <- paste0(otutable, ".lv");
		sampledata <- paste0("sdata.rarefied.", material, ".", soil );
		signat <- paste0("inocSign.", material, ".", soil);
		comp_signat <- paste0(otutable, ".complete_signature");
		file <- paste0(comp_signat, ".csv");
		print( file );
		assign(otutable, bacteria.rarefied[ sdata.rarefied$Material == material & sdata.rarefied$Soil == soil, ]);
		assign(sampledata, sdata.rarefied[ sdata.rarefied$Material == material & sdata.rarefied$Soil == soil, ]);
		assign(otutablenzv, nearZeroVar(get(otutable)));
		print( otutablenzv );
		assign(otutablelv, get(otutable)[ , -get(nzv)$Position ] );
		assign(signat, biosigner::biosign(get(otutablelv), get(sampledata)$Inoculation));
		if( length(getSignatureLs(get(signat))$complete) ){
			assign( comp_signat, bacteria.tax[ rownames(bacteria.tax) %in% getSignatureLs(get(signat))$complete, ] );
			temp <- as.data.frame( get(otutable)[ , colnames(get(otutable)) %in% getSignatureLs(get(signat))$complete ] );
			rownames(temp) <- rownames(get(otutable));
			temp.aggregated <- aggregate( . ~ get(sampledata)$Inoculation, data=temp, FUN='mean' );
			rownames(temp.aggregated) <- temp.aggregated$"get(sampledata)$Inoculation";
			temp.aggregated$"get(sampledata)$Inoculation" <- NULL;
			temp.aggregated.t <- t(temp.aggregated);
			assign( comp_signat, cbind( get(comp_signat), temp.aggregated.t ) );
			write.table( get(comp_signat), file, sep="\t", dec=".", col.names=NA );
		} else {
			print( "No features selected" );
		}
		for( status in c('early', 'late') ){
			otutable <- paste0("bacteria.rarefied.", material, ".", soil, ".", status);
			otutablenzv <- paste0(otutable, ".nzv");
			otutablelv <- paste0(otutable, ".lv");
			sampledata <- paste0("sdata.rarefied.", material, ".", soil, ".", status);
			signat <- paste0("inocSign.", material, ".", soil, ".", status );
			comp_signat <- paste0(otutable, ".complete_signature");
			file <- paste0(comp_signat, ".csv");
			print( file );
			assign(otutable, bacteria.rarefied[ sdata.rarefied$Material == material & sdata.rarefied$Soil == soil & sdata.rarefied$status == status, ]);
			assign(sampledata, sdata.rarefied[ sdata.rarefied$Material == material & sdata.rarefied$Soil == soil & sdata.rarefied$status == status, ]);
			assign(otutablenzv, nearZeroVar(get(otutable)));
			print( otutablenzv );
			assign(otutablelv, get(otutable)[ , -get(nzv)$Position ] );
			assign(signat, biosigner::biosign(get(otutablelv), get(sampledata)$Inoculation));
			if( length(getSignatureLs(get(signat))$complete) ){
				assign( comp_signat, bacteria.tax[ rownames(bacteria.tax) %in% getSignatureLs(get(signat))$complete, ] );
				temp <- as.data.frame( get(otutable)[ , colnames(get(otutable)) %in% getSignatureLs(get(signat))$complete ] );
				rownames(temp) <- rownames(get(otutable));
				temp.aggregated <- aggregate( . ~ get(sampledata)$Inoculation, data=temp, FUN='mean' );
				rownames(temp.aggregated) <- temp.aggregated$"get(sampledata)$Inoculation";
				temp.aggregated$"get(sampledata)$Inoculation" <- NULL;
				temp.aggregated.t <- t(temp.aggregated);
				assign( comp_signat, cbind( get(comp_signat), temp.aggregated.t ) );
				write.table( get(comp_signat), file, sep="\t", dec=".", col.names=NA );
			} else {
				print( "No features selected" );
			}
		}

		for( genotype in c("B", "C", "M") ){
			otutable <- paste0("bacteria.rarefied.", material, ".", soil, ".", genotype);
			otutablenzv <- paste0(otutable, ".nzv");
			otutablelv <- paste0(otutable, ".lv");
			sampledata <- paste0("sdata.rarefied.", material, ".", soil, ".", genotype );
			signat <- paste0("inocSign.", material, ".", soil, ".", genotype);
			comp_signat <- paste0(otutable, ".complete_signature");
			file <- paste0(comp_signat, ".csv");
			print( file );
			assign(otutable, bacteria.rarefied[ sdata.rarefied$Material == material & sdata.rarefied$Soil == soil & sdata.rarefied$Genotype == genotype, ]);
			assign(sampledata, sdata.rarefied[ sdata.rarefied$Material == material & sdata.rarefied$Soil == soil & sdata.rarefied$Genotype == genotype, ]);
			assign(otutablenzv, nearZeroVar(get(otutable)));
			print( otutablenzv );
			assign(otutablelv, get(otutable)[ , -get(nzv)$Position ] );
			assign(signat, biosigner::biosign(get(otutablelv), get(sampledata)$Inoculation));
			if( length(getSignatureLs(get(signat))$complete) ){
				assign( comp_signat, bacteria.tax[ rownames(bacteria.tax) %in% getSignatureLs(get(signat))$complete, ] );
				temp <- as.data.frame( get(otutable)[ , colnames(get(otutable)) %in% getSignatureLs(get(signat))$complete ] );
				rownames(temp) <- rownames(get(otutable));
				temp.aggregated <- aggregate( . ~ get(sampledata)$Inoculation, data=temp, FUN='mean' );
				rownames(temp.aggregated) <- temp.aggregated$"get(sampledata)$Inoculation";
				temp.aggregated$"get(sampledata)$Inoculation" <- NULL;
				temp.aggregated.t <- t(temp.aggregated);
				assign( comp_signat, cbind( get(comp_signat), temp.aggregated.t ) );
				write.table( get(comp_signat), file, sep="\t", dec=".", col.names=NA );
			} else {
				print( "No features selected" );
			}
			for( status in c('early', 'late') ){
				otutable <- paste0("bacteria.rarefied.", material, ".", soil, ".", genotype, ".", status);
				otutablenzv <- paste0(otutable, ".nzv");
				otutablelv <- paste0(otutable, ".lv");
				sampledata <- paste0("sdata.rarefied.", material, ".", soil, ".", genotype, ".", status);
				signat <- paste0("inocSign.", material, ".", soil, ".", genotype, ".", status );
				comp_signat <- paste0(otutable, ".complete_signature");
				file <- paste0(comp_signat, ".csv");
				print( file );
				assign(otutable, bacteria.rarefied[ sdata.rarefied$Material == material & sdata.rarefied$Soil == soil & sdata.rarefied$Genotype == genotype & sdata.rarefied$status == status, ]);
				assign(sampledata, sdata.rarefied[ sdata.rarefied$Material == material & sdata.rarefied$Soil == soil & sdata.rarefied$Genotype == genotype& sdata.rarefied$status == status, ]);
				assign(otutablenzv, nearZeroVar(get(otutable)));
				print( otutablenzv );
				assign(otutablelv, get(otutable)[ , -get(nzv)$Position ] );
				assign(signat, biosigner::biosign(get(otutablelv), get(sampledata)$Inoculation));
				if( length(getSignatureLs(get(signat))$complete) ){
					assign( comp_signat, bacteria.tax[ rownames(bacteria.tax) %in% getSignatureLs(get(signat))$complete, ] );
					temp <- as.data.frame( get(otutable)[ , colnames(get(otutable)) %in% getSignatureLs(get(signat))$complete ] );
					rownames(temp) <- rownames(get(otutable));
					temp.aggregated <- aggregate( . ~ get(sampledata)$Inoculation, data=temp, FUN='mean' );
					rownames(temp.aggregated) <- temp.aggregated$"get(sampledata)$Inoculation";
					temp.aggregated$"get(sampledata)$Inoculation" <- NULL;
					temp.aggregated.t <- t(temp.aggregated);
					assign( comp_signat, cbind( get(comp_signat), temp.aggregated.t ) );
					write.table( get(comp_signat), file, sep="\t", dec=".", col.names=NA );
				} else {
					print( "No features selected" );
				}
			}
		}
	}
}
```

**SM7k.** Identification of differentially abundant ASVs and taxa with ALDEx2

ALDEx2 analysis was carried out on a large-variance subset of rarefied data. It was necessary to complete the computations in reasonable time. Low variance features were identified with the nearZeroVar function of the mixOmics package and removed.

```
library(mixOmics)
library(ALDEx2)
library(BiocParallel)
bpparam <- MulticoreParam(30)

### amplicon data
# whole data – identifying ASVs characteristic for leaves, roots or soils
bacteria.nzv <- nearZeroVar( bacteria )
bacteria.lv <- bacteria[ , -bacteria.nzv$Position ]
bacteria.lv.material.clr <- aldex.clr( as.data.frame(t(bacteria.lv)), sdata$Material, mc.samples=128, useMC=T )
bacteria.lv.material.aldex <- aldex.kw(bacteria.lv.material.clr, useMC=T )
bacteria.lv.material.aldex.significant <- bacteria.lv.material.aldex[ bacteria.lv.material.aldex$kw.eBH < 0.05 | bacteria.lv.material.aldex$glm.eBH < 0.05, ]
bacteria.lv.material.aldex.significant <- merge( bacteria.lv.material.aldex.significant, bacteria.tax, by.x=0, by.y=0, all.x=T )
bacteria.lv.material.aldex.significant <- bacteria.lv.material.aldex.significant[ order(bacteria.lv.material.aldex.significant$kw.eBH, decreasing=T), ]
write.table( bacteria.lv.material.aldex.significant, "bacteria.lv.material.aldex.significant.csv", sep="\t", col.names=NA )

# data divided according to material – identifying ASVs characteristic for late and early samples
for( material in c('L', 'R', 'S') ){
	asvtable <- paste0("bacteria.", material);
	sampledata <- paste0("sdata.", material);
	asvtablenzv <- paste(asvtable, ".nzv");
	asvtablelv <- paste0(asvtable, '.lv');
	clr <- paste0(asvtablelv, ".clr");
	ald <- paste0(asvtablelv, ".aldex");
	aldsign <- paste0(ald, ".significant");
	file <- paste0(aldsign, ".csv");
	assign( asvtable, bacteria[ sdata$Material == material, ] );
	assign( sampledata, sdata[ sdata$Material == material, ] );
	assign( asvtablenzv, nearZeroVar(get(asvtable)) );
	assign( asvtablelv, get(asvtable)[ , -get(asvtablenzv)$Position ] );
	assign( clr, aldex.clr(as.data.frame(t(get(asvtablelv))), get(sampledata)$status, mc.samples=128, useMC=T) );
	assign( ald, aldex.kw(get(clr), useMC=T) );
	assign( aldsign, get(ald)[ get(ald)$kw.eBH < 0.05 | get(ald)$glm.eBH < 0.05, ] );
	assign( aldsign, merge(get(aldsign), bacteria.tax, by.x=0, by.y=0, all.x=T) );
	assign( aldsign, get(aldsign)[ order(get(aldsign)$kw.eBH, decreasing=T), ] );
	write.table( get(aldsign), file, sep="\t", col.names=NA );
}

# data divided according to experimental variant: material * soil * genotype – identifying ASVs characteristic for late and early as well as inoculated and non-inoculated samples

for( material in c( 'S', 'R', 'L' ) ){
	for( soil in c( 'S1', 'S2' ) ){
		for( genotype in c( 'B', 'C', 'M' ) ){
			asvtable <- as.data.frame(round(bacteria.rarefied[ sdata.rarefied$Material == material & sdata.rarefied$Soil == soil & sdata.rarefied$Genotype == genotype, ]));
			sampledata <- sdata.rarefied[ sdata.rarefied$Material == material & sdata.rarefied$Soil == soil & sdata.rarefied$Genotype == genotype, ];
			nzv <- nearZeroVar( asvtable );
			lv <- asvtable[ , -nzv$Position ];
			ald <- paste0("bacteria.rarefied.", material, ".", soil, ".", genotype, ".", "status.aldex");
			file <- paste0( ald, ".csv" );
			aldsign <- paste0("bacteria.rarefied.", material, ".", soil, ".", genotype, ".", "status.aldex.significant");
			if(!file.exists(file) & ncol(lv) > 0 & length(levels(as.factor(sampledata$status))) == 2){
				print( ald );
				clr <- aldex.clr( t(lv), sampledata$status, mc.samples=128, useMC=T );
				assign( ald, aldex.kw( clr, useMC=T ) );
				assign( aldsign, get(ald)[ get(ald)$kw.eBH < 0.05 | get(ald)$glm.eBH < 0.05, ] );
				assign( aldsign, merge(get(aldsign), bacteria.tax, by=0, all.x=T) );
				assign( aldsign, get(aldsign)[ order(get(aldsign)$kw.eBH, decreasing=T), ] );
				write.table(get(aldsign), file, col.names=NA, sep="\t");
				print( "done" );
			}

			for( status in c( 'early', 'late' ) ){
				asvtable <- as.data.frame(round(bacteria.rarefied[ sdata.rarefied$Material == material & sdata.rarefied$Soil == soil & sdata.rarefied$Genotype == genotype & sdata.rarefied$status == status, ]));
				sampledata <- sdata.rarefied[ sdata.rarefied$Material == material & sdata.rarefied$Soil == soil & sdata.rarefied$Genotype == genotype & sdata.rarefied$status == status, ];
				nzv <- nearZeroVar( asvtable );
				lv <- asvtable[ , -nzv$Position ];
				ald <- paste0("bacteria.rarefied.", material, ".", soil, ".", genotype, ".", status, ".Inoculation.aldex");
				file <- paste0( ald, ".csv" );
				aldsign <- paste0("bacteria.rarefied.", material, ".", soil, ".", genotype, ".", status, ".Inoculation.aldex.significant");
				if(!file.exists(file) & ncol(lv) > 0 & length(levels(as.factor(sampledata$Inoculation))) == 2){
					print( ald );
					clr <- aldex.clr( t(lv), sampledata$Inoculation, mc.samples=128, useMC=T );
					assign( ald, aldex.kw( clr, useMC=T ) );
					assign( aldsign, get(ald)[ get(ald)$kw.eBH < 0.05 | get(ald)$glm.eBH < 0.05, ] );
					assign( aldsign, merge(get(aldsign), bacteria.tax, by=0, all.x=T) );
					assign( aldsign, get(aldsign)[ order(get(aldsign)$kw.eBH, decreasing=T), ] );
					write.table(get(aldsign), file, col.names=NA, sep="\t");
					print( "done" );
				}
			}
			for( inoculation in c( 'N', 'I' ) ){
				asvtable <- as.data.frame(round(bacteria.rarefied[ sdata.rarefied$Material == material & sdata.rarefied$Soil == soil & sdata.rarefied$Genotype == genotype & sdata.rarefied$Inoculation == inoculation, ]));
				sampledata <- sdata.rarefied[ sdata.rarefied$Material == material & sdata.rarefied$Soil == soil & sdata.rarefied$Genotype == genotype & sdata.rarefied$Inoculation == inoculation, ];
				nzv <- nearZeroVar( asvtable );
				lv <- asvtable[ , -nzv$Position ];
				ald <- paste0("bacteria.rarefied.", material, ".", soil, ".", genotype, ".", inoculation, ".LateEarly.aldex");
				file <- paste0( ald, ".csv" );
				aldsign <- paste0("bacteria.rarefied.", material, ".", soil, ".", genotype, ".", inoculation, ".LateEarly.aldex.significant");
				if(!file.exists(file) & ncol(lv) > 0 & length(levels(as.factor(sampledata$status))) == 2){
					print( ald );
					clr <- aldex.clr( reads=t(lv), sampledata$status, mc.samples=128, useMC=T );
					assign( ald, aldex.kw( clr, useMC=T ) );
					assign( aldsign, get(ald)[ get(ald)$kw.eBH < 0.05 | get(ald)$glm.eBH < 0.05, ]);
					assign( aldsign, merge(get(aldsign), bacteria.tax, by=0, all.x=T) );
					assign( aldsign, get(aldsign)[ order(get(aldsign)$kw.eBH, decreasing=T), ] );
					write.table(get(aldsign), file, col.names=NA, sep="\t");
					print( "done" );
				}
			}
		}
	}
}


### PICRUSt2-predicted functions
# whole data – identifying functions characteristic for leaves, roots or soils
picrust_ko.nzv <- nearZeroVar( picrust_ko )
picrust_ko.lv <- picrust_ko[ , -picrust_ko.nzv$Position ]
picrust_ko.lv.material.clr <- aldex.clr( as.data.frame(t(picrust_ko.lv)), sdata$Material, mc.samples=128, useMC=T )
picrust_ko.lv.material.aldex <- aldex.kw(picrust_ko.lv.material.clr, useMC=T )
picrust_ko.lv.material.aldex.significant <- picrust_ko.lv.material.aldex[ picrust_ko.lv.material.aldex$kw.eBH < 0.05 | picrust_ko.lv.material.aldex$glm.eBH < 0.05, ]
picrust_ko.lv.material.aldex.significant <- merge( picrust_ko.lv.material.aldex.significant, ko_descriptions, by.x=0, by.y=0, all.x=T )
picrust_ko.lv.material.aldex.significant <- picrust_ko.lv.material.aldex.significant[ order(picrust_ko.lv.material.aldex.significant$kw.eBH, decreasing=T), ]
write.table( picrust_ko.lv.material.aldex.significant, "picrust_ko.lv.material.aldex.significant.csv", sep="\t", col.names=NA )

# data divided according to material – identifying functions characteristic for late and early samples
for( material in c('L','R', 'S') ){
	asvtable <- paste0("picrust_ko.", material);
	sampledata <- paste0("sdata.", material);
	asvtablenzv <- paste(asvtable, ".nzv");
	asvtablelv <- paste0(asvtable, '.lv');
	clr <- paste0(asvtablelv, ".clr");
	ald <- paste0(asvtablelv, ".aldex");
	aldsign <- paste0(ald, ".significant");
	file <- paste0(aldsign, ".csv");
	assign( asvtable, bacteria[ sdata$Material == material, ] );
	assign( sampledata, sdata[ sdata$Material == material, ] );
	assign( asvtablenzv, nearZeroVar(get(asvtable)) );
	assign( asvtablelv, get(asvtable)[ , -get(asvtablenzv)$Position ] );
	assign( clr, aldex.clr(as.data.frame(t(get(asvtablelv))), get(sampledata)$status, mc.samples=128, useMC=T) );
	assign( ald, aldex.kw(get(clr), useMC=T) );
	assign( aldsign, get(ald)[ get(ald)$kw.eBH < 0.05 | get(ald)$glm.eBH < 0.05, ] );
	assign( aldsign, merge(get(aldsign), ko_descriptions, by.x=0, by.y=0, all.x=T) );
	assign( aldsign, get(aldsign)[ order(get(aldsign)$kw.eBH, decreasing=T), ] );
	write.table( get(aldsign), file, sep="\t", col.names=NA );
}

# data divided according to experimental variant: material * soil * genotype – identifying functions characteristic for late and early as well as inoculated and non-inoculated samples

for( material in c( 'S', 'R', 'L' ) ){
	for( soil in c( 'S1', 'S2' ) ){
		for( genotype in c( 'B', 'C', 'M' ) ){
			for( status in c( 'early', 'late' ) ){
				asvtable <- as.data.frame(round(picrust_ko[ sdata.rarefied$Material == material & sdata.rarefied$Soil == soil & sdata.rarefied$Genotype == genotype & sdata.rarefied$status == status, ]));
				sampledata <- sdata.rarefied[ sdata.rarefied$Material == material & sdata.rarefied$Soil == soil & sdata.rarefied$Genotype == genotype & sdata.rarefied$status == status, ];
				nzv <- nearZeroVar( asvtable );
				lv <- asvtable[ , nzv$Position ];
				ald <- paste0("picrust_ko.", material, ".", soil, ".", genotype, ".", status, ".Inoculation.aldex");
				file <- paste0( ald, ".csv" );
				aldsign <- paste0("picrust_ko.", material, ".", soil, ".", genotype, ".", status, ".Inoculation.aldex.significant");
				if(is.null(get0(aldsign)) | !file.exists(file)){
					print( ald );
					clr <- aldex.clr( t(lv), sampledata$Inoculation, mc.samples=128, useMC=T );
					assign( ald, aldex.kw( clr, useMC=T ) );
					assign( aldsign, get(ald)[ get(ald)$kw.eBH < 0.05 | get(ald)$glm.eBH < 0.05, ] );
					assign( aldsign, merge(get(aldsign), ko_descriptions, by=0, all.x=T) );
					assign( aldsign, get(aldsign)[ order(get(aldsign)$kw.eBH, decreasing=T), ] );
					write.table(get(aldsign), file, col.names=NA, sep="\t");
					print( "done" );
				}
			}
			for( inoculation in c( 'N', 'I' ) ){
				asvtable <- as.data.frame(round(picrust_ko[ sdata.rarefied$Material == material & sdata.rarefied$Soil == soil & sdata.rarefied$Genotype == genotype & sdata.rarefied$Inoculation == inoculation, ]));
				sampledata <- sdata.rarefied[ sdata.rarefied$Material == material & sdata.rarefied$Soil == soil & sdata.rarefied$Genotype == genotype & sdata.rarefied$Inoculation == inoculation, ];
				nzv <- nearZeroVar( asvtable );
				lv <- asvtable[ , nzv$Position ];
				ald <- paste0("picrust_ko.", material, ".", soil, ".", genotype, ".", inoculation, ".LateEarly.aldex");
				file <- paste0( ald, ".csv" );
				aldsign <- paste0("picrust_ko.", material, ".", soil, ".", genotype, ".", inoculation, ".LateEarly.aldex.significant");
				if(is.null(get0(aldsign)) | !file.exists(file)){
					print( ald );
					clr <- aldex.clr( reads=t(lv), sampledata$status, mc.samples=128, useMC=T );
					assign( ald, aldex.kw( clr, useMC=T ) );
					assign( aldsign, get(ald)[ get(ald)$kw.eBH < 0.05 | get(ald)$glm.eBH < 0.05, ]);
					assign( aldsign, merge(get(aldsign), ko_descriptions, by=0, all.x=T) );
					assign( aldsign, get(aldsign)[ order(get(aldsign)$kw.eBH, decreasing=T), ] );
					write.table(get(aldsign), file, col.names=NA, sep="\t");
					print( "done" );
				}
			}
		}
	}
}

### PICRUSt2-predicted pathways
# whole data, identifying pathways characteristic for leaves, roots or soils

picrust_pathways.nzv <- nearZeroVar( picrust_pathways )
picrust_pathways.lv <- picrust_pathways[ , -picrust_pathways.nzv$Position ]
picrust_pathways.lv.material.clr <- aldex.clr( as.data.frame(t(round(picrust_pathways.lv))), sdata.rarefied$Material, mc.samples=128, useMC=T )
picrust_pathways.lv.material.aldex <- aldex.kw(picrust_pathways.lv.material.clr, useMC=T )
picrust_pathways.lv.material.aldex.significant <- picrust_pathways.lv.material.aldex[ picrust_pathways.lv.material.aldex$kw.eBH < 0.05 | picrust_pathways.lv.material.aldex$glm.eBH < 0.05, ]
picrust_pathways.lv.material.aldex.significant <- merge( picrust_pathways.lv.material.aldex.significant, pathways_descriptions, by.x=0, by.y=0, all.x=T )
picrust_pathways.lv.material.aldex.significant <- picrust_pathways.lv.material.aldex.significant[ order(picrust_pathways.lv.material.aldex.significant$kw.eBH, decreasing=T), ]
write.table( picrust_pathways.lv.material.aldex.significant, "picrust_pathways.lv.material.aldex.significant.csv", sep="\t", col.names=NA )

# data divided according to material, identifying pathways characteristic for late and early samples

for( material in c('L','R', 'S') ){
	asvtable <- paste0("picrust_pathways.", material);
	sampledata <- paste0("sdata.", material);
	asvtablenzv <- paste(asvtable, ".nzv");
	asvtablelv <- paste0(asvtable, '.lv');
	clr <- paste0(asvtablelv, ".clr");
	ald <- paste0(asvtablelv, ".aldex");
	aldsign <- paste0(ald, ".significant");
	file <- paste0(aldsign, ".csv");
	assign( asvtable, picrust_pathways[ sdata.rarefied$Material == material, ] );
	assign( sampledata, sdata.rarefied[ sdata.rarefied$Material == material, ] );
	assign( asvtablenzv, nearZeroVar(get(asvtable)) );
	assign( asvtablelv, get(asvtable)[ , -get(asvtablenzv)$Position ] );
	assign( clr, aldex.clr(as.data.frame(t(round(get(asvtablelv)))), get(sampledata)$status, mc.samples=128, useMC=T) );
	assign( ald, aldex.kw(get(clr), useMC=T) );
	assign( aldsign, get(ald)[ get(ald)$kw.eBH < 0.05 | get(ald)$glm.eBH < 0.05, ] );
	assign( aldsign, merge(get(aldsign), pathways_descriptions, by.x=0, by.y=0, all.x=T) );
	assign( aldsign, get(aldsign)[ order(get(aldsign)$kw.eBH, decreasing=T), ] );
	write.table( get(aldsign), file, sep="\t", col.names=NA );
}

# data divided according to material * soil * genotype, identifying patheways characteristic for late and early as well as inoculated and non-inoculated samples

for( material in c( 'S', 'R', 'L' ) ){
	for( soil in c( 'S1', 'S2' ) ){
		for( genotype in c( 'B', 'C', 'M' ) ){
			asvtable <- as.data.frame(round(picrust_pathways[ sdata.rarefied$Material == material & sdata.rarefied$Soil == soil & sdata.rarefied$Genotype == genotype, ]));
			sampledata <- sdata.rarefied[ sdata.rarefied$Material == material & sdata.rarefied$Soil == soil & sdata.rarefied$Genotype == genotype, ];
			nzv <- nearZeroVar( asvtable );
			lv <- asvtable[ , -nzv$Position ];
			ald <- paste0("picrust_pathways.", material, ".", soil, ".", genotype, ".", "status.aldex");
			file <- paste0( ald, ".csv" );
			aldsign <- paste0("picrust_pathways.", material, ".", soil, ".", genotype, ".", "status.aldex.significant");
			if(!file.exists(file) & ncol(lv) > 0 & length(levels(as.factor(sampledata$status))) == 2){
				print( ald );
				clr <- aldex.clr( t(lv), sampledata$status, mc.samples=128, useMC=T );
				assign( ald, aldex.kw( clr, useMC=T ) );
				assign( aldsign, get(ald)[ get(ald)$kw.eBH < 0.05 | get(ald)$glm.eBH < 0.05, ] );
				assign( aldsign, merge(get(aldsign), pathways_descriptions, by=0, all.x=T) );
				assign( aldsign, get(aldsign)[ order(get(aldsign)$kw.eBH, decreasing=T), ] );
				write.table(get(aldsign), file, col.names=NA, sep="\t");
				print( "done" );
			}

			for( status in c( 'early', 'late' ) ){
				asvtable <- as.data.frame(round(picrust_pathways[ sdata.rarefied$Material == material & sdata.rarefied$Soil == soil & sdata.rarefied$Genotype == genotype & sdata.rarefied$status == status, ]));
				sampledata <- sdata.rarefied[ sdata.rarefied$Material == material & sdata.rarefied$Soil == soil & sdata.rarefied$Genotype == genotype & sdata.rarefied$status == status, ];
				nzv <- nearZeroVar( asvtable );
				lv <- asvtable[ , -nzv$Position ];
				ald <- paste0("picrust_pathways.", material, ".", soil, ".", genotype, ".", status, ".Inoculation.aldex");
				file <- paste0( ald, ".csv" );
				aldsign <- paste0("picrust_pathways.", material, ".", soil, ".", genotype, ".", status, ".Inoculation.aldex.significant");
				if(!file.exists(file) & ncol(lv) > 0 & length(levels(as.factor(sampledata$Inoculation))) == 2){
					print( ald );
					clr <- aldex.clr( t(lv), sampledata$Inoculation, mc.samples=128, useMC=T );
					assign( ald, aldex.kw( clr, useMC=T ) );
					assign( aldsign, get(ald)[ get(ald)$kw.eBH < 0.05 | get(ald)$glm.eBH < 0.05, ] );
					assign( aldsign, merge(get(aldsign), pathways_descriptions, by=0, all.x=T) );
					assign( aldsign, get(aldsign)[ order(get(aldsign)$kw.eBH, decreasing=T), ] );
					write.table(get(aldsign), file, col.names=NA, sep="\t");
					print( "done" );
				}
			}
			for( inoculation in c( 'N', 'I' ) ){
				asvtable <- as.data.frame(round(picrust_pathways[ sdata.rarefied$Material == material & sdata.rarefied$Soil == soil & sdata.rarefied$Genotype == genotype & sdata.rarefied$Inoculation == inoculation, ]));
				sampledata <- sdata.rarefied[ sdata.rarefied$Material == material & sdata.rarefied$Soil == soil & sdata.rarefied$Genotype == genotype & sdata.rarefied$Inoculation == inoculation, ];
				nzv <- nearZeroVar( asvtable );
				lv <- asvtable[ , -nzv$Position ];
				ald <- paste0("picrust_pathways.", material, ".", soil, ".", genotype, ".", inoculation, ".LateEarly.aldex");
				file <- paste0( ald, ".csv" );
				aldsign <- paste0("picrust_pathways.", material, ".", soil, ".", genotype, ".", inoculation, ".LateEarly.aldex.significant");
				if(!file.exists(file) & ncol(lv) > 0 & length(levels(as.factor(sampledata$status))) == 2){
					print( ald );
					clr <- aldex.clr( reads=t(lv), sampledata$status, mc.samples=128, useMC=T );
					assign( ald, aldex.kw( clr, useMC=T ) );
					assign( aldsign, get(ald)[ get(ald)$kw.eBH < 0.05 | get(ald)$glm.eBH < 0.05, ]);
					assign( aldsign, merge(get(aldsign), pathways_descriptions, by=0, all.x=T) );
					assign( aldsign, get(aldsign)[ order(get(aldsign)$kw.eBH, decreasing=T), ] );
					write.table(get(aldsign), file, col.names=NA, sep="\t");
					print( "done" );
				}
			}
		}
	}
}
```

**SM7m.** Identification of differentially abundant ASVs and taxa with DESeq2

```
library(mixOmics)
library(DESeq2)
library(BiocParallel)

### amplicon data
# whole data – identifying ASVs characteristic for leaves, roots or soils

bacteria.nzv <- nearZeroVar( bacteria )
bacteria.lv <- bacteria[ , -bacteria.nzv$Position ]
bacteria.lv.dds <- DESeqDataSetFromMatrix( countData=as.matrix(t(bacteria.lv)+1), colData=sdata, design= ~ Material )
bacteria.lv.dds <- DESeq( bacteria.lv.dds, parallel=T, BPPARAM=MulticoreParam(30) )
bacteria.lv.dds.resultsLR <- results( bacteria.lv.dds, contrast=c("Material", "L", "R"), parallel=T, BPPARAM=MulticoreParam(30) )
bacteria.lv.dds.resultsLR <- bacteria.lv.dds.resultsLR[ !is.na(bacteria.lv.dds.resultsLR$padj) & bacteria.lv.dds.resultsLR$padj < 0.05 & abs(bacteria.lv.dds.resultsLR$log2FoldChange) >= 1 , ]
bacteria.lv.dds.resultsLR.df <- merge( as.data.frame(bacteria.lv.dds.resultsLR), bacteria.tax, by.x=0, by.y=0, all.x=T )
bacteria.lv.dds.resultsLR.df <- bacteria.lv.dds.resultsLR.df[ order(bacteria.lv.dds.resultsLR.df$log2FoldChange, decreasing=T), ]
write.table( bacteria.lv.dds.resultsLR.df, "bacteria.lv.dds.resultsLR.csv", sep="\t", col.names=NA )
bacteria.lv.dds.resultsLS <- results( bacteria.lv.dds, contrast=c("Material", "L", "S"), parallel=T, BPPARAM=MulticoreParam(30) )
bacteria.lv.dds.resultsLS <- bacteria.lv.dds.resultsLS[ !is.na(bacteria.lv.dds.resultsLS$padj) & bacteria.lv.dds.resultsLS$padj < 0.05 & abs(bacteria.lv.dds.resultsLS$log2FoldChange) >= 1, ]
bacteria.lv.dds.resultsLS.df <- merge( as.data.frame(bacteria.lv.dds.resultsLS), bacteria.tax, by.x=0, by.y=0, all.x=T )
bacteria.lv.dds.resultsLS.df <- bacteria.lv.dds.resultsLS.df[ order(bacteria.lv.dds.resultsLS.df$log2FoldChange, decreasing=T), ]
write.table( bacteria.lv.dds.resultsLS.df, "bacteria.lv.dds.resultsLS.csv", sep="\t", col.names=NA )
bacteria.lv.dds.resultsSR <- results( bacteria.lv.dds, contrast=c("Material", "S", "R"), parallel=T, BPPARAM=MulticoreParam(30) )
bacteria.lv.dds.resultsSR <- bacteria.lv.dds.resultsSR[ !is.na(bacteria.lv.dds.resultsSR$padj) & bacteria.lv.dds.resultsSR$padj < 0.05 & abs(bacteria.lv.dds.resultsSR$log2FoldChange) >= 1, ]
bacteria.lv.dds.resultsSR.df <- merge( as.data.frame(bacteria.lv.dds.resultsSR), bacteria.tax, by.x=0, by.y=0, all.x=T )
bacteria.lv.dds.resultsSR.df <- bacteria.lv.dds.resultsSR.df[ order(bacteria.lv.dds.resultsSR.df$log2FoldChange, decreasing=T), ]
write.table( bacteria.lv.dds.resultsSR.df, "bacteria.lv.dds.resultsSR.csv", sep="\t", col.names=NA )

# data divided according to material – identifying ASVs characteristic for late and early samples
for( material in c('L', 'R', 'S') ){
	asvtable <- paste0("bacteria.", material);
	sampledata <- paste0("sdata.", material);
	asvtablenzv <- paste(asvtable, ".nzv");
	asvtablelv <- paste0(asvtable, '.lv');
	dds <- paste0(asvtablelv, ".status.dds");
	dds.results <- paste0(dds, ".results");
	dds.results.df <- paste0(dds.results, ".df");
	file <- paste0(dds.results, ".csv");
	assign( asvtable, bacteria[ sdata$Material == material, ] );
	assign( sampledata, sdata[ sdata$Material == material, ] );
	assign( asvtablenzv, nearZeroVar(get(asvtable)) );
	assign( asvtablelv, get(asvtable)[ , -get(asvtablenzv)$Position ] );
	assign( dds, DESeqDataSetFromMatrix(countData=as.matrix(t(get(asvtablelv)+1)), colData=get(sampledata), design= ~ status) );
	assign( dds, DESeq(get(dds), parallel=T, BPPARAM=MulticoreParam(30)) );
	assign( dds.results, results(get(dds), contrast=c("status", "early", "late"), parallel=T, BPPARAM=MulticoreParam(30)) );
	assign( dds.results, get(dds.results)[ !is.na(get(dds.results)$padj) & get(dds.results)$padj < 0.05 & abs(get(dds.results)$log2FoldChange) >= 1, ] );
	assign( dds.results.df, merge(as.data.frame(get(dds.results)), bacteria.tax, by.x=0, by.y=0, all.x=T) );
	assign( dds.results.df, get(dds.results.df)[ order(get(dds.results.df)$log2FoldChange, decreasing=T) , ] );
	write.table( get(dds.results.df), file, sep="\t", col.names=NA );
}

# data divided according to material * soil * genotype, identifying ASVs characteristic for late and early as well as inoculated and non-inoculated samples


### PICRUSt2-predicted functions
# whole data, identifying functions characteristic for leaves, roots or soils


# data divided according to material, identifying functions characteristic for late and early samples


# data divided according to material * soil * genotype, identifying functions characteristic for late and early as well as inoculated and non-inoculated samples


### PICRUSt2-predicted pathways
# whole data, identifying pathways characteristic for leaves, roots or soils


# data divided according to material, identifying pathways characteristic for late and early samples


# data divided according to material * soil * genotype, identifying patheways characteristic for late and early as well as inoculated and non-inoculated samples
```

**SM7n.** ASV-specific primer design with DECIPHER

The analysis was carried out on data without inoculant generated as described in SM (pay attention to a difference in fasta file name).

In bash:

rename_asvs_to_otus.perl -f seqs1.fasta -n Otu

In R:

```
library(DECIPHER)
fas <- "seqs1.renamed.fasta"
dbConn <- dbConnect(SQLite(), "asv_db.sqlite")
Seqs2DB(fas, "FASTA", dbConn, "ASV")
desc <- dbGetQuery(dbConn, "select description from Seqs")
Add2DB( data.frame(identifier=desc, stringsAsFactors=F), dbConn)
ids <- data.frame(identifier=desc, stringsAsFactors=F)
Add2DB(ids, dbConn)
colnames(ids) <- c("identifier")
Add2DB(ids, dbConn)
tiles <- TileSeqs(dbConn, add2tbl="Tiles", minCoverage=1)
saveRDS(tiles, "ASVs_tiles.rds", compress=T)

# Different ASVs were chosen, finally primers for ASV10 (Otu0008 here) were used
for( otu in c('Otu00008', 'Otu00057', 'Otu001347', 'Otu00695', 'Otu00010', 'Otu00024', 'Otu00040', 'Otu00498', 'Otu00259', 'Otu00232', 'Otu00046', 'Otu00706', 'Otu00063') ){
result <- paste0('primersASV', sub('Otu', '', otu));
file <- paste0(result, '.csv');
print( result );
assign( result, DesignPrimers(tiles=tiles, minLength=22, divalent=1.5, monovalent=50, dNTPs=0.8, minProductSize=120, maxProductSize=150, minEfficiency=0.95, identifier=otu, numPrimerSets=5, processors=4, minCoverage=1, minGroupCoverage=1) );
write.table( get(result), file, sep='\t')
}
```

**SM7o.** Nestedness analysis

In R:

```
# generation of data for NODF
library(dada2)
library(vegan)

leaves.seqtab <- readRDS('betamikro_WP5_leaves.rds')
roots.seqtab <- readRDS('betamikro_WP5_roots.rds')
soils.seqtab <- readRDS('betamikro_WP5_soils.rds')
WP5_complete.seqtab <- mergeSequenceTables(leaves.seqtab, roots.seqtab, soils.seqtab)
soils.seqtab <- soils.seqtab[ -grep('KN', rownames(soils.seqtab)), ]
roots.seqtab <- roots.seqtab[ -grep('KN', rownames(roots.seqtab)), ]
leaves.seqtab <- leaves.seqtab[ -grep('KN', rownames(leaves.seqtab)), ]
WP5_complete.seqtab <- mergeSequenceTables(leaves.seqtab, roots.seqtab, soils.seqtab)
WP5_complete.taxa <- assignTaxonomy(WP5_complete.seqtab, "~/db/silva/v138/silva_nr99_v138.1_train_set.fa.gz", multithread=T)
WP5_complete.taxa <- as.data.frame( WP5_complete.taxa)

WP5_complete.taxa <- WP5_complete.taxa[ WP5_complete.taxa$Kingdom == 'Bacteria', ]
WP5_complete.taxa <- WP5_complete.taxa[ WP5_complete.taxa$Order != 'Chloroplast', ]
WP5_complete.taxa <- WP5_complete.taxa[ WP5_complete.taxa$Family != 'Mitochondria', ]

WP5_complete.seqtab.bacteria <- WP5_complete.seqtab[ , colnames(WP5_complete.seqtab) %in% rownames(WP5_complete.taxa) ]
seqs1.fasta <- data.frame(seq=colnames(WP5_complete.seqtab.bacteria), row.names=paste(">ASV", seq(1, dim(WP5_complete.seqtab.bacteria)[2], 1), "\n", sep=""))
write.table(seqs1.fasta, "seqs1.fasta", quote=F, sep="", col.names=F)

seqs1.count_table <- as.data.frame(t(WP5_complete.seqtab.bacteria))
rownames(seqs1.count_table) <- paste(">ASV", seq(1, dim(WP5_complete.seqtab.bacteria)[2], 1), "\n", sep="")
seqs1.count_table$total <- rowSums(seqs1.count_table)
seqs1.count_table <- seqs1.count_table[, c(ncol(seqs1.count_table), 1:(ncol(seqs1.count_table)-1))]
seqs1.count_table$Representative_sequence <- paste("ASV", seq(1, dim(WP5_complete.seqtab.bacteria)[2], 1), sep="")
seqs1.count_table <- seqs1.count_table[, c(ncol(seqs1.count_table), 1:(ncol(seqs1.count_table)-1))]
write.table(seqs1.count_table, "seqs1.count_table", quote=F, sep="\t", row.names=F)

sdata <- read.table("WP5.design.csv", header=T, sep=",")
rownames(sdata) <- sdata$sample
sdata$Sample <- gsub("G|K|L", "", rownames(sdata))
sdata$Material <- substr(rownames(sdata), 1,1)
sdata$Material <- gsub("G", "S", sdata$Material)
sdata$Material <- gsub("K", "R", sdata$Material)
sdata <- sdata[ rownames(sdata) %in% rownames(WP5_complete.seqtab.bacteria),  ]
WP5_complete.seqtab.bacteria <- WP5_complete.seqtab.bacteria[ rownames(WP5_complete.seqtab.bacteria) %in% rownames(sdata), ] # only those samples, for which there are data
sdata <- sdata[ rownames(sdata) %in% rownames(WP5_complete.seqtab.bacteria), ]
WP5_complete.seqtab.bacteria <- WP5_complete.seqtab.bacteria[ sort(rownames(WP5_complete.seqtab.bacteria)), ]
sdata <- sdata[ sort(rownames(sdata)), ]
identical(rownames(WP5_complete.seqtab.bacteria), rownames(sdata)) # must be TRUE

colnames( WP5_complete.seqtab.bacteria) <- paste0("ASV", seq(1, dim(WP5_complete.seqtab.bacteria)[2], 1) )
WP5_complete.seqtab.bacteria.rarefied <- Rarefy(WP5_complete.seqtab.bacteria, depth=500)$otu.tab.rff
sdata.rarefied <- sdata[ rownames(sdata) %in% rownames(WP5_complete.seqtab.bacteria.rarefied), ]

for( sample in 1:781 ){
	if( length(rownames(WP5_complete.seqtab.bacteria.rarefied[ rownames(sdata.rarefied[sdata.rarefied$Sample == sample, ]), ])) > 2 ){
		print( rownames(sdata.rarefied[ sdata.rarefied$Sample == sample, ]) );
		name <- paste0(sdata.rarefied[sdata.rarefied$Sample == sample, ]$Soil[1], '_', sdata.rarefied[sdata.rarefied$Sample == sample, ]$Genotype[1], '_', sdata.rarefied[sdata.rarefied$Sample == sample, ]$Time[1], '_', sdata.rarefied[sdata.rarefied$Sample == sample, ]$Inoculation[1], '_', sdata.rarefied[sdata.rarefied$Sample == sample, ]$Biological_replicate[1], '_', sdata.rarefied[sdata.rarefied$Sample == sample, ]$Technical_replicate[1] );
		print( name );
		assign( name, t(WP5_complete.seqtab.bacteria.rarefied[rownames(sdata.rarefied[sdata.rarefied$Sample == sample, ]), ]) );
		assign( name, get(name)[ rowSums(get(name)) > 0, ] );
		write.table( get(name), paste0(name, ".txt"), quote=F, sep="\t", row.names=rownames(get(name)) );
	}
}
```

In Windows:

\# NODF running script – data generated above need to be transferred to a Windows machine on which NODF is to be run

In R

```
# graphs preparation
nodf <- read.table("NODF.csv", header=T, sep="\t", dec=',', row.names=1)
nodf$status <- 'early'
nodf$status[ nodf$Time == 'T3' | nodf$Time == 'T4' ] <- 'late'

svg( "bacteria_nodf_time.svg", width=3.5, height=3.5, pointsize=6 );
p1 <- ggplot( nodf, aes_(x=nodf$Time, y=nodf$NODF)) + geom_boxplot() + facet_wrap( ~ nodf$Soil * nodf$Genotype) + scale_x_discrete(drop=T) + xlab("Timepoint") + ylab("Nestedness (weighted NODF)")
print(p1)
dev.off()


svg( "bacteria_nodf_inoculation.svg", width=3.5, height=3.5, pointsize=6 );
p1 <- ggplot( nodf, aes_(x=nodf$status, y=nodf$NODF, fill=nodf$Inoculation)) + geom_boxplot() + facet_wrap( ~ nodf$Soil * nodf$Genotype) + scale_x_discrete(drop=T) + xlab("Status") + ylab("Nestedness (weighted NODF)") + labs(fill="")
print(p1)
dev.off()
```

**SM7p.** Statistical analyses

R script:

```
library(Rfit)
library(dunn.test)
# assessment of non-mitochondrial and non-plastidic 16S rRNA gene sequences in seeds and seedlings
# B335f-B769r (Dorn-In et al. 2015) - a pair amplifying bacterial 16S rRNA sequences, excluding mitochondrial and plastidic ones
bac335_curve <- read.table("bacteria_standard_curve_kit2022.csv", header=T, sep="\t", dec=',')
bac335_curve$mean_ct <- rowMeans(bac335_curve[ , 3:6 ], na.rm=T)
bac335_curve$log_num_mol <- log(bac335_curve$num_mol, base=10)
bac335_curve.lm <- lm( mean_ct ~ log_num_mol, data=bac335_curve)
summary(bac335_curve.lm)
qpcr_antibiotics <- read.table("antibiotics_qPCR_results.csv", header=T, sep="\t", dec=",")
qpcr_antibiotics$bac335_mean_ct <- rowMeans( as.data.frame(qpcr_antibiotics[ , 26:28 ]), na.rm=T )
qpcr_antibiotics$bac335_num_mol <- 10^((qpcr_antibiotics$bac335_mean_ct - bac335_curve.lm$coefficients[1])/bac335_curve.lm$coefficients[2] )
qpcr_antibiotics$bac335_num_mol[ qpcr_antibiotics$bac335_mean_ct == 45 ] <- 0 # values below detection limit (last standard Ct=45.385)
qpcr_antibiotics_seeds <- qpcr_antibiotics[ qpcr_antibiotics$material == 'seeds', ]
qpcr_antibiotics_passages <- qpcr_antibiotics[ qpcr_antibiotics$material == 'plantlets', ]
qpcr_antibiotics_seedlings <- qpcr_antibiotics[ qpcr_antibiotics$material == 'seedlings', ]
# Fig. SR2 A
svg( "bac335_seeds.svg", width=3.5, height=3.5, pointsize=6 );
p3 <- ggplot( qpcr_antibiotics_seeds, aes_(x=qpcr_antibiotics_seeds$genotype, y=qpcr_antibiotics_seeds$bac335_num_mol, fill=qpcr_antibiotics_seeds$variant)) + geom_bar(stat='identity', position=position_dodge()) + scale_x_discrete(drop=T) + xlab("Genotype") + ylab("16S rRNA counts/ng of DNA") + labs(fill="Variant")
print(p3)
dev.off()
# Fig. SR2 B
svg( "bac335_seedlings.svg", width=3.5, height=3.5, pointsize=6 );
p3 <- ggplot( qpcr_antibiotics_seedlings, aes_(x=qpcr_antibiotics_seedlings$genotype, y=qpcr_antibiotics_seedlings$bac335_num_mol, fill=qpcr_antibiotics_seedlings$variant)) + geom_bar(stat='identity', position=position_dodge()) + scale_x_discrete(drop=T) + xlab("Genotype") + ylab("16S rRNA counts/ng of DNA") + labs(fill="Variant")
print(p3)
dev.off()

# correlation of ASV10 abundance in sequencing and qPCR results
bacteria.rarefied <- as.data.frame(bacteria.rarefied)
ASV10 <- data.frame( sample=rownames(bacteria.rarefied), ASV10=bacteria.rarefied$ASV10, row.names=rownames(bacteria.rarefied) )
asv10_curve <-read.table("bacteria_ASV8_standard_curve.csv", header=T, sep="\t", dec=",")
asv10_curve$mean_ct <- rowMeans(asv10_curve[ , 3:6 ], na.rm=T)
asv10_curve$log_num_mol <- log(asv10_curve$num_mol, base=10)
asv10_curve.lm <- lm( mean_ct ~ log_num_mol, data=asv10_curve)
summary(asv10_curve.lm)
# Roots
ASV10.R <- ASV10[ grep("K", rownames(ASV10)), ] # roots
qpcr_asv10_roots <- read.table("qPCR_ASV8_roots.csv", header=T, sep="\t", dec=",")
rownames(qpcr_asv10_roots) <- qpcr_asv10_roots$sample
qpcr_asv10_roots$mean_ct <- rowMeans( as.data.frame(qpcr_asv10_roots[ , 8:11 ]), na.rm=T )
qpcr_asv10_roots$num_mol <- 10^((qpcr_asv10_roots$mean_ct - asv10_curve.lm$coefficients[1])/asv10_curve.lm$coefficients[2])
qpcr_asv10_roots$num_mol[ qpcr_asv10_roots$mean_ct >= 27 ] <- 0
qpcr_asv10_roots <- qpcr_asv10_roots[ rownames(qpcr_asv10_roots) %in% rownames(ASV10.R), ]
qpcr_asv10_roots <- qpcr_asv10_roots[ order(rownames(qpcr_asv10_roots)), ]
ASV10.R <- ASV10.R[ rownames(ASV10.R) %in% rownames(qpcr_asv10_roots), ]
ASV10.R <- ASV10.R[ order(rownames(ASV10.R)), ]
identical( rownames(ASV10.R), rownames(qpcr_asv10_roots) ) # must be true
qpcr_vs_sequencing_spearman.R.cor <- rcorr( ASV10.R$ASV10, qpcr_asv10_roots$num_mol, type='spearman' )
ASV10.R.lm <- lm( ASV10.R$ASV10 ~ qpcr_asv10_roots$num_mol )
summary( ASV10.R.lm )
# Leaves
ASV10.L <- ASV10[ grep("L", rownames(ASV10)), ] # leaves
qpcr_asv10_leaves <- read.table("qPCR_ASV8_leaves.csv", header=T, sep="\t", dec=",")
rownames(qpcr_asv10_leaves) <- qpcr_asv10_leaves$sample
qpcr_asv10_leaves$mean_ct <- rowMeans( as.data.frame(qpcr_asv10_leaves[ , 8:11 ]), na.rm=T )
qpcr_asv10_leaves$num_mol <- 10^((qpcr_asv10_leaves$mean_ct - asv10_curve.lm$coefficients[1])/asv10_curve.lm$coefficients[2])
qpcr_asv10_leaves$num_mol[ qpcr_asv10_leaves$mean_ct >= 27 ] <- 0
qpcr_asv10_leaves <- qpcr_asv10_leaves[ rownames(qpcr_asv10_leaves) %in% rownames(ASV10.L), ]
qpcr_asv10_leaves <- qpcr_asv10_leaves[ order(rownames(qpcr_asv10_leaves)), ]
ASV10.L <- ASV10.L[ rownames(ASV10.L) %in% rownames(qpcr_asv10_leaves), ]
ASV10.L <- ASV10.L[ order(rownames(ASV10.L)), ]
identical( rownames(ASV10.L), rownames(qpcr_asv10_leaves) ) # must be true
qpcr_vs_sequencing_spearman.L.cor <- rcorr( ASV10.L$ASV10, qpcr_asv10_leaves$num_mol, type='spearman' )
ASV10.L.lm <- lm( ASV10.L$ASV10 ~ qpcr_asv10_leaves$num_mol )
summary( ASV10.L.lm )

### Bacterial load analysis
# Roots
acterial_load_roots <- read.table("qPCR_bacteria_roots.csv", header=T, sep="\t", dec=',')
rownames(bacterial_load_roots) <- bacterial_load_roots$sample
bacterial_load_curve2022 <-read.table("bacteria_standard_curve_kit2022.csv", header=T, sep="\t", dec=",")
bacterial_load_curve2022$mean_ct <- rowMeans(bacterial_load_curve2022[ , 3:6 ], na.rm=T)
bacterial_load_curve2022$log_num_mol <- log(bacterial_load_curve2022$num_mol, base=10)
bacterial_load_curve2022.lm <- lm( mean_ct ~ log_num_mol, data=bacterial_load_curve2022)
summary(bacterial_load_curve2022.lm)

bacterial_load_roots$mean_ct <- rowMeans(as.data.frame(bacterial_load_roots[ , 8:11]), na.rm=T)
bacterial_load_roots$num_mol <- 10^((bacterial_load_roots$mean_ct - bacterial_load_curve2022.lm$coefficients[1])/bacterial_load_curve2022.lm$coefficients[2])
bacterial_load_roots$num_mol[ bacterial_load_roots$mean_ct >= 39 ] <- NA # last standard's mean Ct was 45.12, but the samples were run for 45 cycles only and there is no Cts beyond 40th cycle
bacterial_load_roots$material <- 'R'

svg( "bacterial_load_roots_timepoint.svg", width=1.75, height=1.75, pointsize=3 ); # Fig. 3C
p1 <- ggplot( bacterial_load_roots, aes_(x=bacterial_load_roots$time, y=bacterial_load_roots$num_mol)) + geom_boxplot() + scale_x_discrete(drop=T) + xlab("Timepoint") + ylab("seqs/ng of DNA") + ggtitle("Load")
print(p1)
dev.off()

bacterial_load_roots.raov <- raov( num_mol ~ soil * genotype * time, data=bacterial_load_roots)
kruskal.test(bacterial_load_roots$num_mol, bacterial_load_roots$inoculation)
aggregate( bacterial_load_roots$num_mol ~ bacterial_load_roots$inoculation, FUN='mean')

# Leaves
bacterial_load_leaves <- read.table("qPCR_bacteria_leaves.csv", header=T, sep="\t", dec=',')
rownames(bacterial_load_leaves) <- bacterial_load_leaves$sample
bacterial_load_leaves$mean_ct <- rowMeans(as.data.frame(bacterial_load_leaves[ , 8:11]), na.rm=T)
bacterial_load_leaves$num_mol <- 10^((bacterial_load_leaves$mean_ct - bacterial_load_curve2022.lm$coefficients[1])/bacterial_load_curve2022.lm$coefficients[2])
bacterial_load_leaves$num_mol[ bacterial_load_leaves$mean_ct >= 45 ] <- NA # last standard's mean Ct was 45.12
bacterial_load_leaves$material <- 'L'

svg( "bacterial_load_leaves_timepoint.svg", width=1.75, height=1.75, pointsize=3 ); # Fig. 3C
p1 <- ggplot( bacterial_load_leaves, aes_(x=bacterial_load_leaves$time, y=bacterial_load_leaves$num_mol)) + geom_boxplot() + scale_x_discrete(drop=T) + xlab("Timepoint") + ylab("seqs/ng of DNA") + ggtitle("Load")
print(p1)
dev.off()

bacterial_load_leaves.raov <- raov( num_mol ~ soil * genotype * time, data=bacterial_load_leaves)
kruskal.test(bacterial_load_leaves$num_mol, bacterial_load_leaves$inoculation)
aggregate( bacterial_load_leaves$num_mol ~ bacterial_load_leaves$inoculation, FUN='mean')


bacterial_load <- rbind(bacterial_load_roots, bacterial_load_leaves)

svg( "bacterial_load_material.svg", width=1.75, height=1.75, pointsize=3 ) # Fig. 3
p1 <- ggplot( bacterial_load, aes_(x=bacterial_load$material, y=bacterial_load$num_mol) ) + geom_boxplot() + scale_x_discrete(drop=T) + xlab("Material") + ylab("seqs/ng of DNA") + ggtitle("Load")
print(p1)
dev.off()

# Inoculant
bacterial_load_inoculant <- read.table("qPCR_bacteria_inoculant.csv", header=T, sep="\t", dec=',')
rownames(bacterial_load_inoculant) <- bacterial_load_inoculant$sample
bacterial_load_inoculant$mean_ct <- rowMeans(as.data.frame(bacterial_load_inoculant[ , 2:4]), na.rm=T)
bacterial_load_inoculant$num_mol <- 10^((bacterial_load_inoculant$mean_ct - bacterial_load_curve2022.lm$coefficients[1])/bacterial_load_curve2022.lm$coefficients[2])
bacterial_load_inoculant$num_mol[ bacterial_load_inoculant$mean_ct >= 45 ] <- NA # last standard's mean Ct was 45.12
bacterial_load_inoculant$num_mol_per_g <- bacterial_load_inoculant$num_mol * bacterial_load_inoculant$volume * bacterial_load_inoculant$concentration/bacterial_load_inoculant$weight # number of copies per g of material = number of copies per ng of DNA * number of ng/g of material (concentration * volume/weight of material in g)
mean(bacterial_load_inoculant$num_mol)
sd(bacterial_load_inoculant$num_mol)
mean(bacterial_load_inoculant$num_mol_per_g)
sd(bacterial_load_inoculant$num_mol_per_g)

# Alpha-diversity of amplicons
library(lawstat)
library(Rfit)
# Testing differences in alpha-diversity
shapiro.test(bacteria.rarefied.diversity$sobs) # significant deviation from normality, non-parametric test needs to be used
shapiro.test(bacteria.rarefied.diversity$H) # significant deviation from normality, non-parametric test needs to be used
shapiro.test(bacteria.rarefied.diversity$E) # significant deviation from normality, non-parametric test needs to be used

sobs.raov <- raov(sobs ~ material * soil * genotype * time, data=bacteria.rarefied.diversity)
summary(sobs.raov)
sobs1.raov <- raov( sobs ~ material * soil * genotype * status * inoculation, data=bacteria.rarefied.diversity )
summary(sobs1.raov)

# Material (tested with oneway.rfit)
alpha_diversity_material <- as.data.frame( aggregate(H ~ material, data=bacteria.rarefied.diversity, FUN="mean") )
rownames(alpha_diversity_material) <- alpha_diversity_material$material
alpha_diversity_material$H_sd <- aggregate( H ~ material, data=bacteria.rarefied.diversity, FUN="sd" )$H
alpha_diversity_material$E <- aggregate( E ~ material, data=bacteria.rarefied.diversity, FUN="mean" )$E
alpha_diversity_material$E_sd <- aggregate( E ~ material, data=bacteria.rarefied.diversity, FUN="sd" )$E
alpha_diversity_material$sobs <- aggregate( sobs ~ material, data=bacteria.rarefied.diversity, FUN="mean" )$sobs
alpha_diversity_material$sobs_sd <- aggregate( sobs ~ material, data=bacteria.rarefied.diversity, FUN="sd" )$sobs

alpha_diversity_material_H.rfit <- oneway.rfit(bacteria.rarefied.diversity$H, bacteria.rarefied.diversity$material)
alpha_diversity_material_E.rfit <- oneway.rfit(bacteria.rarefied.diversity$E, bacteria.rarefied.diversity$material)
alpha_diversity_material_sobs.rfit <- oneway.rfit(bacteria.rarefied.diversity$sobs, bacteria.rarefied.diversity$material)

# Soil (tested with wilcox.test)
alpha_diversity_soil <- as.data.frame( aggregate(H ~ soil, data=bacteria.rarefied.diversity, FUN="mean") )
rownames(alpha_diversity_soil) <- alpha_diversity_soil$soil
alpha_diversity_soil$H_sd <- aggregate( H ~ soil, data=bacteria.rarefied.diversity, FUN="sd" )$H
alpha_diversity_soil$E <- aggregate( E ~ soil, data=bacteria.rarefied.diversity, FUN="mean" )$E
alpha_diversity_soil$E_sd <- aggregate( E ~ soil, data=bacteria.rarefied.diversity, FUN="sd" )$E
alpha_diversity_soil$sobs <- aggregate( sobs ~ soil, data=bacteria.rarefied.diversity, FUN="mean" )$sobs
alpha_diversity_soil$sobs_sd <- aggregate( sobs ~ soil, data=bacteria.rarefied.diversity, FUN="sd" )$sobs

alpha_diversity_soil_H.wilcox <- wilcox.test( H ~ soil, data=bacteria.rarefied.diversity )
alpha_diversity_soil_E.wilcox <- wilcox.test( E ~ soil, data=bacteria.rarefied.diversity )
alpha_diversity_soil_sobs.wilcox <- wilcox.test( sobs ~ soil, data=bacteria.rarefied.diversity )

# Genotype (tested with oneway.rfit)
alpha_diversity_genotype <- as.data.frame( aggregate(H ~ genotype, data=bacteria.rarefied.diversity, FUN="mean") )
rownames(alpha_diversity_genotype) <- alpha_diversity_genotype$genotype
alpha_diversity_genotype$H_sd <- aggregate( H ~ genotype, data=bacteria.rarefied.diversity, FUN="sd" )$H
alpha_diversity_genotype$E <- aggregate( E ~ genotype, data=bacteria.rarefied.diversity, FUN="mean" )$E
alpha_diversity_genotype$E_sd <- aggregate( E ~ genotype, data=bacteria.rarefied.diversity, FUN="sd" )$E
alpha_diversity_genotype$sobs <- aggregate( sobs ~ genotype, data=bacteria.rarefied.diversity, FUN="mean" )$sobs
alpha_diversity_genotype$sobs_sd <- aggregate( sobs ~ genotype, data=bacteria.rarefied.diversity, FUN="sd" )$sobs

alpha_diversity_genotype_H.rfit <- oneway.rfit(bacteria.rarefied.diversity$H, bacteria.rarefied.diversity$genotype)
alpha_diversity_genotype_E.rfit <- oneway.rfit(bacteria.rarefied.diversity$E, bacteria.rarefied.diversity$genotype)
alpha_diversity_genotype_sobs.rfit <- oneway.rfit(bacteria.rarefied.diversity$sobs, bacteria.rarefied.diversity$genotype)

# Time (tested with oneway.rfit)
alpha_diversity_time <- as.data.frame( aggregate(H ~ time, data=bacteria.rarefied.diversity, FUN="mean") )
rownames(alpha_diversity_time) <- alpha_diversity_time$time
alpha_diversity_time$H_sd <- aggregate( H ~ time, data=bacteria.rarefied.diversity, FUN="sd" )$H
alpha_diversity_time$E <- aggregate( E ~ time, data=bacteria.rarefied.diversity, FUN="mean" )$E
alpha_diversity_time$E_sd <- aggregate( E ~ time, data=bacteria.rarefied.diversity, FUN="sd" )$E
alpha_diversity_time$sobs <- aggregate( sobs ~ time, data=bacteria.rarefied.diversity, FUN="mean" )$sobs
alpha_diversity_time$sobs_sd <- aggregate( sobs ~ time, data=bacteria.rarefied.diversity, FUN="sd" )$sobs

alpha_diversity_time_H.rfit <- oneway.rfit(bacteria.rarefied.diversity$H, bacteria.rarefied.diversity$time)
alpha_diversity_time_E.rfit <- oneway.rfit(bacteria.rarefied.diversity$E, bacteria.rarefied.diversity$time)
alpha_diversity_time_sobs.rfit <- oneway.rfit(bacteria.rarefied.diversity$sobs, bacteria.rarefied.diversity$time)

# Status (tested with wilcox.test)
alpha_diversity_status <- as.data.frame( aggregate(H ~ status, data=bacteria.rarefied.diversity, FUN="mean") )
rownames(alpha_diversity_status) <- alpha_diversity_status$status
alpha_diversity_status$H_sd <- aggregate( H ~ status, data=bacteria.rarefied.diversity, FUN="sd" )$H
alpha_diversity_status$E <- aggregate( E ~ status, data=bacteria.rarefied.diversity, FUN="mean" )$E
alpha_diversity_status$E_sd <- aggregate( E ~ status, data=bacteria.rarefied.diversity, FUN="sd" )$E
alpha_diversity_status$sobs <- aggregate( sobs ~ status, data=bacteria.rarefied.diversity, FUN="mean" )$sobs
alpha_diversity_status$sobs_sd <- aggregate( sobs ~ status, data=bacteria.rarefied.diversity, FUN="sd" )$sobs

alpha_diversity_status_H.wilcox <- wilcox.test( H ~ status, data=bacteria.rarefied.diversity )
alpha_diversity_status_E.wilcox <- wilcox.test( E ~ status, data=bacteria.rarefied.diversity )
alpha_diversity_status_sobs.wilcox <- wilcox.test( sobs ~ status, data=bacteria.rarefied.diversity )

# Inoculation (tested with wilcox.test)
alpha_diversity_inoculation <- as.data.frame( aggregate(H ~ inoculation, data=bacteria.rarefied.diversity, FUN="mean") )
rownames(alpha_diversity_inoculation) <- alpha_diversity_inoculation$inoculation
alpha_diversity_inoculation$H_sd <- aggregate( H ~ inoculation, data=bacteria.rarefied.diversity, FUN="sd" )$H
alpha_diversity_inoculation$E <- aggregate( E ~ inoculation, data=bacteria.rarefied.diversity, FUN="mean" )$E
alpha_diversity_inoculation$E_sd <- aggregate( E ~ inoculation, data=bacteria.rarefied.diversity, FUN="sd" )$E
alpha_diversity_inoculation$sobs <- aggregate( sobs ~ inoculation, data=bacteria.rarefied.diversity, FUN="mean" )$sobs
alpha_diversity_inoculation$sobs_sd <- aggregate( sobs ~ inoculation, data=bacteria.rarefied.diversity, FUN="sd" )$sobs

alpha_diversity_inoculation_H.wilcox <- wilcox.test( H ~ inoculation, data=bacteria.rarefied.diversity )
alpha_diversity_inoculation_E.wilcox <- wilcox.test( E ~ inoculation, data=bacteria.rarefied.diversity )
alpha_diversity_inoculation_sobs.wilcox <- wilcox.test( sobs ~ inoculation, data=bacteria.rarefied.diversity )

### Analyis of culturable bacteria density in inoculant
culturable_bac_inoc <- read.table("culturable_bacteria_density_inoculant.csv", header=T, sep="\t", dec=",")
culturable_bac_inoc$cfu_per_g <- culturable_bac_inoc$colonies/culturable_bac_inoc$dry_weight/culturable_bac_inoc$dilution
mean(culturable_bac_inoc$cfu_per_g)
sd(culturable_bac_inoc$cfu_per_g)

### Analysis of inoculant microbiome's resistance to NaCl
library(reshape2)
library(ggplot2)
NaCl_resistance <- read.table("inoculant_resistance_to_NaCl.csv", sep="\t", dec=",", header=T)
rownames(NaCl_resistance) <- paste0(NaCl_resistance$sample, NaCl_resistance$plate)
NaCl_resistance$sample <- NULL
NaCl_resistance$plate <- NULL
NaCl_resistance_aggregated <- aggregate( . ~ inoculation, data=NaCl_resistance, FUN='mean')
NaCl_resistance_aggregated_molten <- melt(NaCl_resistance_aggregated, id.vars=c("inoculation"), value.name='OD600', variable.name='NaCl_concentration')
NaCl_resistance_aggregated_sd <- aggregate( . ~ inoculation, data=NaCl_resistance, FUN='sd')
NaCl_resistance_aggregated_sd_molten <- melt(NaCl_resistance_aggregated_sd, id.vars=c("inoculation"), value.name='OD600', variable.name='NaCl_concentration')
NaCl_resistance_aggregated_molten$SD <- NaCl_resistance_aggregated_sd_molten$OD600
NaCl_resistance_aggregated_molten$NaCl_concentration <- sub("X", "", NaCl_resistance_aggregated_molten$NaCl_concentration)
NaCl_resistance_aggregated_molten$NaCl_concentration <- sub("mM", "", NaCl_resistance_aggregated_molten$NaCl_concentration)
NaCl_resistance_aggregated_molten$NaCl_concentration <- factor(NaCl_resistance_aggregated_molten$NaCl_concentration, levels=c('0', '50', '100', '150', '200', '300', '400', '500', '600', '700', '800', '900'))
svg("inoculant_microbiome_NaCl_resistance.svg", width=3.5, height=3.5, pointsize=4) # Fig. 6B
p <- ggplot(NaCl_resistance_aggregated_molten, aes(x=NaCl_concentration, y=OD600, group=inoculation, color=inoculation)) + geom_line() + geom_point() + geom_errorbar(aes(ymin=OD600-SD, ymax=OD600+SD), width=0.2, position=position_dodge(0.05))
p <- p + theme_classic() + labs( x=expression(NaCl~concentration~(mM)), y=expression(OD[600]), color="" ) + theme( axis.text.x=element_text(angle=45, hjust=1))
print(p)
dev.off()

### Analysis of nestedness

# testing significance of differences between early and late samples – Table SR3

nodf_status_tests <- data.frame( matrix(ncol=3, nrow=0) )
for( soil in c('S1', 'S2') ){
	for( genotype in c('B', 'C', 'M') ){
		result <- paste0("nodf.", soil, ".", genotype, ".wilcoxon");
		chunk.early <- nodf[ nodf$Soil == soil & nodf$Genotype == genotype & nodf$status == 'early', ];
		chunk.late <- nodf[ nodf$Soil == soil & nodf$Genotype == genotype & nodf$status == 'late', ];
		assign( result, wilcox.test(chunk.early$NODF, chunk.late$NODF) );
		row <- c(soil, genotype, get(result)$p.value);
		nodf_status_tests <- rbind( nodf_status_tests, row );
	}
}
colnames(nodf_status_tests) <- c('Soil', 'Genotype', 'p.value')
nodf_status_tests$p.adj <- p.adjust( nodf_status_tests$p.value, method='BH' )

# testing significance of differences between inoculated and non-inoculated samples – Table SR

nodf_inoculation_tests <- data.frame( matrix(ncol=4, nrow=0) )
for( soil in c('S1', 'S2') ){
	for( genotype in c('B', 'C', 'M') ){
		for( status in c('early', 'late') ){
			result <- paste0("nodf.", soil, ".", genotype, ".", status, ".wilcoxon");
			chunk.I <- nodf[ nodf$Soil == soil & nodf$Genotype == genotype & nodf$status == status & nodf$Inoculation == 'I', ];
			chunk.N <- nodf[ nodf$Soil == soil & nodf$Genotype == genotype & nodf$status == status & nodf$Inoculation == 'N', ];
			assign( result, wilcox.test(chunk.I$NODF, chunk.N$NODF) );
			row <- c(soil, genotype, status, get(result)$p.value);
			nodf_inoculation_tests <- rbind( nodf_inoculation_tests, row );
		}
	}
}
colnames(nodf_inoculation_tests) <- c('Soil', 'Genotype', 'status', 'p.value')
nodf_inoculation_tests$p.adj <- p.adjust( nodf_inoculation_tests$p.value, method='BH' )
```
