## Supplementary Results for "Rhizosphere bacterial colonization of beet occurs in discrete phases regardless of bioinoculation with the wild sea beet root community"

**Regardless of bioinoculation rapid colonization of axenic beet plants by rhizosphere bacteria takes place in two phases**

Marcin Gołębiewski^1,2*^, Marcin Sikora^2^, Justyna Mazur^2^, Sonia Szymańska^3^, Jarosław Tyburski^1^, Katarzyna Hrynkiewicz^3^, Werner Ulrich^4^

**Soils of contrasting physico-chemical characteristics were used in experiments**

Soil S1 was more abundant in silt/clay and less abundant in nutrients than S2, which was more sandy (Table SR1).

**Table SR1.** Soils characteristics

|  | S1 | S2 |
| --- | --- | --- |
| Sand (%) | 60 | 74 |
| Silt (%) | 31 | 19 |
| Clay (%) | 9 | 7 |
| pH (H_2_O) | 7.450 | 7.560 |
| Total carbon (TC) (%) | 0.987 | 1.698 |
| Total organic carbon (TOC) (%) | 0.945 | 1.600 |
| Total inorganic carbon (TIC) (%) | 0.043 | 0.099 |
| Total nitrogen (TN) (%) | 0.220 | 0.269 |
| P (g/kg) | 0.662 | 0.700 |
| K (g/kg) | 3.299 | 2.101 |
| Na (g/kg) | 0.396 | 0.206 |
| Ca (g/kg) | 2.944 | 7.848 |
| Mg (g/kg) | 2.243 | 2.351 |

**Bacterial counts in surface-sterilized beet seeds are below detection limit**

Primer pair B335f-B769r (see Table SM1 and SM6a in Supplementary Methods) did not amplify plastidic nor mitochondrial sequences. Reactions were run as described in SM6a, but using 100 ng of a 27f-1492r amplicon generated on *Escherichia coli* DH10B as well as beet mitochondrial and chloroplastic 16S rRNA genes cloned into pCR4TOPO as templates. The reactions were run on a 1.5% agarose gel and visualized under UV illumination (Fig. SR1).

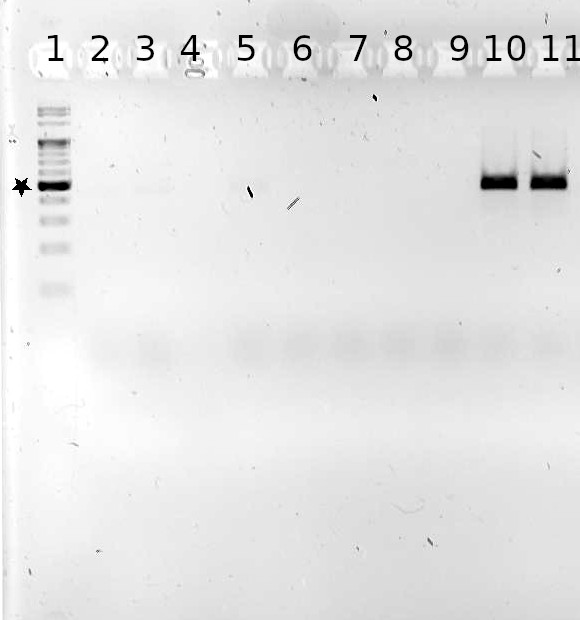

**Fig. SR1.** B335f-B769r primer pair does not amplify beet mitochondrial nor plastidic 16S rRNA gene fragments. 1.5% agarose gel, lanes: 1 – GeneRuler 100 bp Plus DNA ladder (ThermoFisher) (500 bp band is marked with a star), 2 – negative control (water), 3-5 – plastidic 16S rRNA gene, 6 – no template control (water), 7-9 – mitochondrial 16S rRNA gene, 10-11 – positive control (*E. coli* DH10B 16S rRNA gene).

Detection limit of the qPCR analysis was 1000 16S rRNA copies/ng of DNA, as it was the number of copies in the most diluted standard used. The standard had C_t_ = 45.385, while negative controls had C_t_ of over 50. Parameters of a standard curve: 62.491 intercept, -5.544 slope, 0.999 adjusted R^2^. The fit was significant, ANOVA F_1,4_ = 5218, p = 2.201e-07. Calculated efficiency was 1.515. Analyses were performed in quadruplicate (technical replicates), for seeds there was one biological replicate (DNA isolated from 10-15 seeds), while for seedlings there were at least three biological replicates (seedlings) per variant.

Samples whose counts are 0 on Fig. SR2 had indeterminable C_t_ values, as the ‘second-derivative’ algorithm of LC480 software was unable to determine their C_t_ value after 55 cycles.

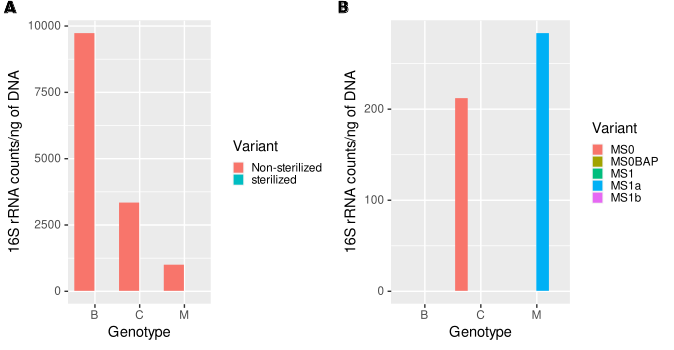

**Fig. SR2.** Surface sterilization of beet seeds causes decrease of the number of non-plastidic, non-mitochondrial 16S rRNA sequences below the detection limit, regardless of genotype (A). Regardless of growth medium variant, seedlings emerging from surface-sterilized seeds have 16S rRNA counts below detection limit (B). Media used for germination were: MS0 – standard Murashige and Skoog, MS0BAP – Murashige and Skoog with BAP, MS1 – Murashige and Skoog with cefotaxime and vancomycin, MS1a – Murashige and Skoog with cefotaxime, vancomycin and tetracyclin, MS1b – Murashige and Skoog with cefotaxime, vancomycin and chloramphenicol.

**Sequencing statistics**

**Table SR2.** Sequencing statistics. Means and SD values were rounded to the nearest integer. Total number of ASVs is not the sum of numbers found in all pools, as there are ones found in more than one pool.

|  | **No. samples** | **Raw reads** | **Final seqs^#^** | **Raw reads mean ± SD** | **Final reads mean ± SD** | **No. ASVs** |
| --- | --- | --- | --- | --- | --- | --- |
| Soils | 782 | 20 387 303 | 8 714 779 | 26 070 ± 10 918 | 11 144 ± 4 883 | 35 992 |
| Roots | 782 | 11 662 884 | 8 210 732 | 14 914 ± 6 242 | 10 500 ± 4 377 | 11 874 |
| Leaves | 778 | 12 318 975 | 10 516 589 | 15 834 ± 4 435 | 13 517 ± 3 939 | 25 893 |
| Inoculant^$^ | 21 | 192 641 | 129 554 | 9 173 ± 2 142 | 6 169 ± 1 605 | 1 224 |
| Total | 2 363 | 44 561 803 | 27 571 654 | 18 858 ± 7 158 | 11 668 ± 4 376 | 70 605 |

\# - the number of denoised, merged and non-chimeric sequences; $ - apart from inoculant, there were mock community, root and leaf samples in this pool

Positive control (staggered mock community HM-283D from BEI Resources) was sequenced along with inoculant samples. 18 amplicons, out of 20 that should be present, were recovered. The missing ones were those of the lowest concentration (*Actinomyces baumanii* and *Streptococcus pneumoniae*), showing that sequencing coverage was low for this sample. No additional ASVs were found in mock community, which suggested that stringency of reads processing was adequate.

Negative controls (libraries generated from C6 (elution) buffer) were sequenced with each pool of samples. As they contained no or only a few reads, likely due to barcodes misassignment, they were excluded from analysis. Consistently, no systematic batch effect was detected, e.g. leaf samples sequenced along with inoculant clustered with other libraries derived from leaves.

In spite of the use of blocking primers numbers of mitochondrial and plastidic sequences was high in libraries derived from plant tissues, particularly in leaves (in the range of 20-90%). Removal of such sequences reduced the number of ASVs to 62 289, and sequences to 10 847 891 (4 618 ± 5 208 per sample). Rarefying to 900 sequences per sample further decreased the number of samples to 1 496. There were 1 322 087 sequences from 26 963 ASVs in the final, rarefied dataset.

**Abundances of a differentially abundant ASV measured by amplicon sequencing and qPCR are significantly correlated**

Primers specific for ASV10 (classified as *Pseudomonas*) were designed using DECIPHER as described in SM7m. Standard curve was constructed using purified amplicon generated on mixed roots and leaves DNA. Detection limit was 1000 copies/ng of DNA, the number of molecules in the most dilute standard. Negative controls had C_t_ over 35. Parameters of the standard curve: 37.18 intercept, -3.49 slope, 0.9999 adjusted R^2^. The fit was highly significant: ANOVA F_1,4_ = 50 040, p = 2.4e-09. Calculated efficiency was 1.93.

ASV10 abundances were extracted from rarefied sequencing data (for roots and leaves separately) and their Spearman’s correlation with qPCR results was highly significant in both cases (leaves: p = 9.8e-06, roots: p = 0). However, correlation was much weaker in leaves, probably due to the ASV10 abundance being below detection limits of qPCR (Fig. SR3A). Significant correlations corroborate the sequencing results.

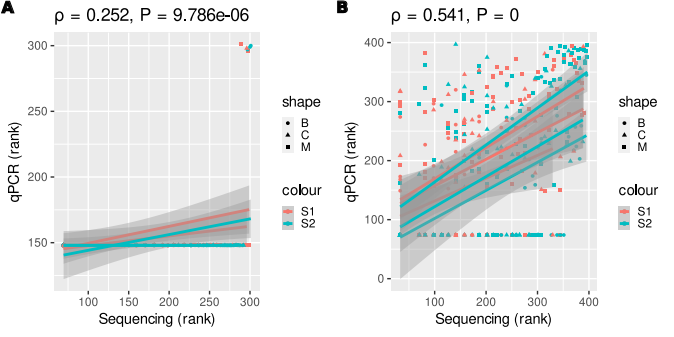

**Fig. SR3**. Spearman’s correlation of ASV10 abundance measured by qPCR and amplicon sequencing. A - leaves B - roots

**There is no significant influence of sequencing run and pot on both alpha and beta-diversity**

To assess influence of random factors (sequencing run and pot, which represent technical variance) we used variance partitioning. As sequencing run was almost identical to material (soil, leaf, root and inoculant libraries were sequenced in different MiSeq runs) estimation of run influence was restricted to leaves and roots samples that were sequenced with inoculant ones (n = 14)

**PICRUSt2 results are of medium quality**

Weighted nearest sequenced taxon index (NSTI), a measure of PICRUSt2 predictions quality, was relatively high for analyzed communities (Fig. SR4), with mean values of 0.115, 0.099, and 0.093 for leaves, roots and soils, respectively. The differences between groups were significant, as assessed by robust ANOVA (overall test for location shift: n = 1493, F_3,1490_ = 80.562, p = 0). Such a high NSTI value is indicative of large proportion of taxa with unsequenced genomes in analyzed communities and suggests that the results need to be interpreted with caution.

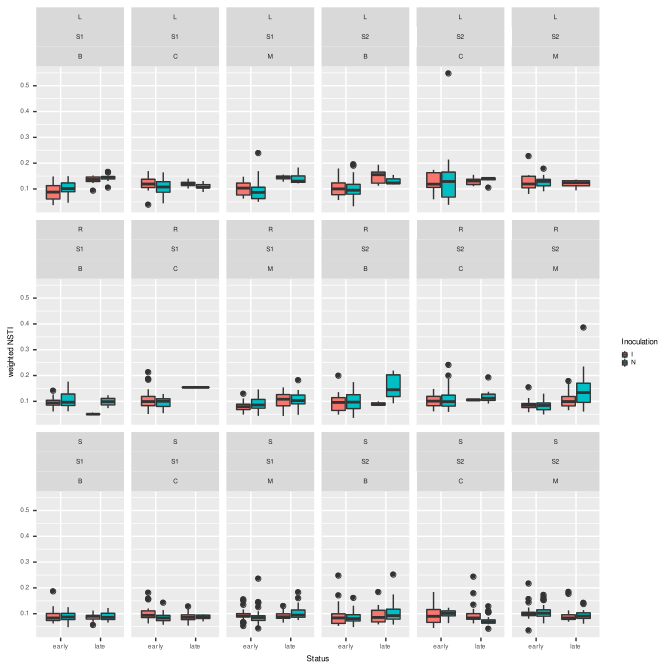
**Fig. SR4**. NSTI (Nearest Sequenced Taxon Indes) values in experimental variants. Values > 0.03 indicate large proportion of unsequenced taxa.

**Soils differ in bacterial community structure, alpha-diversity and functional potential**

Soils used as growth substrate (T0 samples – before inoculation, but after four weeks of axenic plants’ growth) differed in bacterial community structure (Fig. SR5A; permutational test of a dbRDA model: n = 89, F_1_ = 10.87, p = 0.001, betadisper permutational test: F_1_ = 20.00, p = 2.33e-5, explained variance: 10.49%). The same applied to PICRUSt2-predicted functional potential (Fig. SR5B; permutational test of a dbRDA model: F_1_ = 11.21, p = 0.001, betadisper permutational test: F_1_ = 2.59, p = 0.11, explained variance: 10.79%). Diversity (Shannon’s H’), evenness and richness were lower in S2 than in S1 (Fig. SR5C; Wilcoxon test: H’ – W = 1292, p = 0.013, E – W = 1252, p = 0.031, observed number of ASVs – W = 1397.5, p = 8.35e-4). Functional diversity was similar in both soils, but richness was lower, and evenness higher in S2 (Fig. SR5D; Wilcoxon test: H’ – W = 760, p = 0.059, E – W = 572 p = 5.00e-4, observed number of functions – W = 1454.5, p = 1.40e-4). Libraries generated from S1 samples were more abundant in sequences affiliated with *Bacillus* and rare taxa, while *Brevundimonas*, *Delftia* and *Pseudoxanthomonas* were more frequent in S2 (Fig. SR5E).

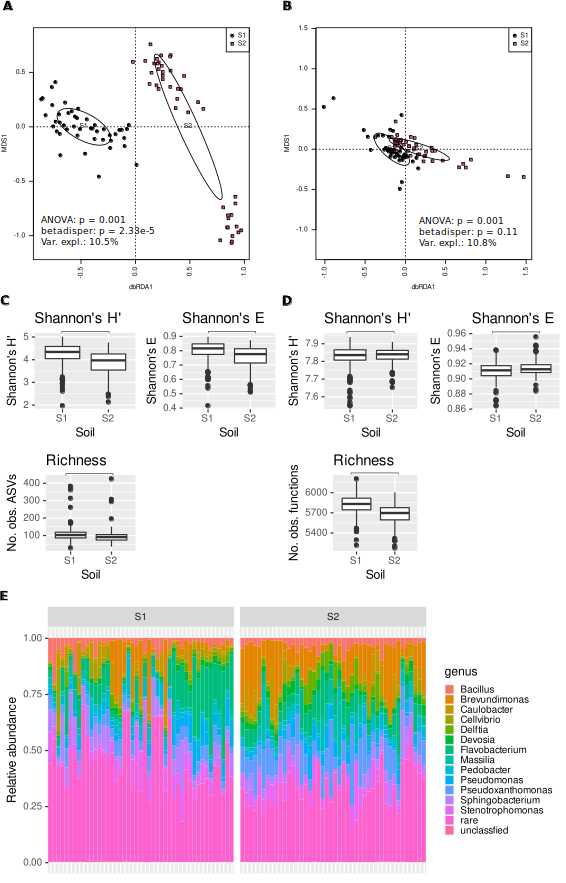

**Fig. SR5.** Microbiomes of S1 and S2 soils differ in terms of structure (A), functional potential (B), alpha-diversity of ASVs (C), functional diversity (D), and taxonomic compostion (E).

**Communities in leaves, roots and soils differ**

**Table SR3.** ASVs characteristic for leaves, soils and roots. Rows containing ASVs characteristic for plant tissues were written in boldface font, while those containing ASVs characteristic for soils were underlined.

| **ASV** | **Comparison^1^** | **Fold change^2^** | **DESeq2 q^3^** | **ALDEx2 q^4^** | **Taxonomy** | **Material^5^** |
| --- | --- | --- | --- | --- | --- | --- |
| ASV13 | LR | 2.12 | 2.55e-91 | 5.17e-69 | Sphingobacterium | L |
| **ASV23** | **LR** | **1.22** | **7.63e-24** | **9.56e-176** | **Chryseobacterium** | **P** |
| ASV57 | LR | 1.16 | 1.15e-23 | 7.35e-24 | Azospirillum | L |
| ASV22 | LR | -1.11 | 7.15-28 | 0.00018 | Comamonas | R |
| ASV76 | LR | -1.12 | 7.69e-28 | 0.00016 | Chitinophagaceae | R |
| ASV16 | LR | -1.49 | 1.46e-38 | 7.52e-18 | Stenotrophomonas | R |
| ASV18 | LR | -2.10 | 2.31e-76 | 0.0015 | Pseudoxanthomonas |  |
| ASV20 | LR | -2.59 | 9.95e-142 | 7.72e-33 | Fluviicola |  |
| ASV7 | LR | -3.00 | 7.53e-149 | 9.13e-28 | Cellvibrio | R |
| **ASV23** | **LS** | **6.24** | **0** | **9.56e-176** | **Chryseobacterium** | **P** |
| **ASV80** | **LS** | **2.06** | **7.32e-95** | **2.39e-09** | **Achromobacter** | **P** |
| ASV57 | LS | 2.05 | 1.39e-72 | 7.35e-24 | Azospirillum | L |
| **ASV15** | **LS** | **1.77** | **7.02e-43** | **2.74e-106** | **Flavobacterium** | **P** |
| **ASV4** | **LS** | **1.59** | **1.09e-50** | **3.17e-45** | **Flavobacterium** | **P** |
| ASV11 | LS | 1.55 | 5.72e-51 | 1.33e-08 | Caulobacter |  |
| ASV522 | LS | 1.33 | 7.46e-50 | 4.43e-16 | Bacillus |  |
| ASV7 | LS | -1.07 | 2.31e-20 | 9.13e-28 | Cellvibrio |  |
| ASV10 | LS | -1.09 | 1.02e-20 | 2.54e-15 | Pseudomonas |  |
| ASV29 | LS | -1.17 | 1.36e-25 | 1.46e-05 | Pseudomonas |  |
| ASV6 | LS | -1.29 | 1.41e-39 | 9.75e-10 | Brevundimonas | S |
| ASV44 | LS | -1.62 | 1.71e-92 | 9.77e-54 | Reyranella | S |
| ASV18 | LS | -1.74 | 7.24e-54 | 0.0015 | Pseudoxanthomonas |  |
| ASV14 | LS | -1.83 | 7.36e-96 | 6.01e-52 | Devosia | S |
| ASV9 | LS | -1.94 | 3.27e-77 | 0.00060 | Stenotrophomonas | S |
| ASV20 | LS | -2.04 | 4.56e-90 | 7.72e-33 | Fluviicola |  |
| ASV33 | LS | -2.43 | 3.15e-137 | 1.39e-34 | Micropepsaceae |  |
| ASV24 | LS | -2.50 | 4.62e-208 | 9.38e-98 | Pseudohongiellaceae - BIyi10 | S |
| ASV27 | LS | -2.69 | 5.73e-190 | 1.50e-102 | Pedobacter | S |
| ASV3 | LS | -3.07 | 6.75e-201 | 1.76e-32 | Pseudoxanthomonas | S |
| ASV21 | LS | -3.60 | 2.86e-225 | 3.04e-36 | Sphingopyxis | S |
| ASV27 | SR | 3.60 | 0 | 1.50e-102 | Pedobacter | S |
| ASV21 | SR | 3.30 | 3.58e-198 | 3.04e-36 | Sphingopyxis | S |
| ASV3 | SR | 2.86 | 2.20e-176 | 6.75e-201 | Pseudoxanthomonas | S |
| ASV33 | SR | 2.77 | 3.79e-178 | 1.39e-34 | Micropepsaceae | S |
| ASV13 | SR | 2.73 | 5.48e-154 | 5.17e-69 | Sphingobacterium | S,L |
| ASV24 | SR | 2.65 | 1.80e-241 | 9.38e-98 | BIyi10 | S |
| ASV44 | SR | 1.76 | 4.42e-112 | 9.77e-54 | Reyranella | S |
| ASV19 | SR | 1.76 | 2.45e-60 | 0.0039 | Caulobacter |  |
| ASV9 | SR | 1.59 | 8.83e-54 | 0.00060 | Stenotropho  monas | S |
| ASV6 | SR | 1.25 | 2.26e-37 | 9.75e-10 | Brevundimonas | S |
| ASV14 | SR | 1.03 | 1.05e-32 | 6.01e-52 | Devosia | S |
| **ASV4** | **SR** | **-1.48** | **1.76e-44** | **3.17e-45** | **Flavobacterium** | **P** |
| **ASV15** | **SR** | **-1.66** | **1.44e-37** | **2.74e-106** | **Flavobacterium** | **P** |
| **ASV80** | **SR** | **-1.90** | **4.46e-81** | **2.39e-09** | **Achromobacter** | **P** |
| ASV7 | SR | -1.94 | 1.63e-66 | 9.13e-28 | Cellvibrio | R |
| ASV16 | SR | -1.98 | 2.14e-69 | 7.52e-18 | Stenotrophomonas | R |
| **ASV23** | **SR** | **-5.03** | **0** | **9.56e-176** | **Chryseobacterium** | **P** |

1 - LR – leaves vs. roots (positive FC – higher abundance in leaves), LS – leaves vs. soils, SR – soils vs. roots

2 – log_2_ fold change (FC) reported by DESeq2

3 – Benjamini-Hochberg corrected p-value of Wald test reported by DESeq2

4 – Benjamini-Hochberg corrected p-value of Kruskal-Wallis test reported by ALDEx2

5 – Material for which a given ASV is characteristic – S – soil, R – roots, L – leaves, P – plant (roots + leaves)

**Table SR4.** PICRUSt2-predicted functions characteristic for soils, roots and leaves. Ten KO functions with largest absolute values of log_2_ fold change per comparison are shown.

| **Function** | **Comp.** | **Fold change** | **DESeq2 q** | **ALDEx2 q** | **Description** | **Pathway** | **Process** |
| --- | --- | --- | --- | --- | --- | --- | --- |
| K16638 | LR | 2,44 | 0 |  | K16638, exoU; exoenzyme U | - | Plant pathogenicity |
| K15502 | LR | 2,44 | 0 |  | ANKRD28; serine/threonine-protein phosphatase 6 regulatory ankyrin repeat subunit A | - | DNA repair and recombination |
| K10921 | LR | 2,29 | 0 |  | toxR; cholera toxin transcriptional activator | - | Biofilm formation |
| K06931 | LR | 2,2 | 0 |  | K06931; uncharacterized protein | - | - |
| K08651 | LR | 2,19 | 0 |  | E3.4.21.66; thermitase [EC:3.4.21.66] | - | Protein folding |
| K19278 | LR | -2,66 | 0 |  | aac6-Ib; aminoglycoside 6'-N-acetyltransferase Ib [EC:2.3.1.82] | - | Inter-organismal interactions (antibiotic resistance) |
| K12997 | LR | -2,71 | 0 |  | rgpB; rhamnosyltransferase [EC:2.4.1.-] | - | Lipopolysaccharide biosynthesis, inter-organismal interactions |
| K19301 | LR | -2,72 | 0 |  | aac6-II; aminoglycoside 6'-N-acetyltransferase II [EC:2.3.1.82] | - | Inter-organismal interactions |
| K13243 | LR | -2,88 | 0 |  | dos; c-di-GMP-specific phosphodiesterase [EC:3.1.4.52] | - | Regulation |
| K12516 | LR | -3,04 | 0 |  | bigA; putative surface-exposed virulence protein | - | Pathogenicity |
| K19694 | LS | 3,86 | 0 |  | chiS; two-component system, sensor histidine kinase ChiS | - | Regulation of natural competence |
| K18642 | LS | 3,23 | 0 |  | creS; crescentin |  | Regulation of cell shape |
| K07257 | LS | 3,09 | 0 |  | spsF; spore coat polysaccharide biosynthesis protein SpsF |  | Spore formation |
| K15894 | LS | 2,72 | 0 |  | pseB; UDP-N-acetylglucosamine 4,6-dehydratase [EC:4.2.1.115] | Amino sugar and nucleotide sugar metabolism | Inter-organismal interactions (cell wall antigen biosynthesis) |
| K06931 | LS | 2,69 | 0 |  | K06931; uncharacterized protein | - | - |
| K05709 | LS | -1,78 | 0 |  | hcaF, hcaA2; 3-phenylpropionate/trans-cinnamate dioxygenase subunit beta [EC:1.14.12.19] | Phenylalanine metabolism | Degradation of aromatic compounds |
| K01822 | LS | -1,78 | 0 |  | E5.3.3.1; steroid Delta-isomerase [EC:5.3.3.1] | Steroid degradation | Xenobiotics biodegradation and metabolism |
| K02638 | LS | -1,83 | 0 |  | petE; plastocyanin | Photosynthesis | Photosynthesis |
| K07505 | LS | -1,97 | 0 |  | repA; regulatory protein RepA | - | DNA replication |
| K14439 | LS | -2,14 | 0 |  | SMARCAD1; SWI/SNF-related matrix-associated actin-dependent regulator of chromatin subfamily A containing DEAD/H box 1 [EC:3.6.4.12] | Artefact – eukaryotic function, should not be present in PICRUSt2 results |  |
| K07505 | SR | 3,12 | 0 |  | repA; regulatory protein RepA | - | DNA replication |
| K19170 | SR | 2,81 | 0 |  | dndC; DNA sulfur modification protein DndC | - | Prokaryotic defense against phages |
| K19171 | SR | 2,69 | 0 |  | dndD; DNA sulfur modification protein DndD | - | Prokaryotic defense against phages |
| K14439 | SR | 2,69 | 0 |  | SMARCAD1; SWI/SNF-related matrix-associated actin-dependent regulator of chromatin subfamily A containing DEAD/H box 1 [EC:3.6.4.12] | Artefact – eukaryotic function, should not be present in PICRUSt2 results |  |
| K03395 | SR | 2,59 | 0 |  | aac3-I; aminoglycoside 3-N-acetyltransferase I [EC:2.3.1.60] | - | Inter-organismal interactions (antibiotic resistance) |
| K13552 | SR | -2,32 | 0 |  | btrQ; paromamine 6'-oxidase [EC:1.1.3.43] | Neomycin, kanamycin and gentamycin biosynthesis | Inter-organismal interactions (antibiotic biosynthesis) |
| K18642 | SR | -2,35 | 0 |  | creS; crescentin |  | Regulation of cell shape |
| K12516 | SR | -2,36 | 0 |  | bigA; putative surface-exposed virulence protein | - | Pathogenicity |
| K18336 | SR | -2,46 | 0 | 3,70060888642378E-25 | K18336; 2,4-diketo-3-deoxy-L-fuconate hydrolase [EC:3.7.1.-] | Fructose and mannose metabolism | Carbohydrate metabolism |
| K19694 | SR | -2,73 | 0 | 2,10751808638163E-84 | chiS; two-component system, sensor histidine kinase ChiS | - | Regulation of natural competence |

**Table SR5.** Variables’ influence on bacterial communities, assessed based on d05 generalized UniFrac distance matrix. Means of variance explained for two models (one involving time, and the other status) are given. Time and status were not tested together.

| **Variable** | **Total var. expl.^1^** | **Unique var. expl.^2^** | **R sq.** | **p-value^3^** |
| --- | --- | --- | --- | --- |
| Material | 12.1 | 9.6 | 92.82 | 0.001 |
| Time | 8.2 | 5.8 | 28.37 | 0.001 |
| Status | 6.9 | 4.7 | 90.27 | 0.001 |
| Soil | 2.9 | 2.7 | 54.75 | 0.001 |
| Genotype | 1.8 | 1.7 | 17.72 | 0.001 |
| Inoculation | 0.28 | 0.15 | 3.54 | 0.001 |

1 – total variance explained, 2 – variance explained solely by a variable in question (i.e. with other variables conditioned out), 3 – p-value of a permutational test of unique variation fraction significance in a dbRDA model

**Samples cluster into ‘late’ and ‘early’ groups regardless of material, soil and plant genotype**

Grouping according to status was significant in each experimental variant apart from roots of genotype C grown in S1, where there were only one late sample and, consequently, there was no possibility of conducting tests. However, tests for homogeneity of multidimensional dispersion showed significant differences of variance between early and late groups, which suggested possibility of the permutational tests of dbRDA models yielding significant p-value not because of difference in location but in spread. Nevertheless, it seems unlikely, as separation of clusters is obvious in each ordination plot generated.

Interestingly, percent of variance explained by status was drastically lower in roots than in soils and leaves and consistently lower in S2 than in S1.

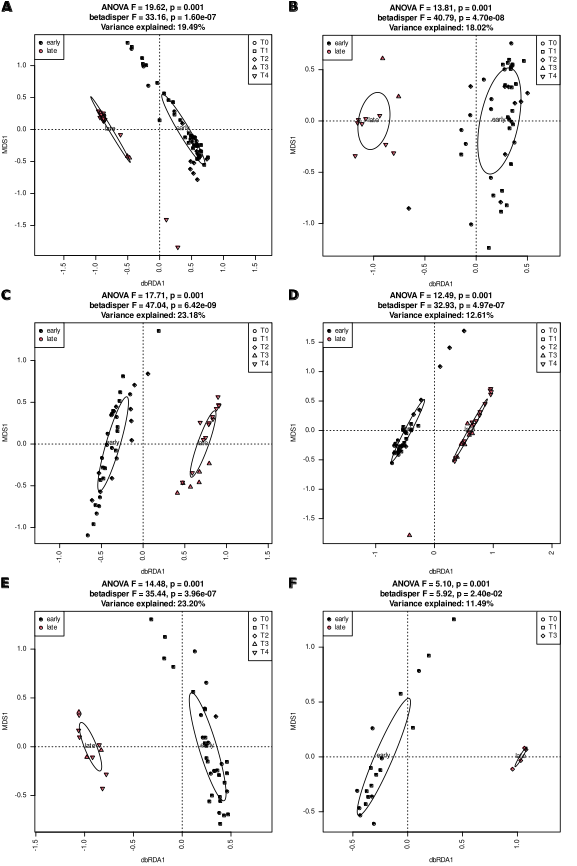

**Fig. SR6.** Leaf samples form ‘early’ and ‘late’ clusters regardless of soil and plant genotype. dbRDA analysis based on d05 generalized UniFrac distance matrix. AB – genotype B, CD - genotype C, EF – genotype M; ACE – soil S1, BDF – soil S2.

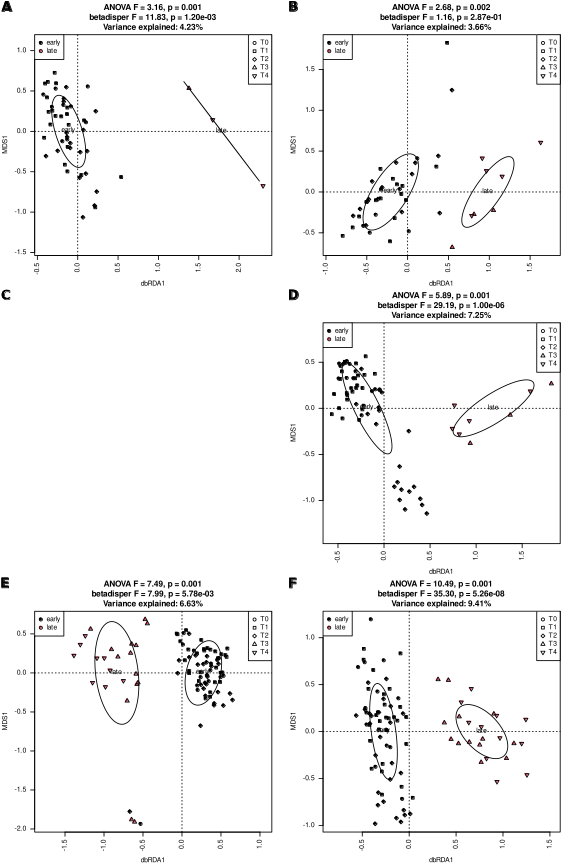

**Fig. SR7.** Root samples form ‘early’ and ‘late’ clusters regardless of soil and plant genotype. dbRDA analysis based on d05 generalized UniFrac distance matrix. AB – genotype B, CD - genotype C, EF – genotype M; ACE – soil S1, BDF – soil S2.

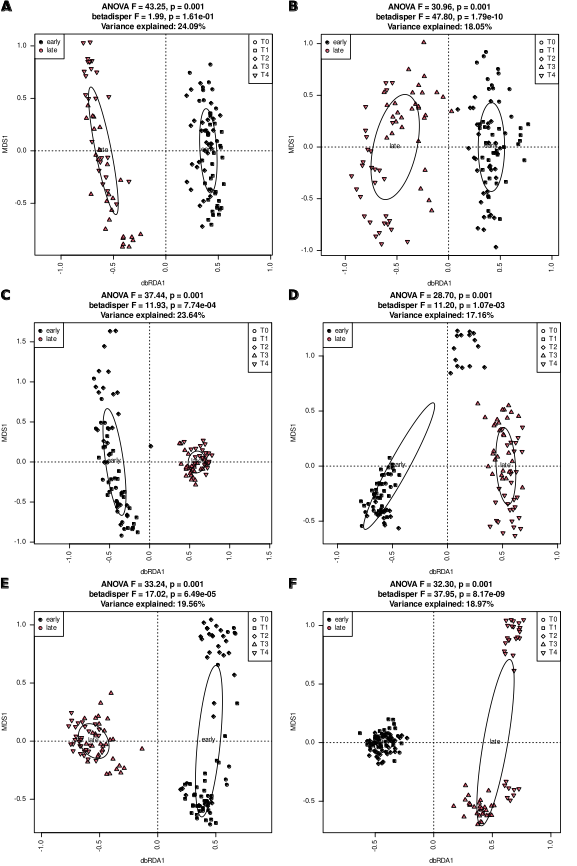

**Fig. SR8.** Soil samples form ‘early’ and ‘late’ clusters regardless of soil and plant genotype. dbRDA analysis based on d05 generalized UniFrac distance matrix. AB – genotype B, CD - genotype C, EF – genotype M; ACE – soil S1, BDF – soil S2.

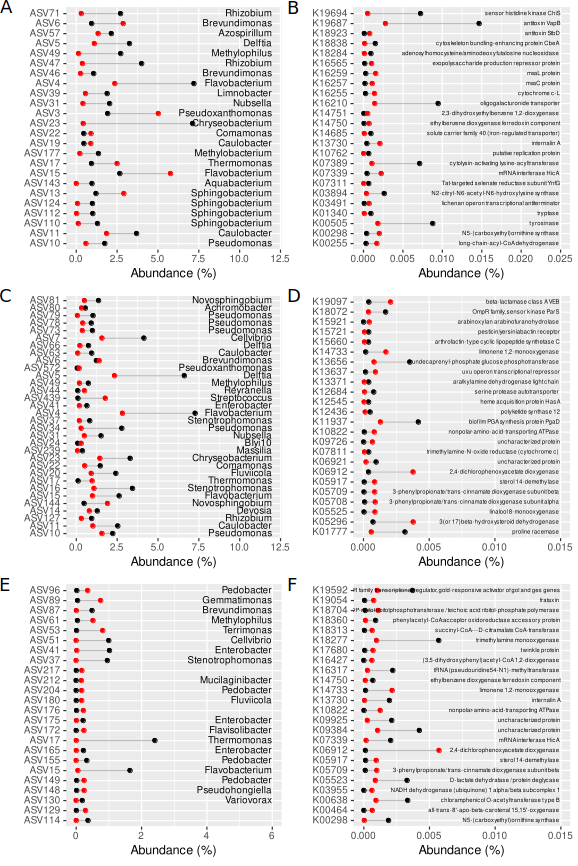

**Fig. SR9.** Organisms (ACE) and functions (BDF) characteristic for early and late samples are different in leaves (AB), roots (CD) and soils (EF). Black dots denote abundance in early samples, red dots abundance in late ones. Twenty-four features of greatest fold-change are shown on each panel

**Nestedness decreases over time and is significantly higher in early samples than in late ones**

**Table SR6.** Significance testing of differences in mean weighted NODF values in early and late samples in different experimental variants. Wilcoxon test was used. Significant p-value is given in boldface font.

| Variant | Test statistics (W) | p-value |
| --- | --- | --- |
| S1 B | 218 | **6.85e-4** |
| S1 C | 86 | 7.65e-2 |
| S1 M | 293 | **5.51e-4** |
| S2 B | 152 | **2.12e-3** |
| S2 C | 410 | **1.57e-10** |
| S2 M | 74 | **2.98e-2** |

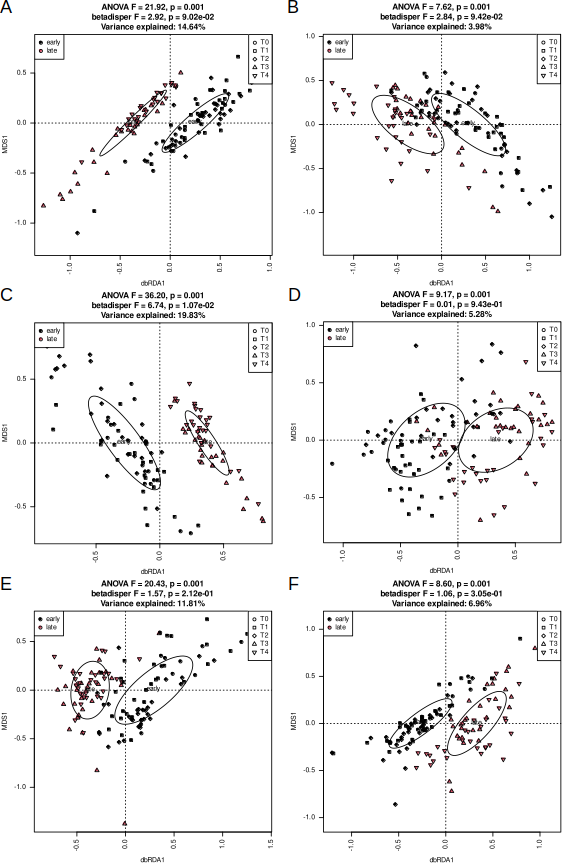

**Fig. SR10**. Functional potential of bacterial communities living in early and late soil samples differ, but variance does not. Results of dbRDA analysis of Morisita-Horn distance matrices derived from PICRUSt2-predicted KO abundance tables are shown for soil S1 (ACE) and S2 (BDF) and genotype B (AB), C (CD) and M (EF). dbRDA and betadisper models were tested with permutational tests.

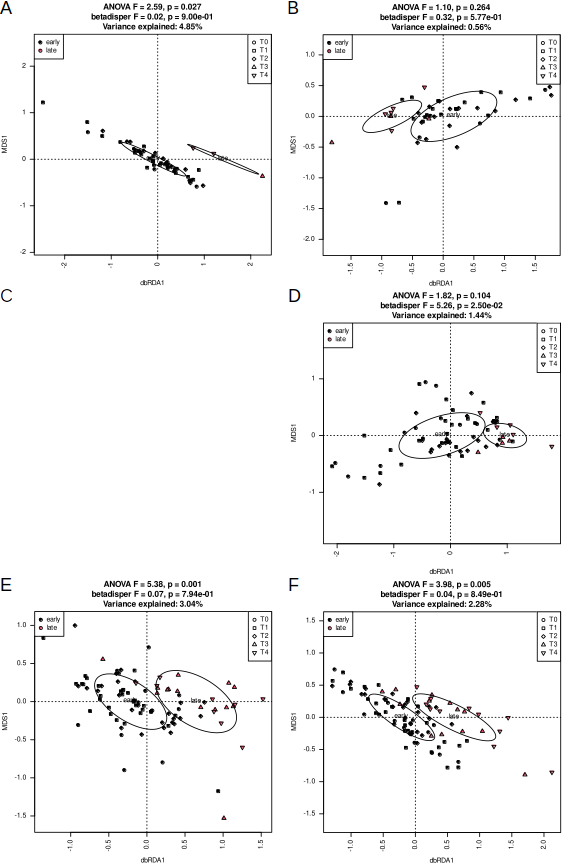

**Fig. SR11.** Functional potential of bacterial communities living in early and late root samples differ, but variance does not. Results of dbRDA analysis of Morisita-Horn distance matrices derived from PICRUSt2-predicted KO abundance tables are shown for soil S1 (ACE) and S2 (BDF) and genotype B (AB), C (CD) and M (EF). dbRDA and betadisper models were tested with permutational tests.

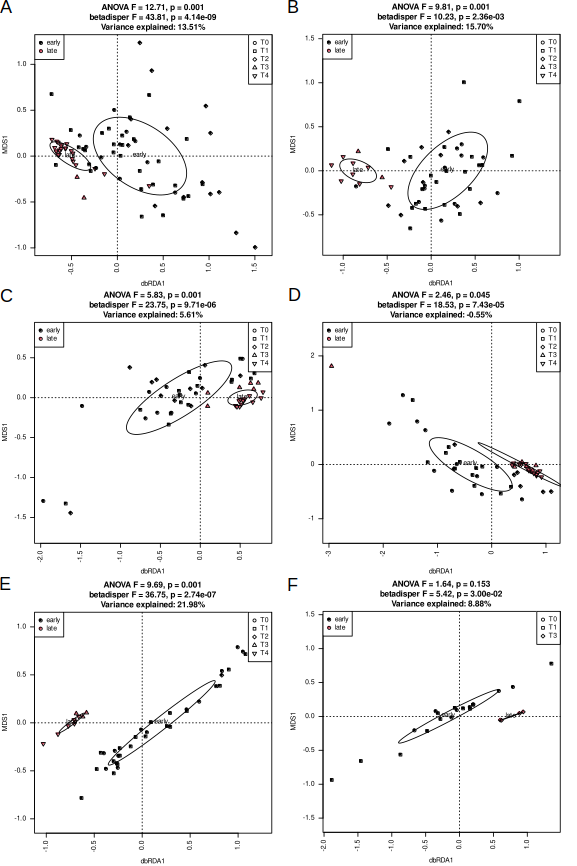

**Fig. SR12.** Functional potential of bacterial communities living in early and late leaf samples differ, and variance does not. Results of dbRDA analysis of Morisita-Horn distance matrices derived from PICRUSt2-predicted KO abundance tables are shown for soil S1 (ACE) and S2 (BDF) and genotype B (AB), C (CD) and M (EF). dbRDA and betadisper models were tested with permutational tests.

**Genotype influenced bacterial communities in soils and beet plants regardless of soil type and status**

**Genotype influences both soil and endophytic bacterial communities regardless of status**

Bacterial communities and sets of imputed KO functions clustered according to genotype regardless of material, soil as well as status (Figs. SR13, SR16, SR19 and SR22, SR25, SR28 for ASVs and KO functions, respectively). Genotype influence was strongest (i.e. percent of variance explained was highest) in soils and stronger at the level of ASVs than of KO functions.

Alpha-diversity varied depending on material, soil and status at the level of ASVs. Bacterial load differed only in early root samples, where it was highest in genotype M (Figs. SR14, SR17 and SR20). In general, in soil S1 diversity was highest in genotype B samples, regardless of material, in soil S2, the same was true for genotype M. The functional diversity pattern was essentially the same (Figs. SR23, SR26 and SR29). In all materials and regardless of status ASVs classified as *Gammaproteobacteria* were characteristic for sea beet (genotype M), while *Alphaproteobacteria* and *Bacteroidia* were typical for cultivated beets (genotypes B and C) (Figs. SR15, SR18 and SR21). Sets of KO functions characteristic for particular genotypes also depended on soil and status, and the number of differentially represented functions was highest in soils and lowest in leaves. Moreover, endophytic bacterial communities were functionally more diverse in early samples than in late ones regardless of genotype.

Regardless of soil and status the degree of nestedness was higher in sea beet than in sugar beet varieties, however in late samples grown in soil S1 this difference was not significant (Fig. SR31). Genotypes did not differ clearly in shares of community assembly mechanisms, regardless of the stage (Fig. 10).

Bacterial community structure differed in the three analyzed genotypes, regardless of material, soil and status (Figs. SR13 (soils), SR16 (roots) and SR19 (leaves)). Genotype influence was greater in soils than *in planta* and in early samples than in late ones. The influence at the level of functional potential was smaller than at ASVs level, but the general pattern was the same (Figs. SR22, SR25 and SR28). Alpha-diversity was

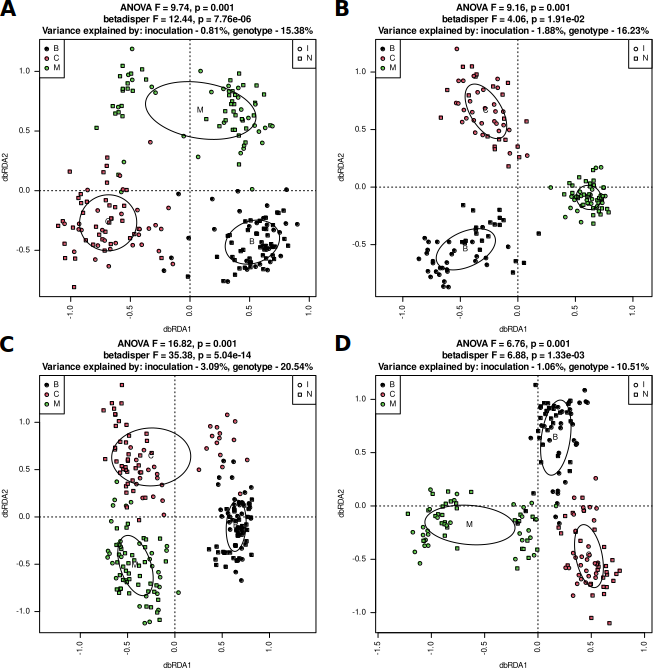
**Fig. SR13**. Soil samples group according to genotype regardless of soil and status. Early samples (AC), late (BD), soil S1 (AB), S2 (CD). dbRDA models based on d05 generalized UniFrac distance matrices were tested with permutational test (999 permutations). Homogeneity of multi-dimensional variance was tested with betadisper test, and percentage of explained variance was assessed with varpart.

**Fig. SR14.** Alpha-diversity measures are higher in samples of soil S2 in which genotype M was grown than in soils in which genotypes B and C were grown, regardless of status. AB – soil S1, CD – soil S2, AC – early samples, BD – late samples. Means ± SD of Shannon’s diversity (H’), Shannon’s evennes (E) and observed number of ASVs are shown.
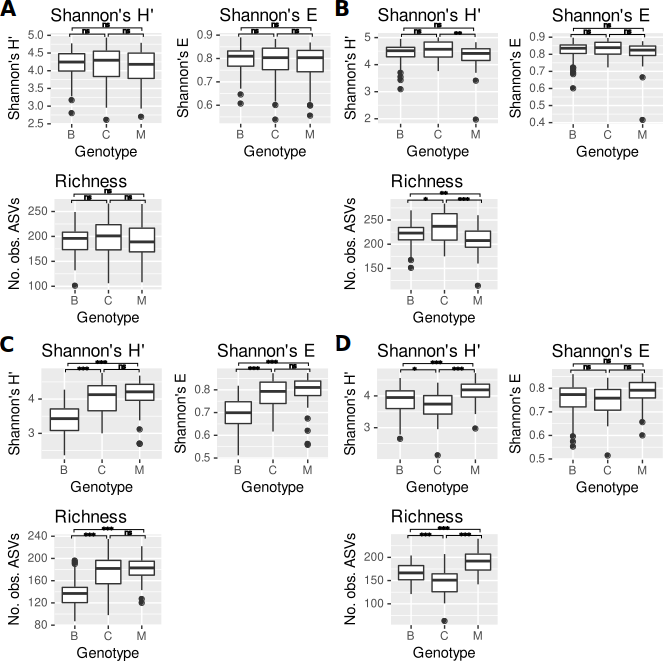
Significance of difference between genotypes (Kruskal-Wallis test with Dunn’s post-hoc procedure using Benjamini-Hochberg FDR) is shown over the plots: *** - p < 0.001, ** - p < 0.01, * - p < 0.05, ns – not significant.

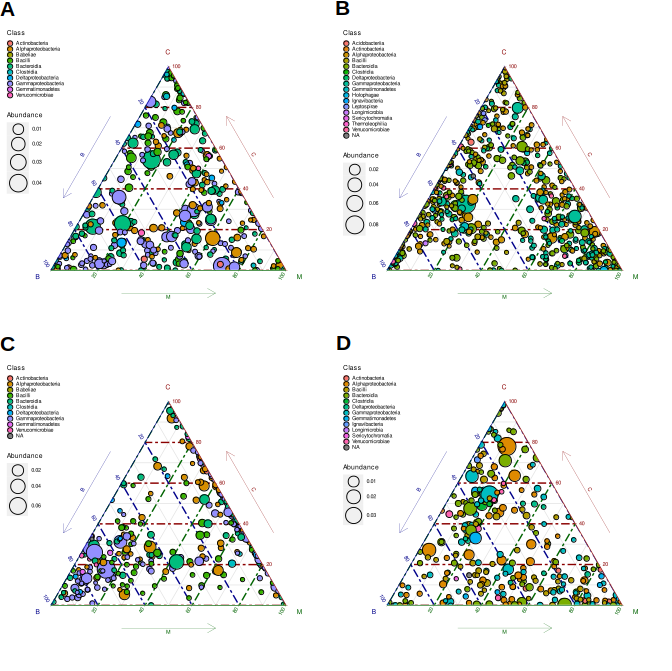
**Fig. SR15.** Sets of organisms characteristic for soil in which different genotypes were grown are different depending on soil and status. Ternary plots of differentially abundant ASVs identified with DESeq2 are shown. AB – soil S1, CD – soil S2, AC – early samples, BD – late samples. Note differeces between panels’ legends.

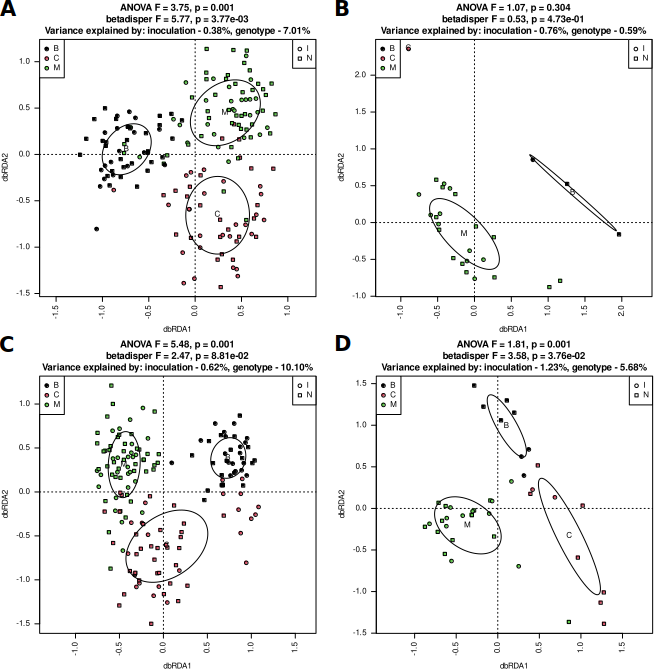
**Fig. SR16**. Root samples group according to genotype regardless of soil and status. Early samples (AB), late (CD), soil S1 (AC), S2 (BD). dbRDA models based on d05 generalized UniFrac distance matrices were tested with permutational test (999 permutations). Homogeneity of multi-dimensional variance was tested with betadisper test, and percentage of explained variance was assessed with varpart.

**Fig. SR17.** Alpha-diversity measures and bacterial load are highest in root samples of genotype M grown in S2 soil , regardless of status, while the same is true for early root samples of genotype C grown in S1. AB – soil S1, CD – soil S2, AC – early samples, BD – late samples. Means ± SD of Shannon’s diversity (H’), Shannon’s evennes (E) and observed number of ASVs are shown.
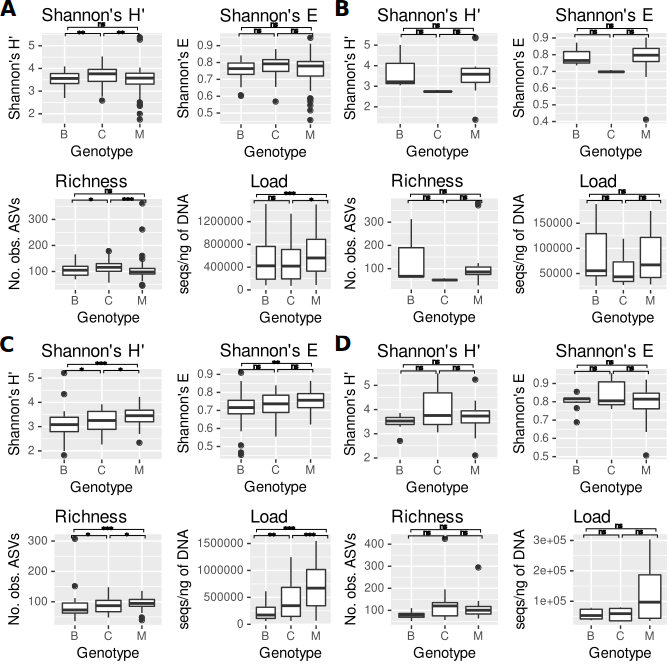
Significance of difference between genotypes (Kruskal-Wallis test with Dunn’s post-hoc procedure using Benjamini-Hochberg FDR) is shown over the plots: *** - p < 0.001, ** - p < 0.01, * - p < 0.05, ns – not significant.

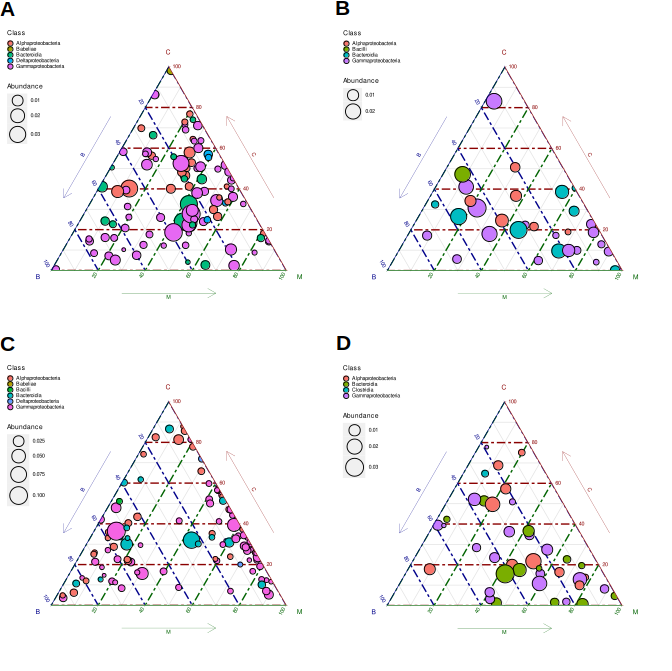
**Fig. SR18.** Sets of organisms characteristic for roots of different genotypes are different depending on soil and status. Ternary plots of differentially abundant ASVs identified with DESeq2 are shown. AB – soil S1, CD – soil S2, AC – early samples, BD – late samples. Note differeces between panels’ legends.

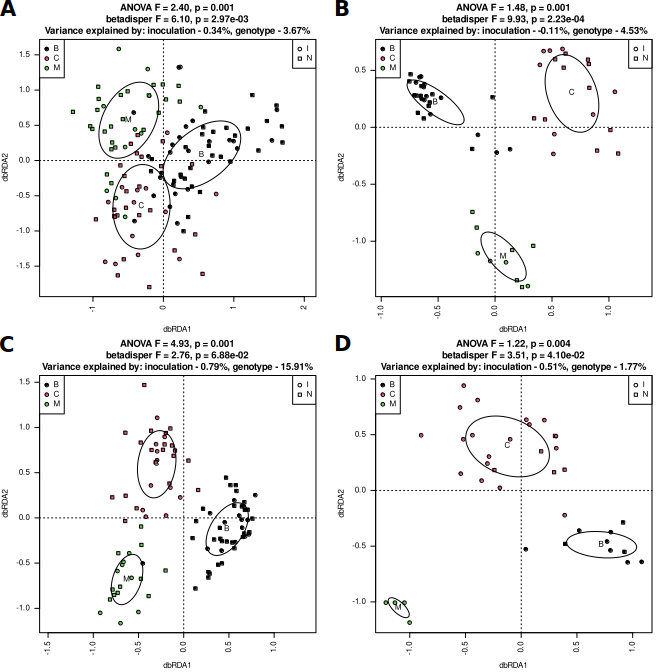
**Fig. SR19**. Leaf samples group according to genotype regardless of soil and status. Early samples (AB), late (CD), soil S1 (AC), S2 (BD). dbRDA models based on d05 generalized UniFrac distance matrices were tested with permutational test (999 permutations). Homogeneity of multi-dimensional variance was tested with betadisper test, and percentage of explained variance was assessed with varpart.

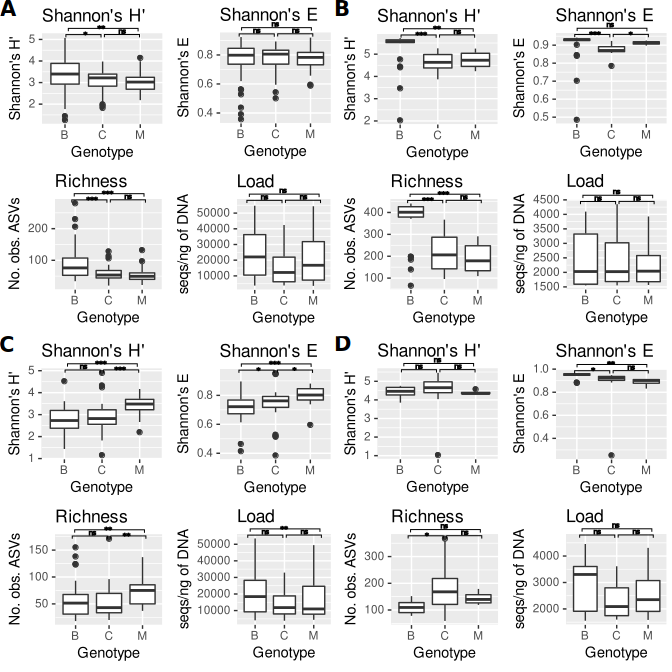
**Fig. SR20.** Alpha-diversity measures but not bacterial load are higher in early leaf samples of genotype M grown in S2 soil while the same is true for genotype B in S1 soil. AB – soil S1, CD – soil S2, AC – early samples, BD – late samples. Means ± SD of Shannon’s diversity (H’), Shannon’s evennes (E) and observed number of ASVs are shown. Significance of difference between genotypes (Kruskal-Wallis test with Dunn’s post-hoc procedure using Benjamini-Hochberg FDR) is shown over the plots: *** - p < 0.001, ** - p < 0.01, * - p < 0.05, ns – not significant.

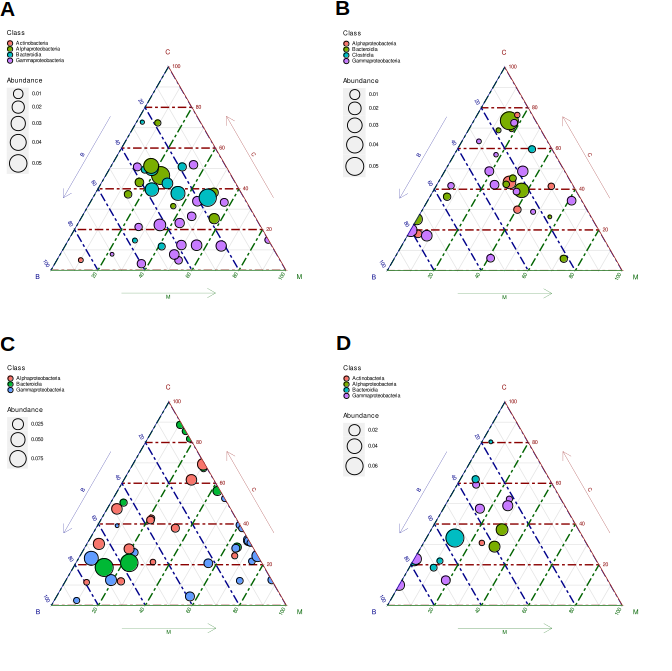
**Fig. SR21.** Sets of organisms characteristic for leaves of different genotypes are different depending on soil and status. Ternary plots of differentially abundant ASVs identified with DESeq2 are shown. AB – soil S1, CD – soil S2, AC – early samples, BD – late samples. Note differeces between panels’ legends.

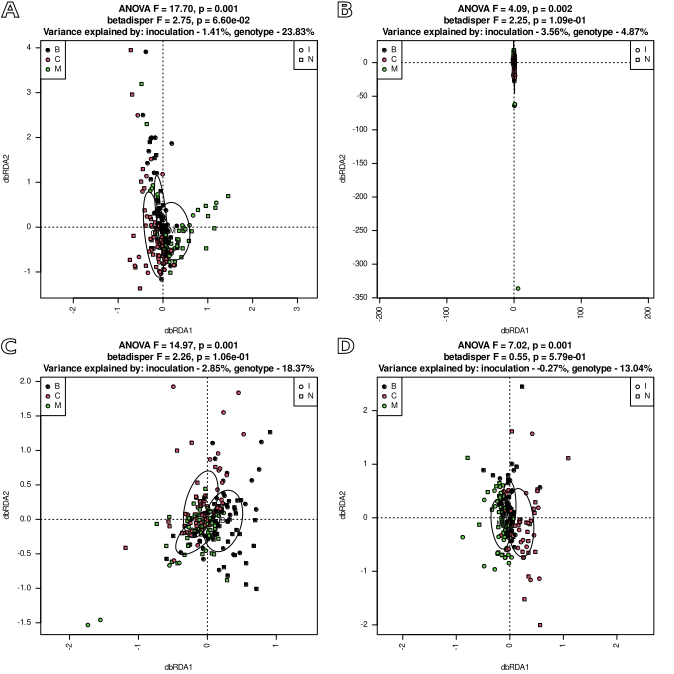

**Fig. SR22**. Sets of KO functions predicted to be encoded by organisms living in soil samples group according to genotype regardless of soil and status. Early samples (AB), late (CD), soil S1 (AC), S2 (BD). dbRDA models based on Morisita-Horn distance matrices were tested with permutational test (999 permutations). Homogeneity of multi-dimensional variance was tested with betadisper test, and percentage of explained variance was assessed with varpart.

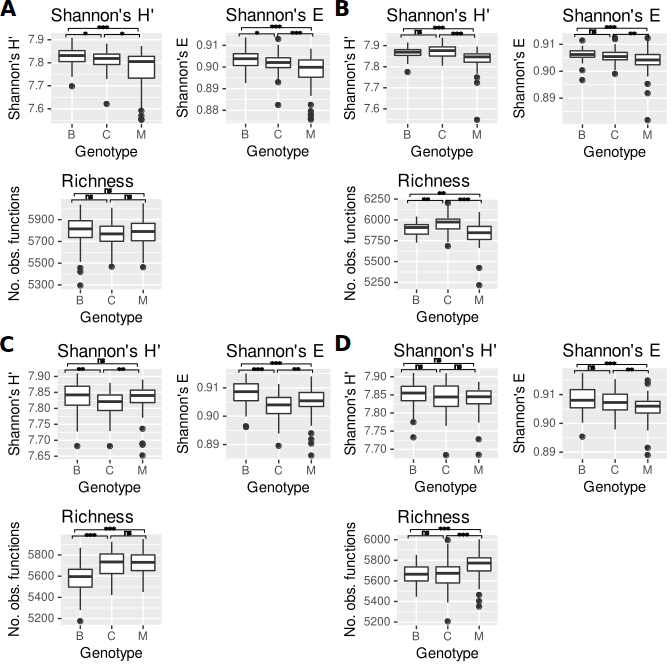
**Fig. SR23.** Functional diversity of soil samples’ bacterial communities depends genotype. AB – soil S1, CD – soil S2, AC – early samples, BD – late samples. Means ± SD of Shannon’s diversity (H’), Shannon’s evennes (E) and observed number of KO functions predicted by PICRUSt2 to be encoded by genomes of organisms thriving in the samples are shown. Significance of difference between genotypes (Kruskal-Wallis test with Dunn’s post-hoc procedure using Benjamini-Hochberg FDR) is shown over the plots: *** - p < 0.001, ** - p < 0.01, * - p < 0.05, ns – not significant.

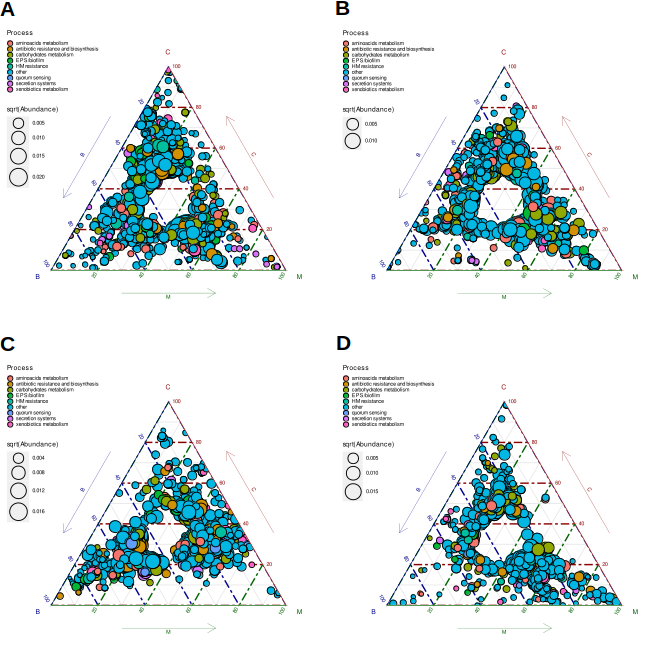
**Fig. SR24.** Sets of KO functions characteristic for soils in which different genotypes grew were different depending on soil and status. Ternary plots of differentially abundant PICRUSt2-predicted KOs identified with DESeq2 are shown. AB – soil S1, CD – soil S2, AC – early samples, BD – late samples. Note differeces between panels’ legends.

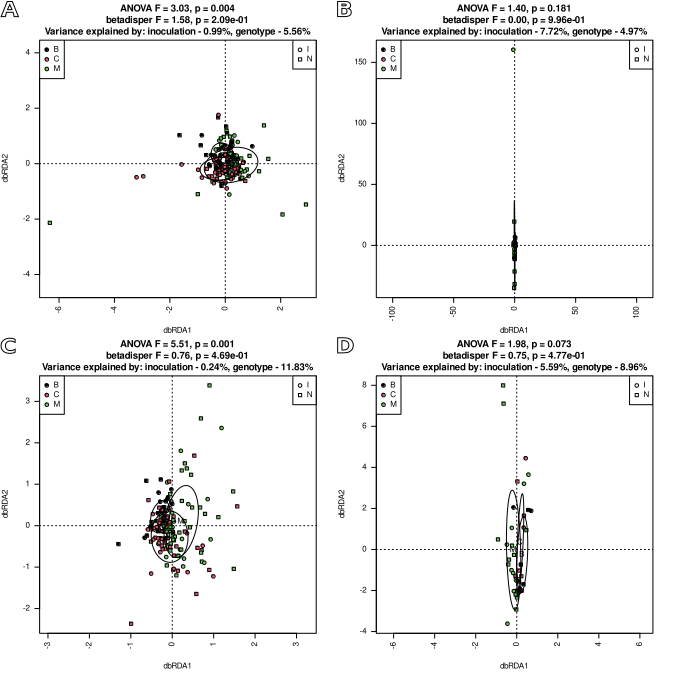
**Fig. SR25**. Sets of KO functions predicted to be encoded by organisms living in root samples group according to genotype regardless of soil and status. Early samples (AB), late (CD), soil S1 (AC), S2 (BD). dbRDA models based on Morisita-Horn distance matrices were tested with permutational test (999 permutations). Homogeneity of multi-dimensional variance was tested with betadisper test, and percentage of explained variance was assessed with varpart.

**Fig. SR26.**
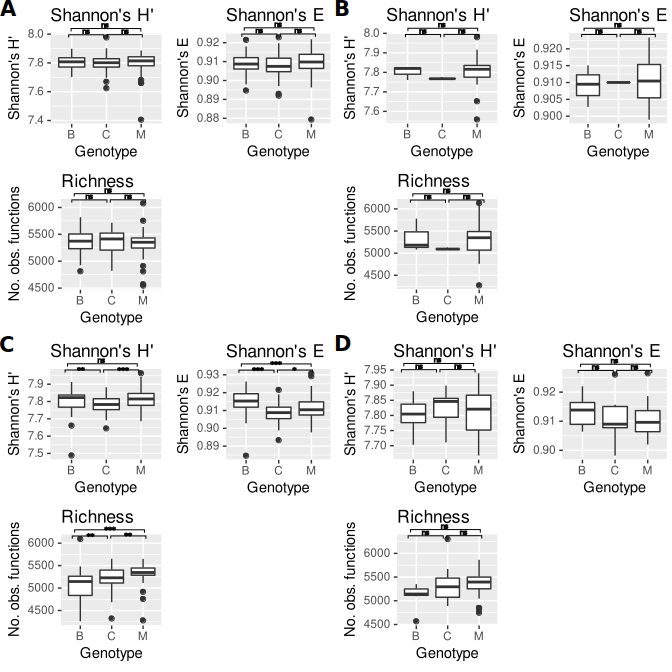
Functional diversity of root samples’ bacterial communities depends genotype. AB – soil S1, CD – soil S2, AC – early samples, BD – late samples. Means ± SD of Shannon’s diversity (H’), Shannon’s evennes (E) and observed number of KO functions predicted by PICRUSt2 to be encoded by genomes of organisms thriving in the samples are shown. Significance of difference between genotypes (Kruskal-Wallis test with Dunn’s post-hoc procedure using Benjamini-Hochberg FDR) is shown over the plots: *** - p < 0.001, ** - p < 0.01, * - p < 0.05, ns – not significant.

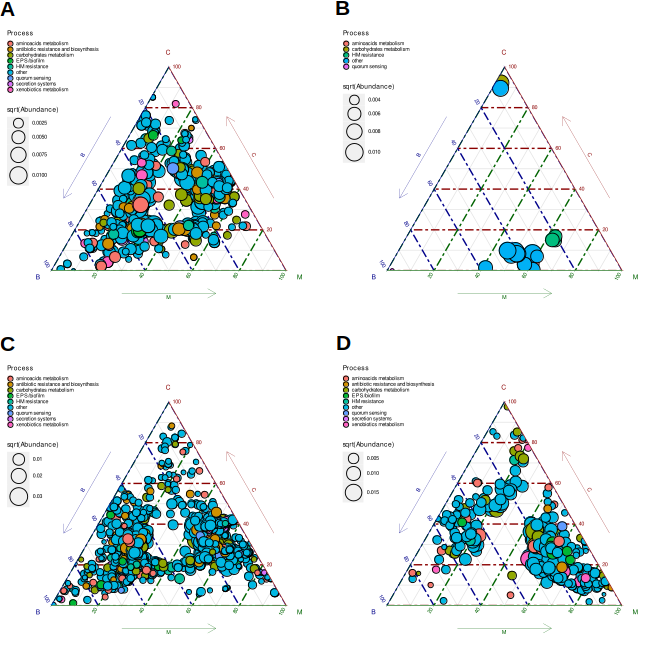
**Fig. SR27.** Sets of KO functions characteristic for roots of different genotypes are different depending on soil and status. Ternary plots of differentially abundant PICRUSt2-predicted KOs identified with DESeq2 are shown. AB – soil S1, CD – soil S2, AC – early samples, BD – late samples. Note differeces between panels’ legends.

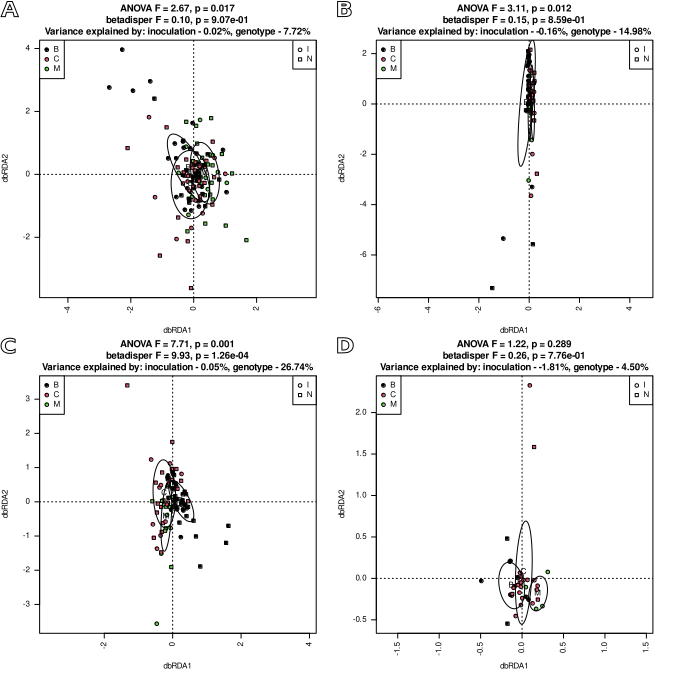
**Fig. SR28**. Sets of KO functions predicted to be encoded by organisms living in leaf samples group according to genotype regardless of soil and status. Early samples (AB), late (CD), soil S1 (AC), S2 (BD). dbRDA models based on Morisita-Horn distance matrices were tested with permutational test (999 permutations). Homogeneity of multi-dimensional variance was tested with betadisper test, and percentage of explained variance was assessed with varpart.

**Fig. SR29.**
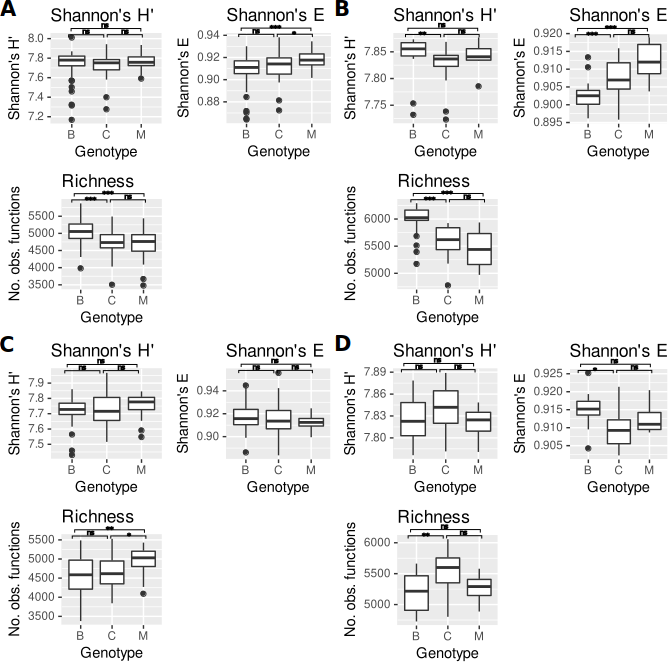
Functional diversity of leaf samples’ bacterial communities depends genotype. AB – soil S1, CD – soil S2, AC – early samples, BD – late samples. Means ± SD of Shannon’s diversity (H’), Shannon’s evennes (E) and observed number of KO functions predicted by PICRUSt2 to be encoded by genomes of organisms thriving in the samples are shown. Significance of difference between genotypes (Kruskal-Wallis test with Dunn’s post-hoc procedure using Benjamini-Hochberg FDR) is shown over the plots: *** - p < 0.001, ** - p < 0.01, * - p < 0.05, ns – not significant.

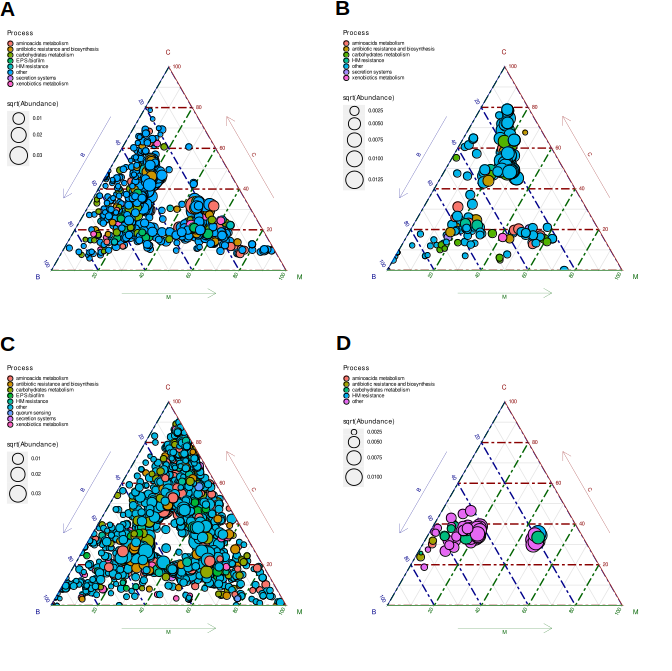
**Fig. SR30.** Sets of KO functions characteristic for leaves of different genotypes are different depending on soil and status. Ternary plots of differentially abundant PICRUSt2-predicted KOs identified with DESeq2 are shown. AB – soil S1, CD – soil S2, AC – early samples, BD – late samples. Note differeces between panels’ legends.

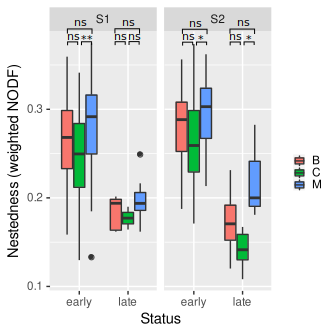

**Fig. SR31.** Nestedness is higher in genotype M than in B and C. Means of weighted NODF index are shown. Significance of difference between genotypes (Kruskal-Wallis test with Dunn’s post-hoc procedure using Benjamini-Hochberg FDR) is shown over the plots: *** - p < 0.001, ** - p < 0.01, * - p < 0.05, ns – not significant.

**Community structure in inoculated and non-inoculated samples differs significantly and the differing ASVs depend on compartment, soil and genotype**

**Table SR7.** Results of permutational tests of dbRDA models based on d05 generalized UniFrac (ASVs) and Morisita-Horn (KOs) distance matrices and involving inoculation.

| **Material** | **Soil** | **Genotype** | **Status** | **Df^1^** | **ASVs** | | **KOs** | |
| --- | --- | --- | --- | --- | --- | --- | --- | --- |
|  |  |  |  |  | **F** | **p** | **F** | **p** |
| S | S1 | B | early | 73 | **2.388** | **0.001** | 1.291 | 0.258 |
|  |  |  | late | 49 | **5.869** | **0.001** | **6.612** | **0.002** |
|  |  | C | early | 65 | **2.292** | **0.007** | **3.764** | **0.010** |
|  |  |  | late | 48 | **1.959** | **0.014** | 1.321 | 0.227 |
|  |  | M | early | 73 | **2.326** | **0.019** | **4.420** | **0.013** |
|  |  |  | late | 58 | **2.873** | **0.001** | 1.743 | 0.175 |
|  | S2 | B | early | 73 | **4.429** | **0.001** | 1.944 | 0.074 |
|  |  |  | late | 58 | **3.463** | **0.001** | **3.217** | **0.009** |
|  |  | C | early | 69 | **11.085** | **0.001** | **8.902** | **0.001** |
|  |  |  | late | 57 | **5.106** | **0.001** | 1.891 | 0.113 |
|  |  | M | early | 72 | **4.532** | **0.001** | **4.230** | **0.011** |
|  |  |  | late | 58 | **1.912** | **0.049** | 0.509 | 0.584 |
| R | S1 | B | early | 46 | **1.606** | **0.040** | 1.215 | 0.299 |
|  |  |  | late | - | - | - | - | - |
|  |  | C | early | 50 | 1.384 | 0.058 | 1.820 | 0.116 |
|  |  |  | late | - | - | - | - | - |
|  |  | M | early | 68 | 1.031 | 0.410 | 0.639 | 0.602 |
|  |  |  | late | 21 | 1.040 | 0.383 | 1.952 | 0.138 |
|  | S2 | B | early | 36 | 1.396 | 0.077 | 0.944 | 0.435 |
|  |  |  | late | 6 | **2.519** | **0.018** | 0.625 | 0.609 |
|  |  | C | early | 56 | **3.319** | **0.001** | 1.166 | 0.283 |
|  |  |  | late | 7 | 1.441 | 0.161 | 3.099 | 0.086 |
|  |  | M | early | 66 | 1.289 | 0.122 | 0.687 | 0.561 |
|  |  |  | late | 22 | 0.939 | 0.541 | 1.967 | 0.137 |
| L | S1 | B | early | 52 | **2.355** | **0.001** | 1.505 | 0.236 |
|  |  |  | late | 24 | 0.872 | 0.884 | 0.804 | 0.451 |
|  |  | C | early | 36 | 1.057 | 0.373 | 0.207 | 0.935 |
|  |  |  | late | 17 | 0.995 | 0.447 | 2.232 | 0.160 |
|  |  | M | early | 34 | **1.733** | **0.013** | 0.578 | 0.695 |
|  |  |  | late | 8 | 1.091 | 0.138 | 1.976 | 0.114 |
|  | S2 | B | early | 42 | 1.492 | 0.069 | 0.901 | 0.402 |
|  |  |  | late | 8 | 0.877 | 0.912 | 0.630 | 0.698 |
|  |  | C | early | 29 | **1.620** | **0.033** | 0.578 | 0.743 |
|  |  |  | late | 21 | 1.199 | 0.099 | 0.740 | 0.504 |
|  |  | M | early | 17 | **1.579** | **0.026** | 1.294 | 0.271 |
|  |  |  | late | - |  |  |  |  |

1 – degrees of freedom in the denominator of the pseudo-F distribution used by anova.cca

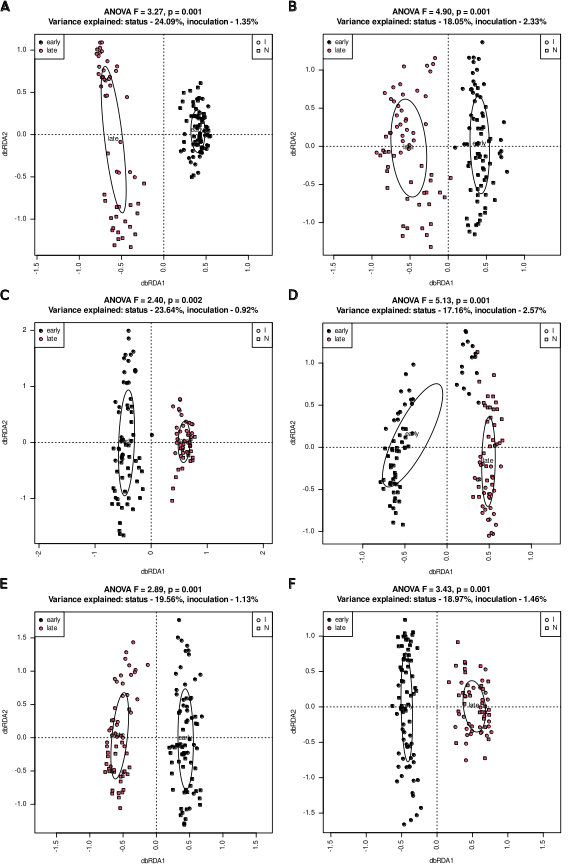

**Fig. SR32**. Bacterial communities in inoculated (squares) and non-inoculated (circles) soil samples differ regardless of an experimental variant and status. dbRDA models based on d05 generalized UniFrac distance matrices and involving inoculation and status are shown for soil S1 (ACE) and S2 (BDF) and genotypes B (AB), C (CD) and M (EF). P-value of a dbRDA model involving inoculation and with status conditioned out is given

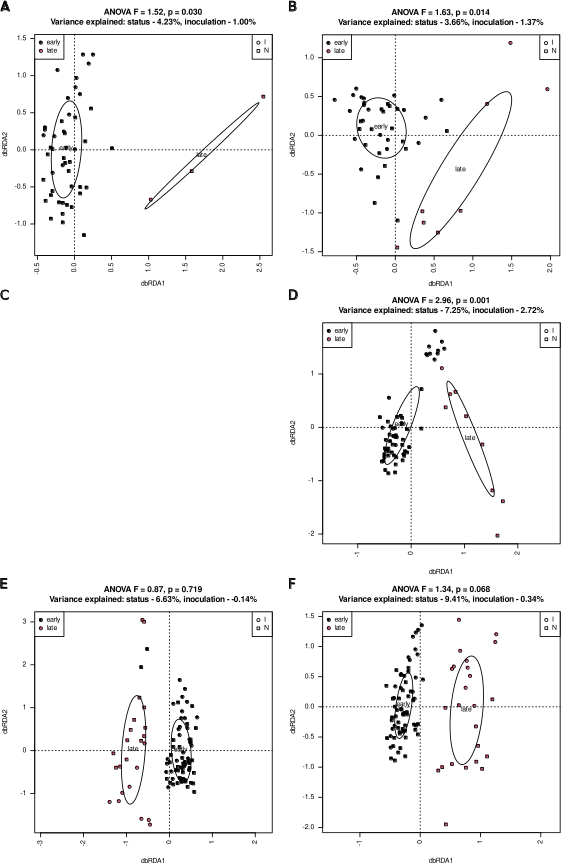

**Fig. SR33**. Bacterial communities in inoculated (squares) and non-inoculated (circles) root samples differ more in late than in early ones regardless of an experimental variant. dbRDA models based on d05 generalized UniFrac distance matrices are shown for soil S1 (ACE) and S2 (BDF) and genotypes B (AB), C (CD) and M (EF). P-value of a dbRDA model involving inoculation and with status conditioned out is given

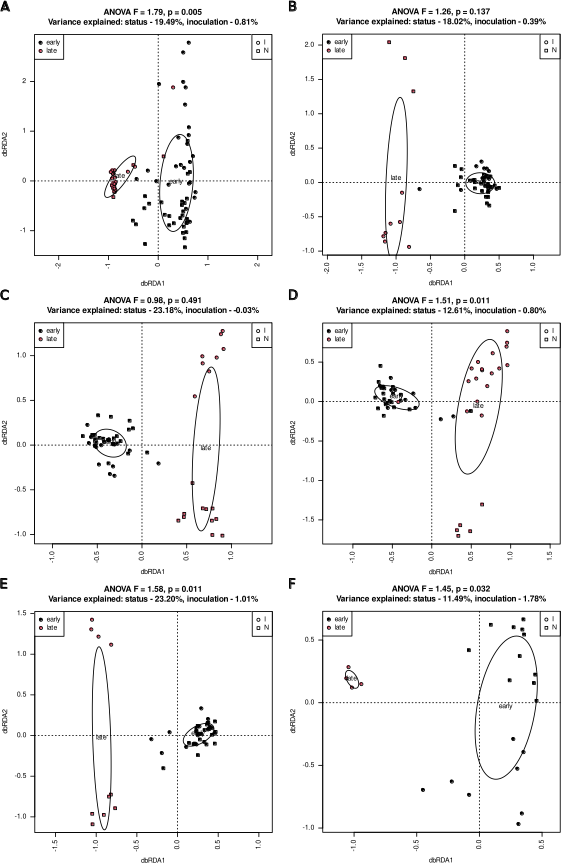

**Fig. SR34**. Bacterial communities in inoculated (squares) and non-inoculated (circles) late but not early leaf samples differ regardless of an experimental variant. dbRDA models based on d05 generalized UniFrac distance matrices are shown for soil S1 (ACE) and S2 (BDF) and genotypes B (AB), C (CD) and M (EF). P-value of a dbRDA model involving inoculation and with status conditioned out is given

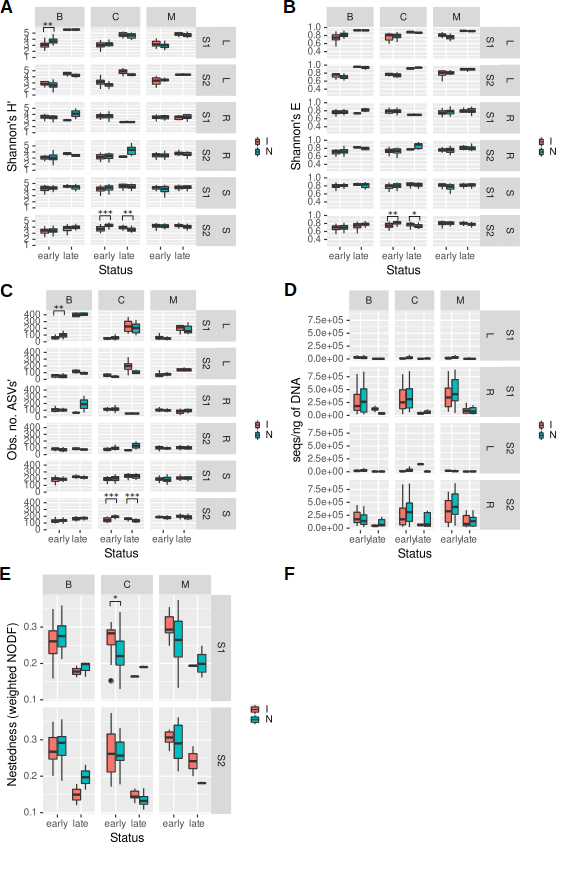

**Fig. SR35.** Alpha-diversity measures, bacterial load and nestedness are not influenced by inoculation regardless of an experimental variant. A – Shannon’s diversity index (H’), B – Shannon’s evenness (E), C – observed number of ASVs, D – bacterial load measured as number of 16S rRNA gene fragments per ng of DNA, E – nestedness measured as weighted NODF index. Statistical significance was assessed with Wilcoxon test and p-value was corrected for multiple comparisons using Benjamini-Hochberg FDR. Significant p-values are shown over the plots: *** - p < 0.001, ** - p < 0.01, * - p < 0.05.

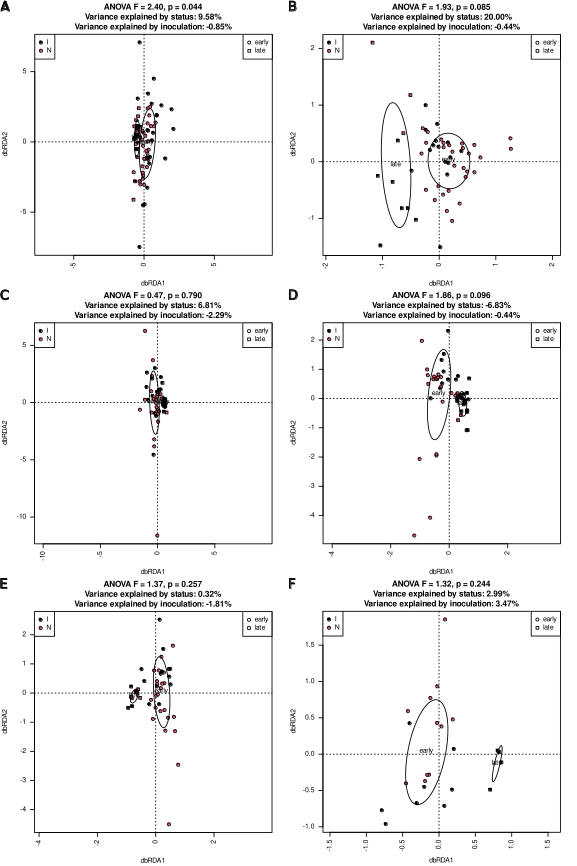

**Fig. SR36**. Functional potential of bacterial communities in inoculated and non-inoculated leaf samples do not differ significantly. dbRDA models based on Morisita-Horn distance matrices and involving status and inoculation are shown for soil S1 (ACE) and S2 (BDF) and genotypes B (AB), C (CD) and M (EF). Negative values of variance explained should be interpreted as zero. P-value of a dbRDA model involving inoculation and with status conditioned out is given

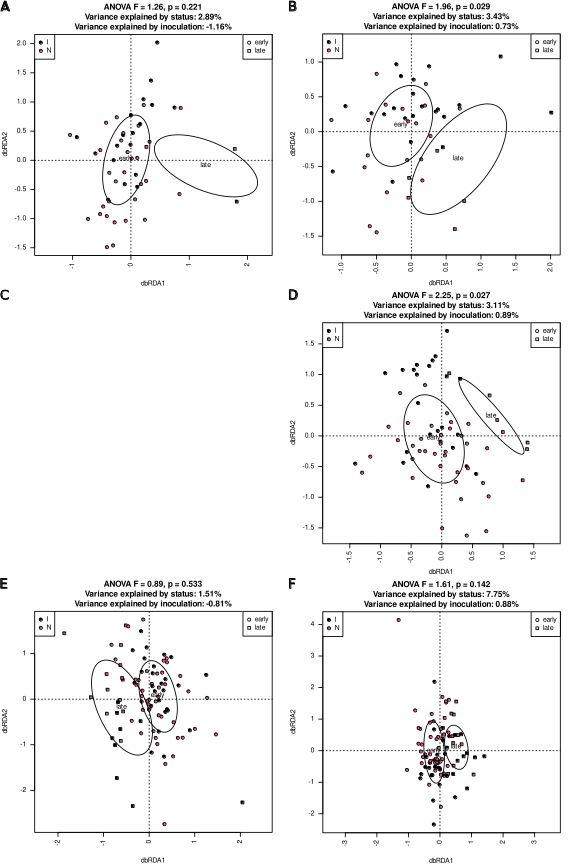

**Fig. SR37**. Functional potential of bacterial communities in inoculated and non-inoculated root samples does not differ significantly. dbRDA models based on Morisita-Horn distance matrices and involving status and inoculation are shown for soil S1 (ACE) and S2 (BDF) and genotypes B (AB), C (CD) and M (EF). Negative values of variance explained should be interpreted as zero. P-value of a dbRDA model involving inoculation and with status conditioned out is given.

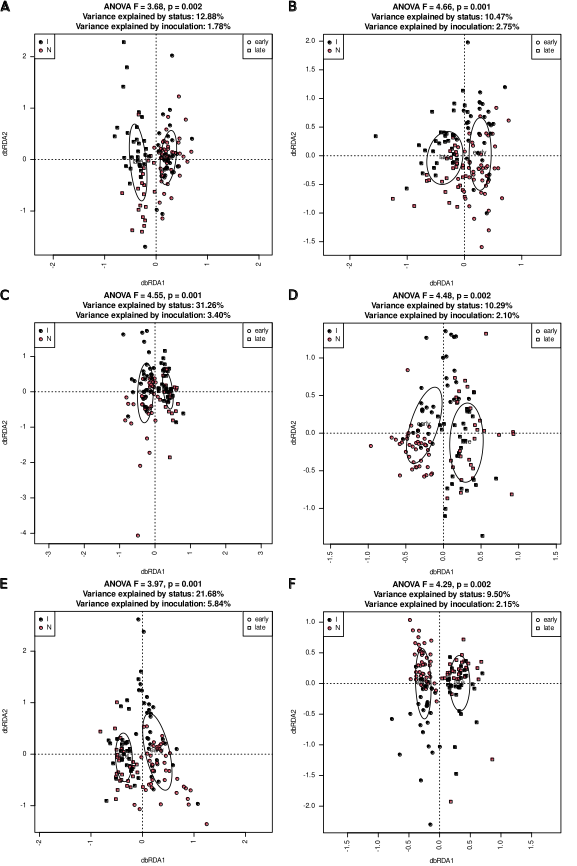

**Fig. SR38**. Functional potential of bacterial communities in inoculated and non-inoculated soil samples does not differ significantly. dbRDA models based on Morisita-Horn distance matrices and involving status and inoculation are shown for soil S1 (ACE) and S2 (BDF) and genotypes B (AB), C (CD) and M (EF). Negative values of variance explained should be interpreted as zero. P-value of a dbRDA model involving inoculation and with status conditioned out is given

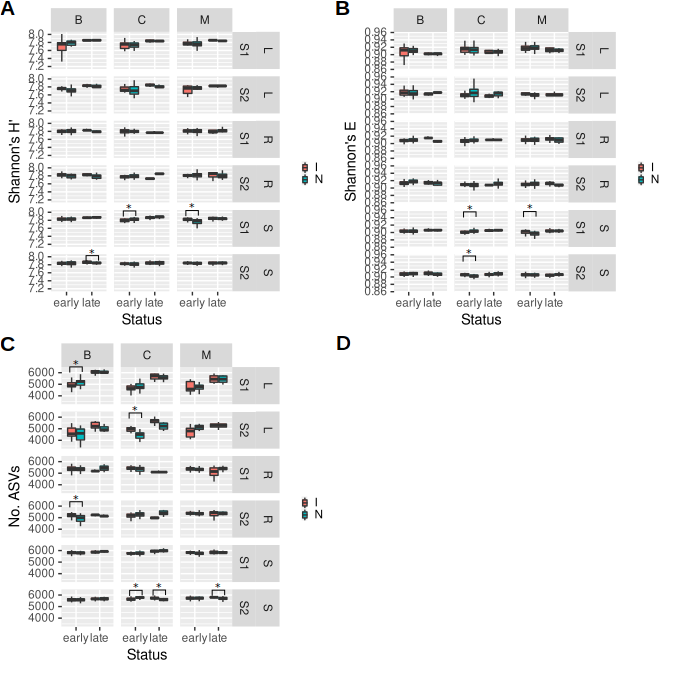

**Fig. SR39.**

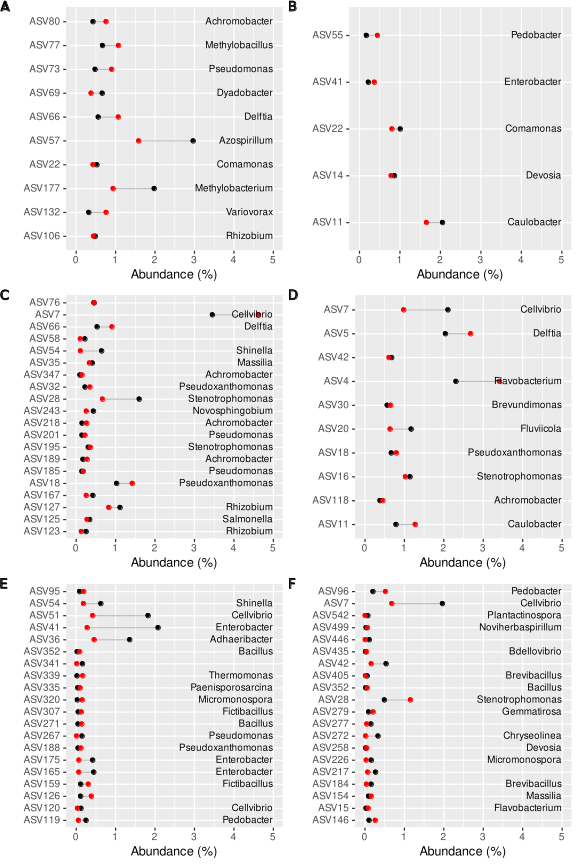

**Fig. SR40.** ASVs differentially represented in inoculated and non-inoculated early (ACE) and late (BDF) leaf (AB), root (CD) and soil (EF) samples. Red circles represent non-inoculated samples, black circles – inoculated ones. At most 24 features of greatest fold-change are shown.

Complete sets of differentially represented ASVs and higher taxa can be found in SupplementaryResultsF1, and sets of differentially represented KO functions are in SupplementaryResultsF2.

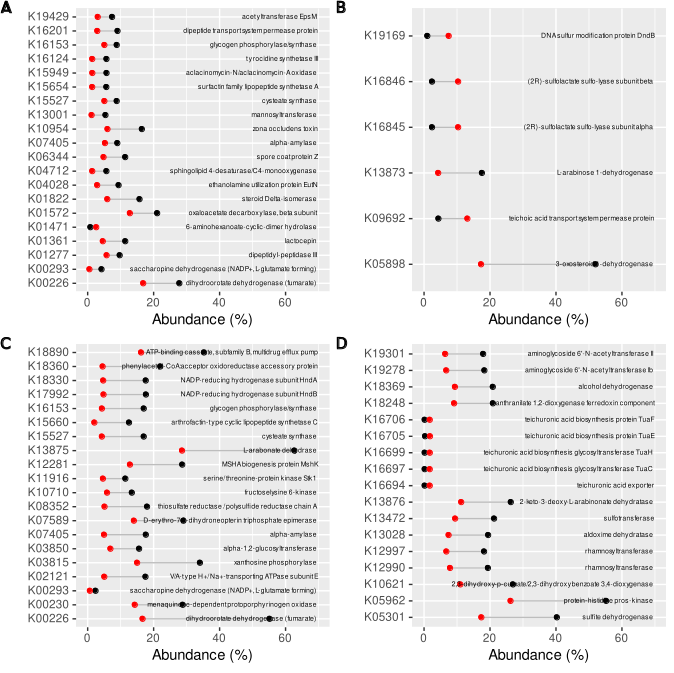
**Fig. SR41**. Functions differentially represented in inoculated and non-inoculated early (AC) and late (BD) root (AB) and soil (CD) samples. Red circles represent non-inoculated samples, black circles – inoculated ones. At most 24 features of greatest fold-change are shown. There were no differentially represented functions in leaf samples.

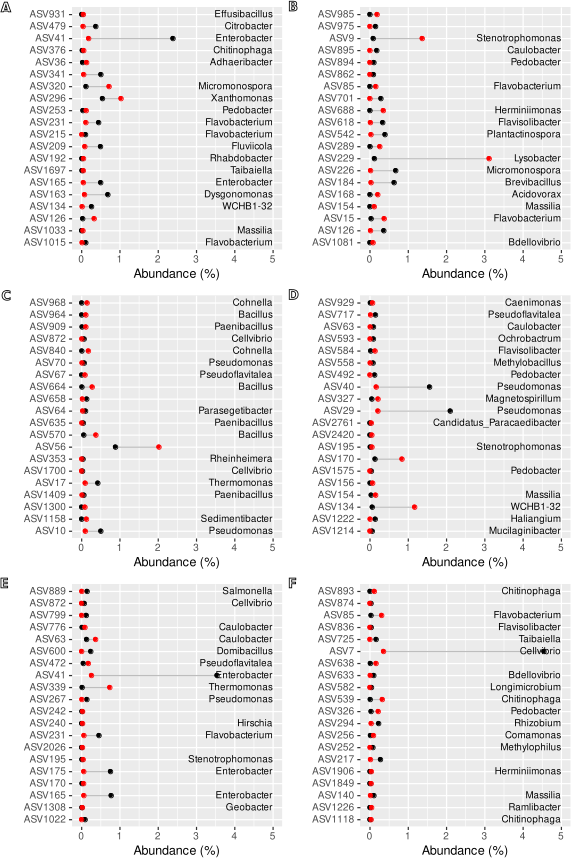

**Fig. SR42**. ASVs differentially represented in inoculated and non-inoculated S1 soil samples. Early samples (ACE) and late ones (BDF), genotype B (AB), C (CD) and M (EF). At most 10 ASVs with greatest and lowest fold change are shown Red circles represent non-inoculated samples, black circles – inoculated ones. Differential abundance analysis was carried out using DESeq2.

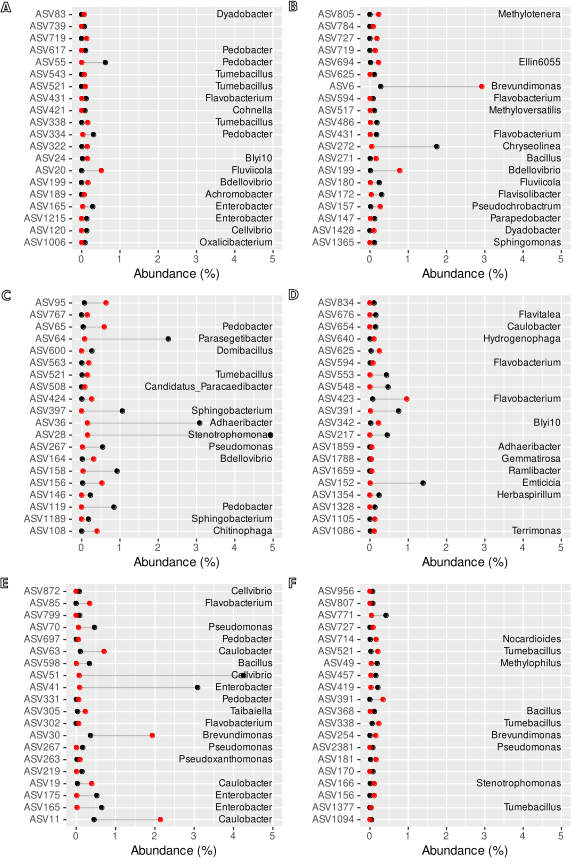

**Fig. SR43**. ASVs differentially represented in inoculated and non-inoculated S2 soil samples. Early samples (ACE) and late ones (BDF), genotype B (AB), C (CD) and M (EF). At most 10 ASVs with greatest and lowest fold change are shown. Red circles represent non-inoculated samples, black circles – inoculated ones. Differential abundance analysis was carried out using DESeq2.

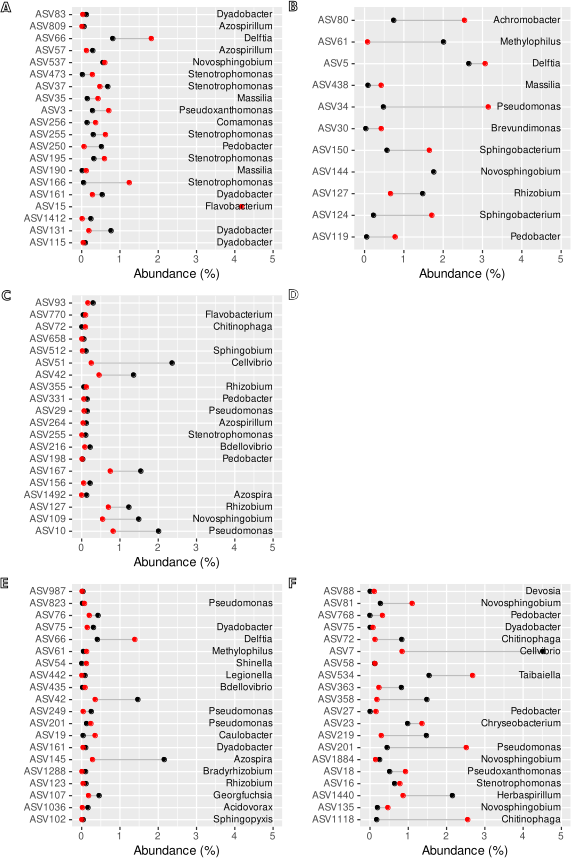

**Fig. SR44**. ASVs differentially represented in inoculated and non-inoculated S1 root samples. Early samples (ACE) and late ones (BDF), genotype B (AB), C (CD) and M (EF). At most 10 ASVs with greatest and lowest fold change are shown. Red circles represent non-inoculated samples, black circles – inoculated ones. Differential abundance analysis was carried out using DESeq2.

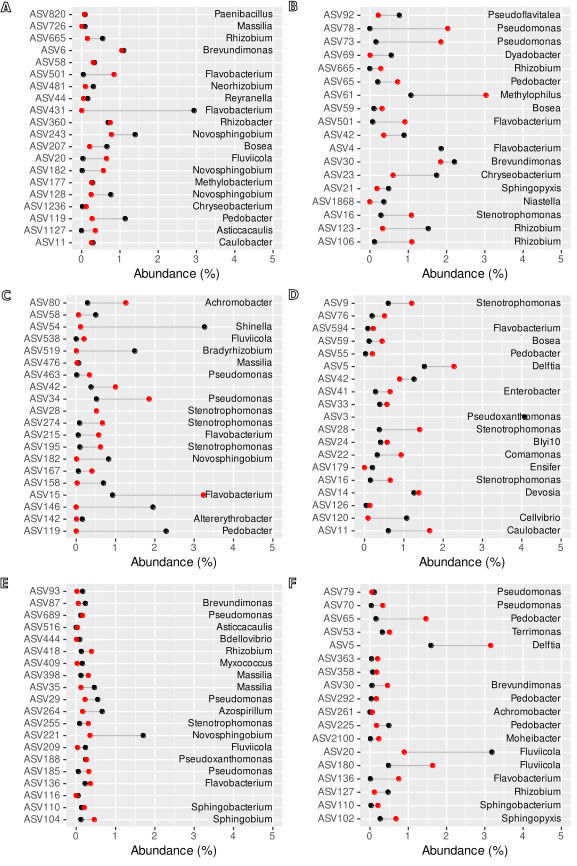

**Fig. SR45**. ASVs differentially represented in inoculated and non-inoculated S2 root samples. Early samples (ACE) and late ones (BDF), genotype B (AB), C (CD) and M (EF). At most 10 ASVs with greatest and lowest fold change are shown. Red circles represent non-inoculated samples, black circles – inoculated ones. Differential abundance analysis was carried out using DESeq2.

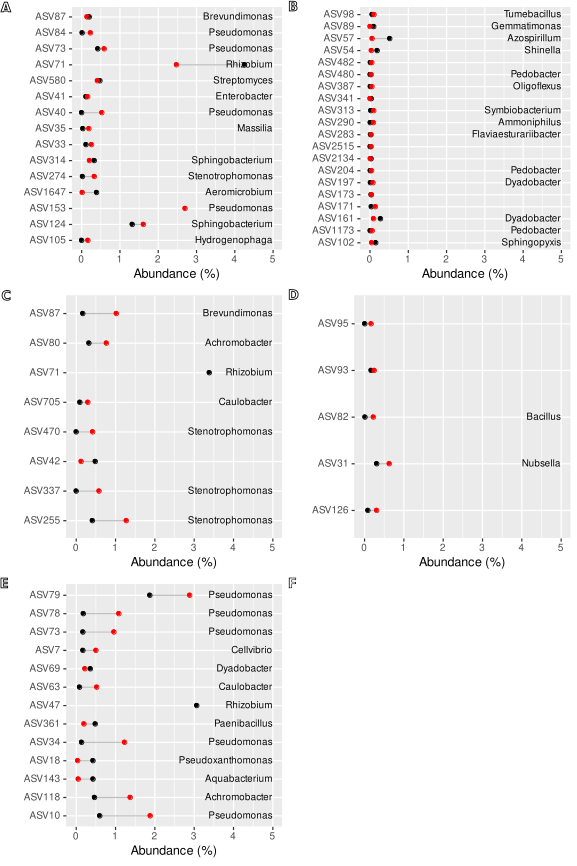

**Fig. SR46**. ASVs differentially represented in inoculated and non-inoculated S1 leaf samples. Early samples (ACE) and late ones (BDF), genotype B (AB), C (CD) and M (EF). At most 10 ASVs with greatest and lowest fold change are shown. Red circles represent non-inoculated samples, black circles – inoculated ones. Differential abundance analysis was carried out using DESeq2.

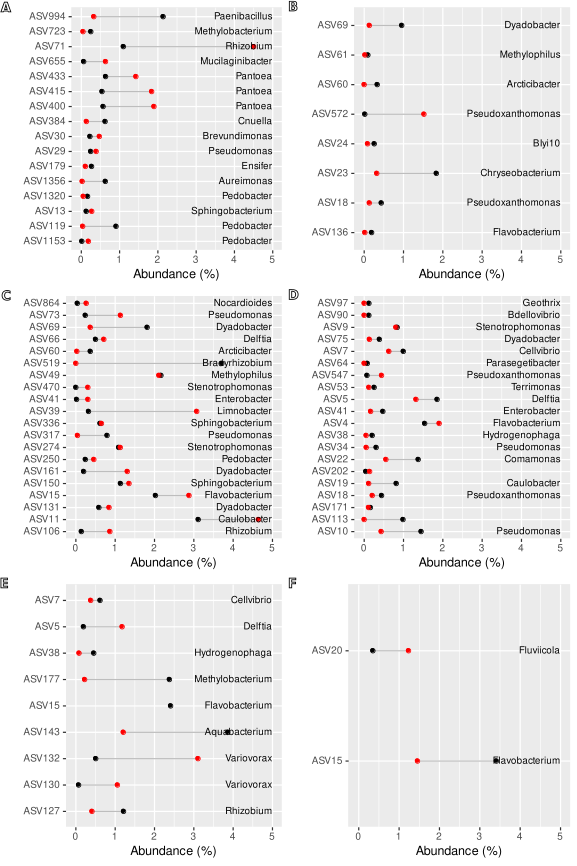

**Fig. SR47**. ASVs differentially represented in inoculated and non-inoculated S2 leaf samples. Early samples (ACE) and late ones (BDF), genotype B (AB), C (CD) and M (EF). At most 10 ASVs with greatest and lowest fold change are shown. Red circles represent non-inoculated samples, black circles – inoculated ones. Differential abundance analysis was carried out using DESeq2.

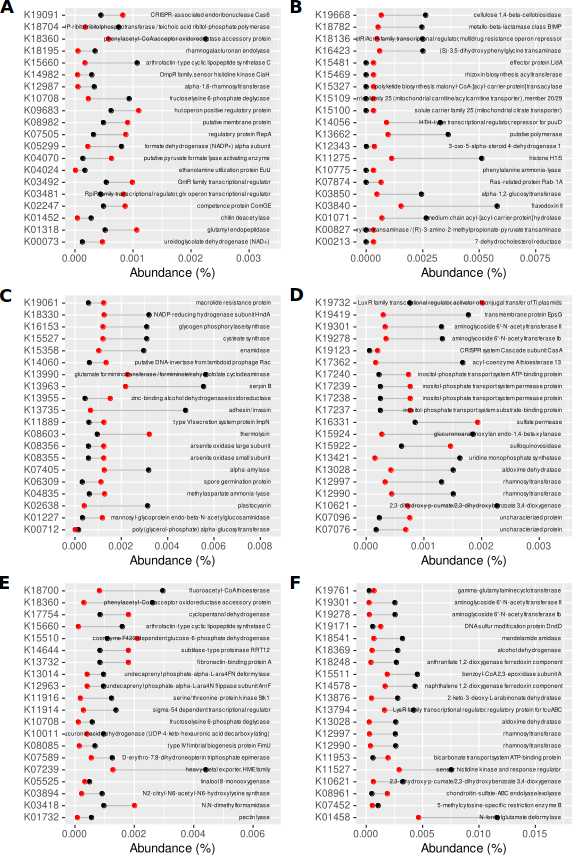

**Fig. SR48**. KO functions differentially represented in genomes of organisms thriving in inoculated and non-inoculated S1 soil samples. Early samples (ACE) and late ones (BDF), genotype B (AB), C (CD) and M (EF). At most 10 functions with greatest and lowest fold change are shown. Red circles represent non-inoculated samples, black circles – inoculated ones. Differential abundance analysis was carried out using DESeq2.

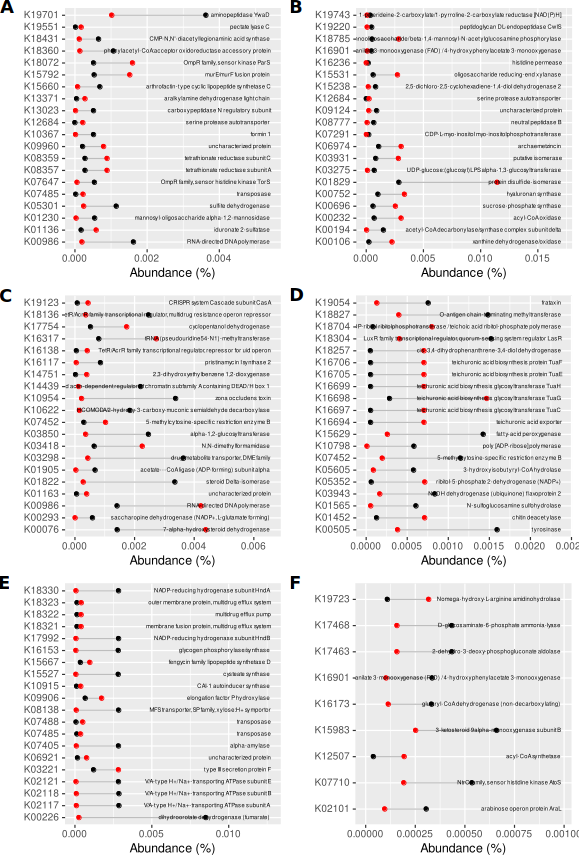

**Fig. SR49**. KO functions differentially represented in genomes of organisms thriving in inoculated and non-inoculated S2 soil samples. Early samples (ACE) and late ones (BDF), genotype B (AB), C (CD) and M (EF). At most 10 functions with greatest and lowest fold change are shown. Red circles represent non-inoculated samples, black circles – inoculated ones. Differential abundance analysis was carried out using DESeq2.

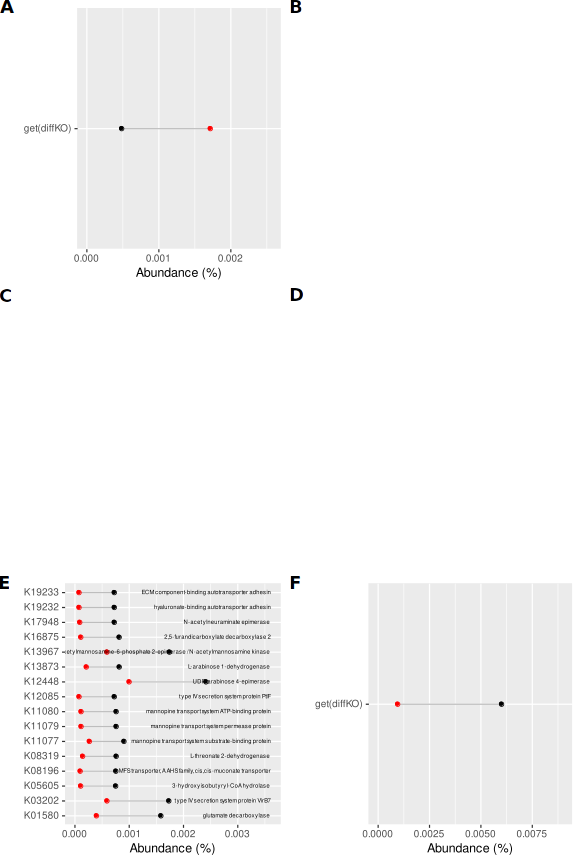

**Fig. SR50**. KO functions differentially represented in genomes of organisms thriving in inoculated and non-inoculated S1 root samples. Early samples (ACE) and late ones (BDF), genotype B (AB), C (CD) and M (EF). At most 10 functions with greatest and lowest fold change are shown. Red circles represent non-inoculated samples, black circles – inoculated ones. Differential abundance analysis was carried out using DESeq2. The lack of a graph means that there were no differentially represented functions.

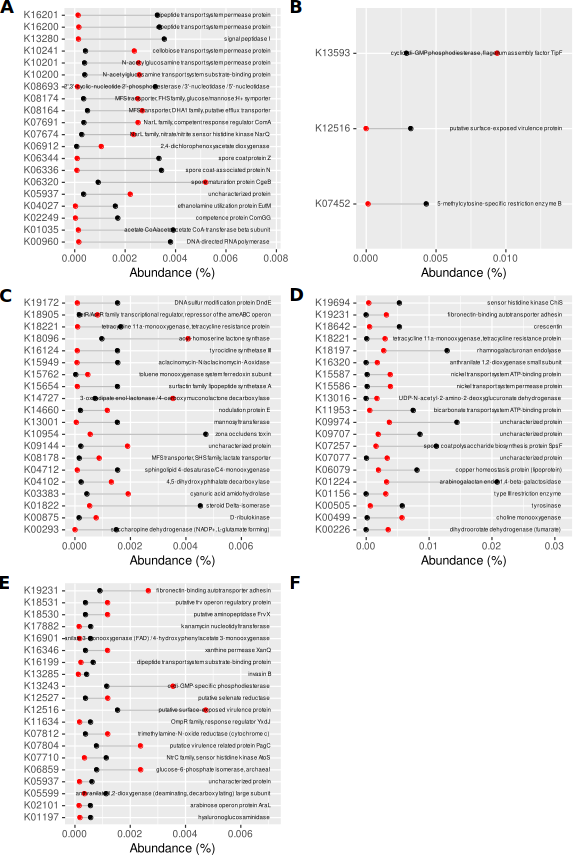

**Fig. SR51**. KO functions differentially represented in genomes of organisms thriving in inoculated and non-inoculated S2 root samples. Early samples (ACE) and late ones (BDF), genotype B (AB), C (CD) and M (EF). At most 10 functions with greatest and lowest fold change are shown. Red circles represent non-inoculated samples, black circles – inoculated ones. Differential abundance analysis was carried out using DESeq2. The lack of a graph means that there were no differentially represented functions.

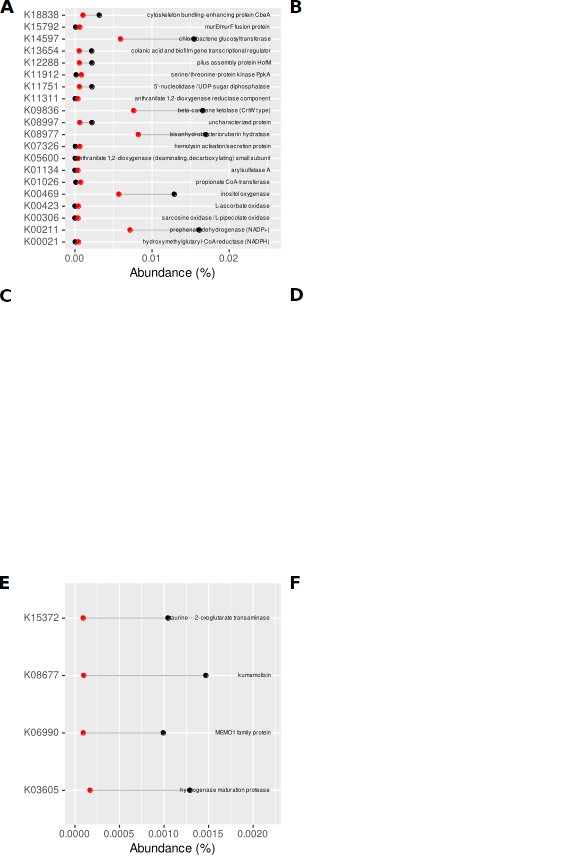

**Fig. SR52**. KO functions differentially represented in genomes of organisms thriving in inoculated and non-inoculated S1 leaf samples. Early samples (ACE) and late ones (BDF), genotype B (AB), C (CD) and M (EF). At most 10 functions with greatest and 10 with lowest fold change are shown. Red circles represent non-inoculated samples, black circles – inoculated ones. Differential abundance analysis was carried out using DESeq2. The lack of a graph means that there were no differentially represented functions.

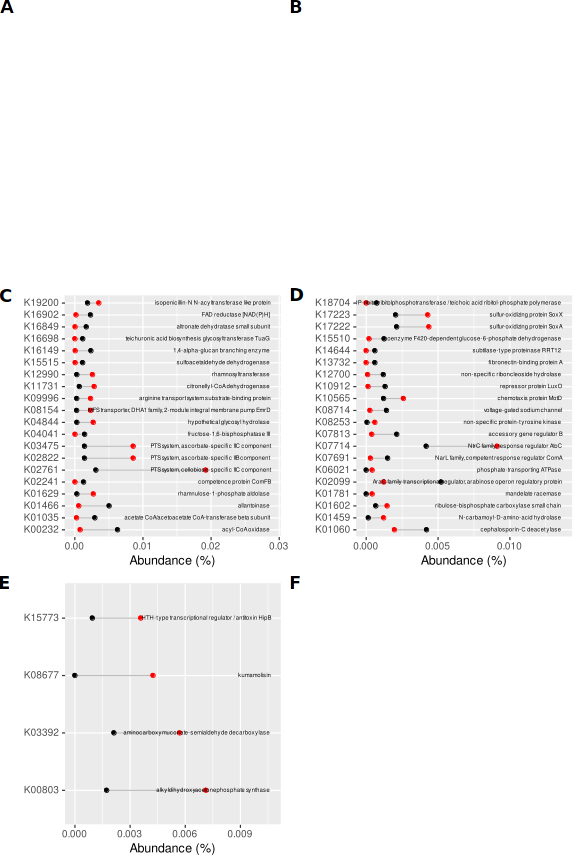

**Fig. SR53**. KO functions differentially represented in genomes of organisms thriving in inoculated and non-inoculated S2 leaf samples. Early samples (ACE) and late ones (BDF), genotype B (AB), C (CD) and M (EF). At most 20 functions with greatest and lowest fold change are shown. Red circles represent non-inoculated samples, black circles – inoculated ones. Differential abundance analysis was carried out using DESeq2. The lack of a graph means that there were no differentially represented functions.

**Table SR8. ASVs constituting signature of inoculated samples identified with biosigner.**

| **ASV** | **Taxonomy** |
| --- | --- |
| ASV3 | Pseudoxanthomonas |
| ASV4 | Flavobacterium |
| ASV5 | Delftia |
| ASV6 | Brevundimonas |
| ASV7 | Cellvibrio |
| ASV10 | Pseudomonas |
| ASV11 | Caulobacter |
| ASV14 | Devosia |
| ASV16 | Stenotrophomonas |
| ASV20 | Fluviicola |
| ASV31 | Nubsella |
| ASV41 | Enterobacter |
| ASV66 | Delftia |
| ASV70 | Pseudomonas |
| ASV77 | Methylobacillus |
| ASV119 | Pedobacter |
| ASV159 | Fictibacillus |
| ASV165 | Enterobacter |
| ASV175 | Enterobacter |
| ASV217 | Memeber of Sphigobacteriaceae |
| ASV273 | Tumebacillus |
| ASV307 | Fictibacillus |
| ASV542 | Plantactinospora |
| ASV603 | Fictibacillus |
| ASV617 | Pedobacter |
| ASV656 | Galbitalea |
| ASV799 | Member of Enterobacteriaceae |
| ASV872 | Cellvibrio |

**Table SR9. ASVs present in inoculant and inoculated samples, but absent from non-inoculated ones.**

| **ASV** | **Taxonomy** |
| --- | --- |
| ASV1254 | Cellvibrio |
| ASV2212 | Devosia |
| ASV2539 | Member of Moraxellaceae |
| ASV3930 | Brevibacillus |
| ASV4420 | Nocardioides |
| ASV4847 | Brevundimonas |
| ASV4848 | Pedobacter |
| ASV4949 | Member of Paenibacillaceae |
| ASV5743 | Member of Paenibacillaceae |
| ASV5988 | Member of Burkholderiaceae |
| ASV7103 | Algoriphagus |
| ASV8922 | Terrimonas |
| ASV9530 | Member of Gammproteobacterial order PLTA13 |
| ASV11034 | Member of Chitinophagaceae |
| ASV12901 | Herbinix |
